# The genomic context for aflatoxin B1-degrading *Pseudomonas* strains

**DOI:** 10.1101/2021.09.24.461607

**Authors:** Jiahong Tang, Dun Deng, Maopeng Song, Zhichang Liu, Sanmei Ma, Yongfei Wang

## Abstract

The whole genomes of three strains were sequenced and annotated. COG (Clusters of Orthologous Groups) and GO (Gene Ontology) annotations of the protein-coding genes from three strains show a conservation of genome-wide protein functions in genus Pseudomonas. However, the AFB1-degrading strains HAI2 and HT3 harbor much more genes belonged to the pathway of xenobiotics biodegradation and metabolism than non-degrading strain 48. Besides, the enzyme families potentially involved in the AFB1 degradation of bacteria are more abundant in the two AFB1-degrading strains. A pan-genome profile was then formed by comparing the genomes against other reference genomes of the corresponding Pseudomonas species. Accordingly, a total of 1,528 genes were found to be specific in AFB1-degrading strains, and 65 genes of them are related to oxidoreductase activity.

## Introduction

Aflatoxins (AF) are a class of secondary metabolites with highly toxic and carcinogenic properties produced by fungi such as *Aspergillus flavus* and *Aspergillus parasiticu*, including aflatoxins B1, B2, G1, G2, M, and M2, among of them, aflatoxin B1 (AFB1) is the most toxic. AFB1 is one of the most toxic mycotoxins found at present, and has carcinogenic and carcinogenic effects. Teratogenic and mutagenic. In 1993, the International Agency for Research on Cancer (IARC) of the World Health Organization classified aflatoxin as a class I carcinogen. Aflatoxin B1 can be transmitted to humans and animals through the food chain. However, the contamination of *Aspergillus flavu*s in peanuts, corn and other food crops and feed ingredients is very common. Therefore, how to prevent and control such toxicity is an urgent problem that needs to be solved.

Biological detoxification of AFB1 is a major toxin removal method, which mainly uses microorganisms or enzymes produced by them and their preparations to detoxify through biocatalysis. Compared with physical and chemical methods, biodegradation method is mild, AFB1 is completely degraded, and toxic residue is weak, so it is favored by researchers. Microorganisms capable of degrading AFB1 are various, and many fungi and bacteria have the ability to degrade AFB1. For example, in bacteria *Bacillus licheniformis*, *Pontibacter sp.* (VGF1), *Pseudomonas aeruginosa*, *Rhodococcus strains*, *Staphylocococcus warneri*, and *Tetragenococcus halophilus*. And some fungi such as *Armillariella tabescens* and *Candida versatilis*, etc., could efficient degrade AFB1. These microorganisms provide a premise for solving the aflatoxin-pollution problem. Understanding the mechanism of microbial degradation of AFB1 is helpful for designing toxin detoxification preparations, but few of these microorganisms are responsible for the degradation of AFB1.

Studies have confirmed that some enzymes have the function of degrading AFB1. MnP (Manganese peroxidase) produced by white rot fungus (*Phanerochaete sordida* YK-624) which can degrade lignin has the function of AFB1 degradation, AFB1 oxidase in *Armillaria*, F420H2 dependent reductases in Mycobacterium and laccase in some microorganisms could be also catalyze AFB1 reactions. These enzymological information provide a necessary basis for us to develop enzyme preparations for degrading AFB1. In the present study, we obtained three *Pseudomonas* strains with completely different degradation performance of AFB1. In this study, whole genome sequencing and comparative genomics was used to construct a genomic architecture of ABF1-degrading *Pseudomonas* strains. Based on functional annotations, potential genes related to AFB1 degradation were excavated, which laid a foundation for the development of corresponding enzyme preparations.

## Materials and Methods

### *Isolation of Pseudomonas* strains

*Pseudomonas* strain HAI-2 was isolated from the mangrove sludge sample, HT-3 was isolated from the soil near the feed warehouse, and 48 was stored in the laboratory. 1g sludge or soil sample was added 9 mL of 0.9 % saline, shaked and performed gradient dilution, then cultured on coumarin medium as previously described (Samuel, M.S et al., 2014). After 72h, HAI2 and HT3 could grow on CM medium after cultured 72h, and single colonies were further tested for AFB_1_-degrading activity and whole-genome analyses.

### Measurement of AFB1 degradation

*Pseudomo*nas strain HAI-2, HT3 and 48 were cultured in LB liquid medium at 200rpm/min, 37°C. After cultured for 24h, the bacteria cells were collected by centrifugation (12000rpm/min, 4 °C) and adjust the bacteria density to 10^9^cfu / mL using a carbon-free medium to preform AFB1 degradation. The reaction was carried out in a 20 mL steriled glass bottle. 1 mL of AFB1 degradation reaction system contained 1000ng/mL AFB_1_ at 30°C in the dark under static conditions. After 24 and 48 h of reaction, the AFB_1_ content was analyzed by HPLC, and E. coli DH5α at the same concentration were served as control.

The reaction mixtures were stoped by addition equal volume chloroform, then residual AFB_1_ were extracted three times with chloroform according to a standard protocol (Yang et al., 2014). The chloroform in the upper layer was incorporated and evaporated under nitrogen gas at 60 °C and dried extracts were dissolved in 50% methanol. After filtered by 0.22-μm pore size filter, the concentration of AFB_1_ analyzed by HPLC equipped with a fluorescence detector (HPLC–FLD). An XBridgeTM C18 (4.6 mm×250 mm, 5 μm) liquid chromatography (LC) column was applied with a mobile phase containing water/acetonitrile/methanol (35: 55: 10, V/V) as the mobile phase at a flow rate of 1 mL/min, a column temperature of 37°C, and an injection volume of 20 μL. AFB1 was derived by a photochemical reactor (Waters, Milford, MA, USA) and detected with a fluorescence detector, using 375 and 430 nm as the wavelengths of excitation and emission, respectively.

### DNA extraction and genome sequencing

Genomic DNA was extracted from *Pseudomonas* strains by DNA extraction and purification with the HiPure Bacterial DNA Kit (Magen, Guangzhou, China) according to the instruction. Each DNA sample was then fragmented into 400-bp fragments by a Covaris M200 sonicator, and the sequencing libraries were generated with NEXTflex Rapid DNA-Seq plus Kit (BIOO, USA). The whole genome sequencing was performed on the Illumina Nextseq550 platform with an average coverage of more than 100×.

### Genome assembly and annotation

The sequencing raw data was filtered with software Trimmomatic (Bolger et al., 2014). Clean reads were then used for *de novo* assembly with software SPAdes v3.6.2 (Bankevich et al., 2012). Gene annotation for each assembled sequence was conducted through software Prokka v1.11 (Seemann, 2014). The protein-coding genes of all strains were further annotated in other public databases including NR (Non-redundant), COG (Clusters of Orthologous Groups), GO (Gene Ontology), and KEGG (Kyoto Encyclopedia of Genes and Genomes).

### Pan-genome analysis

For pan-genome analysis, we added several reference genomic sequences from three *Pseudomonas* species (*P. aeruginosa*, *P. putida*, and *P. stutzeri*) that correspond to the tested species in our study (Table S1). This dataset also included a *P. aeruginosa* strain N17-1, whose ability of AFB1 degradation has been proven (Sangare et al., 2014). The output of Prokka was used as input for the pan-genome pipeline Roary v3.11.2 (Page et al., 2015), with a BLASTP identity cutoff of 70%. We then used a local script to transfer the profile of absence or presence of all genes across all samples into a 0/1 matrix. According to the gene presence and absence matrix, we screened for genes that are specific to AFB1-degrading *Pseudomonas* strains.

### Data Accessibility Statement

The draft genome sequences generated in this study were submitted to the NCBI database under BioProject PRJNA694503.

## Results

### Isolation and identification of AFB1-degrading *Pseudomonas* strains

Three *Pseudomonas* strains were previously obtained from different sources, and a 16S rRNA-based phylogenetic analysis with reference *Pseudomonas* strains in the GenBank database indicated that they were belonged to species *P. putida* (HAI2), *P. aeruginosa* (HT3), and *P. stutzeri* (48), respectively (Figure 1). The degradation efficiency of AFB1 by different strains is shown in Figure 2, it was found that the degradation rates of AFB1 by the three strains of Pseudomonas were quite different. Strain 48 has almost no degradation of AFB1, but HAI-2 and HT-3 can both degrade AFB1 at 1,000ng / mL. In particular, the degradation rate of HT-3 at 24 h and 48 h reached 83.6% and 95.3%, respectively. Strain HT-3 have the highest degradation efficiency to AFB1 among three Pseudomonas strains.

**Figure 1.**
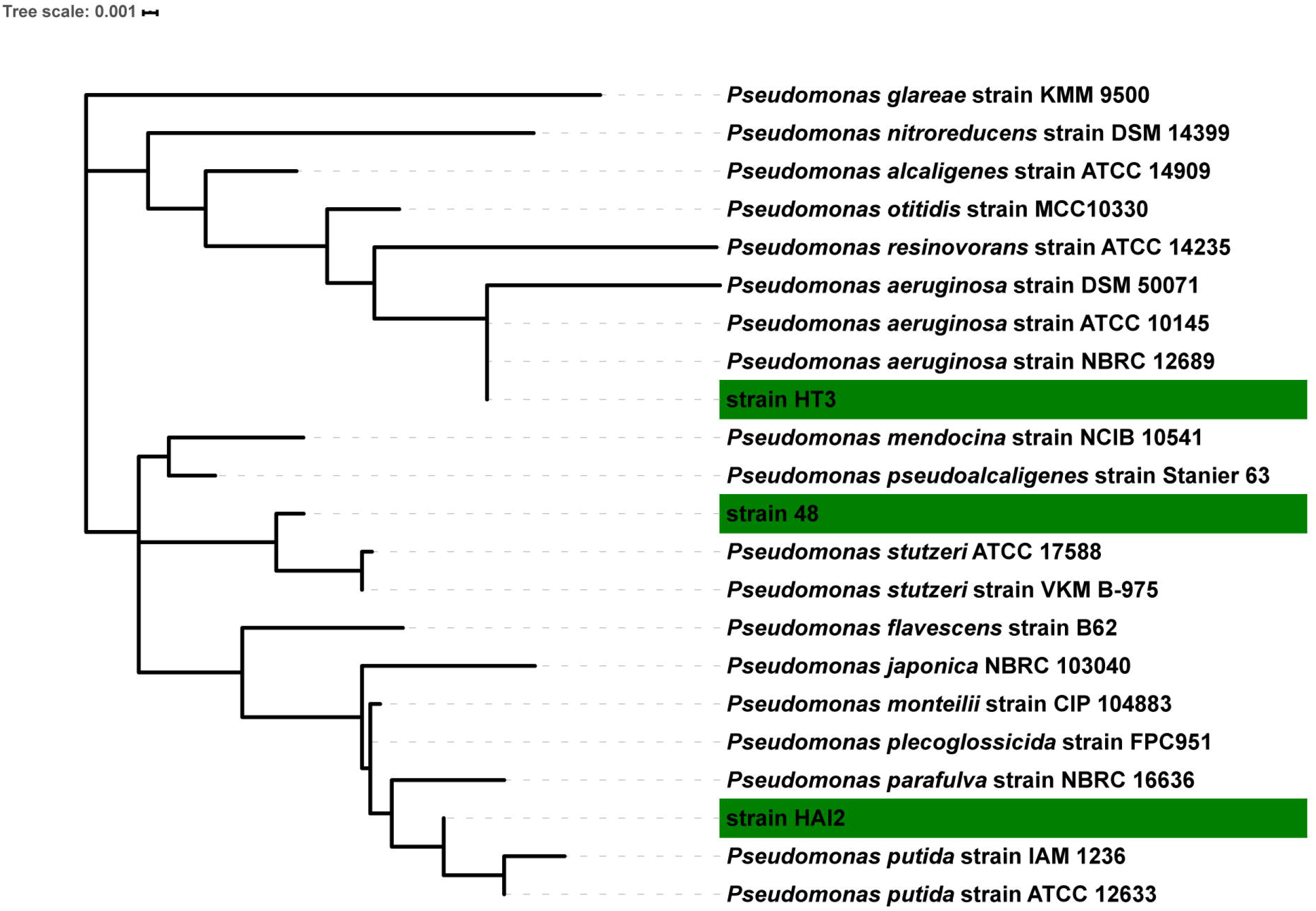
Phylogenetic tree based on the 16S rRNA sequences of *Pseudomonas* strains. The maximum likelihood tree was calculated by MEGA 7.0 with a GTR+G+I model estimated by jModeltest v2.1.7 and 1,000 iterations of bootstrap. The bar indicates a genetic distance of 0.001.

**Figure 2.**
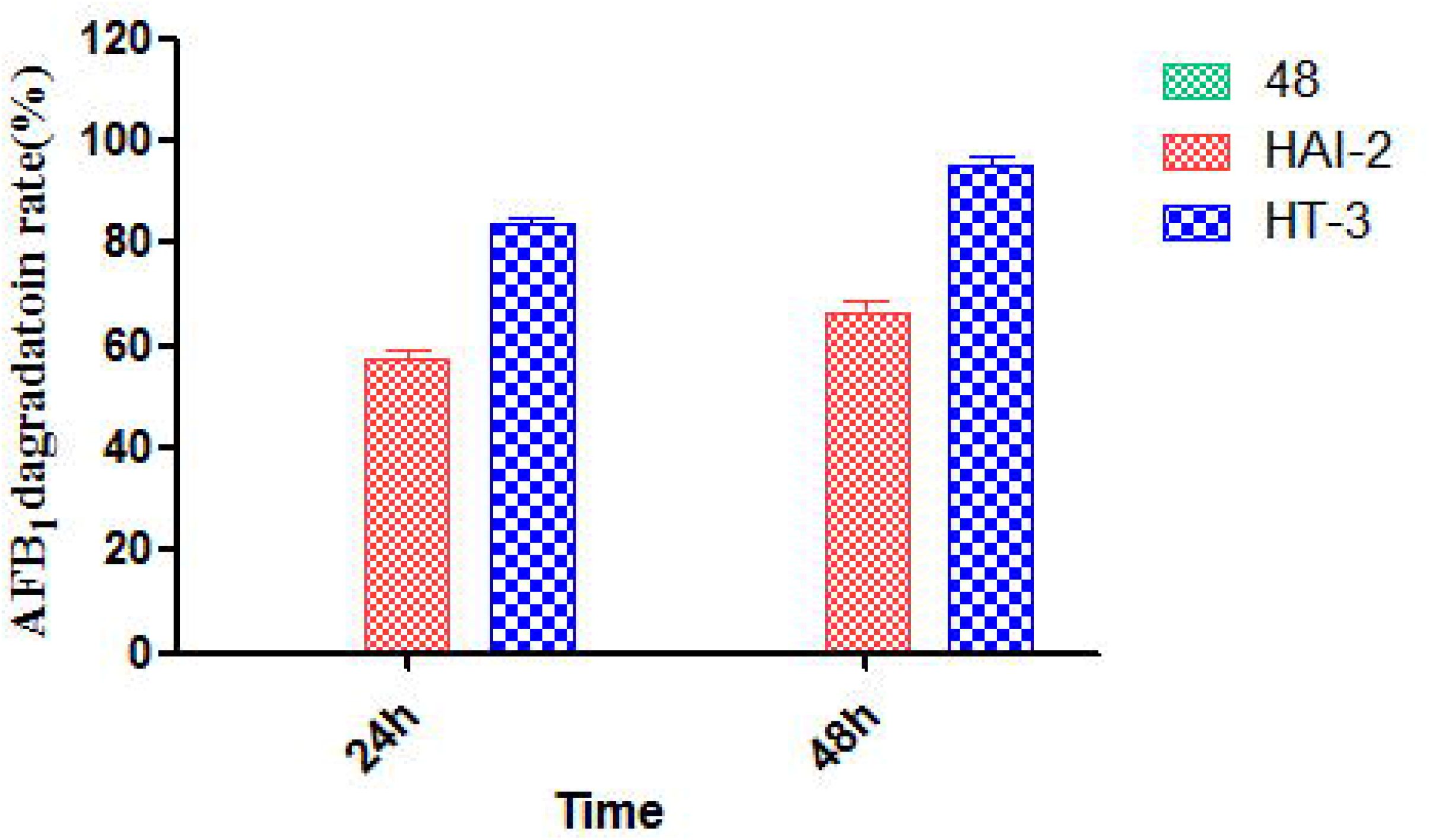
The degradation efficiency of AFB1 by three different strains

### The genome characterization of *Pseudomonas* strains

The genome sequences of two AFB1-degrading strains (HT3 and HAI2) and a non-degrading strain (48) were obtained by Illumina sequencing. Though all 3 genomes are draft assemblies, they represent good sequence quality for performing genomic comparisons (Table 1). The average coverage of the genome sequencing ranges from 100× (strain 48) to 254× (strain HT3), and the contig number of the assemblies was between 11 (strain 48) and 93 (strain HT3). Strain HT3 has the largest genome size (~ 6.59 Mb), while the genome size of strain 48 is smallest (~ 4.72 Mb). Correspondingly, strain HT3 harbors more protein-coding genes (6,039) than the other strains, while strain 48 harbors the lowest number of protein-coding genes (4,238). The GC content of each strain fells into the range of corresponding species whose genome sequences are deposited in the NCBI database (https://www.ncbi.nlm.nih.gov/genome/?term=Pseudomonas), indicating that our sequencing and assembly is reliable.

**Table 1.** Genomic characteristics of *Pseudomonas* strains sequenced in this study

| Strain | HAI2 | HT3 | 48 |
| --- | --- | --- | --- |
| Feature | ABF1 degradation | ABF1 degradation | Non-degradation |
| Sequencing coverage (×) | 236 | 254 | 100 |
| Genome size (bp) | 5,579,564 | 6,588,850 | 4,719,232 |
| Contig number | 79 | 93 | 11 |
| Contig N50 (bp) | 289,640 | 768,697 | 1,097,565 |
| GC content (%) | 61.85 | 66.20 | 62.44 |
| Protein-coding genes | 5,011 | 6,039 | 4,238 |
| rRNA | 72 | 71 | 57 |
| tRNA | 8 | 5 | 5 |

### Functional Protein Classification

The predicted protein sequences of each strain were annotated to the COG database. An average of ~ 81% proteins of these strains were annotated to at least one COG category. Except for categories of general function prediction only and function unknown, the top enriched categories are amino acid transport and metabolism, signal transduction mechanisms, transcription, and energy production and conversion in each strain (Figure 3a). Most of the categories show approximately similar protein proportion and only have slight difference for several functions, indicating a conservation of protein functions in genus *Pseudomonas*.

**Figure 3.**
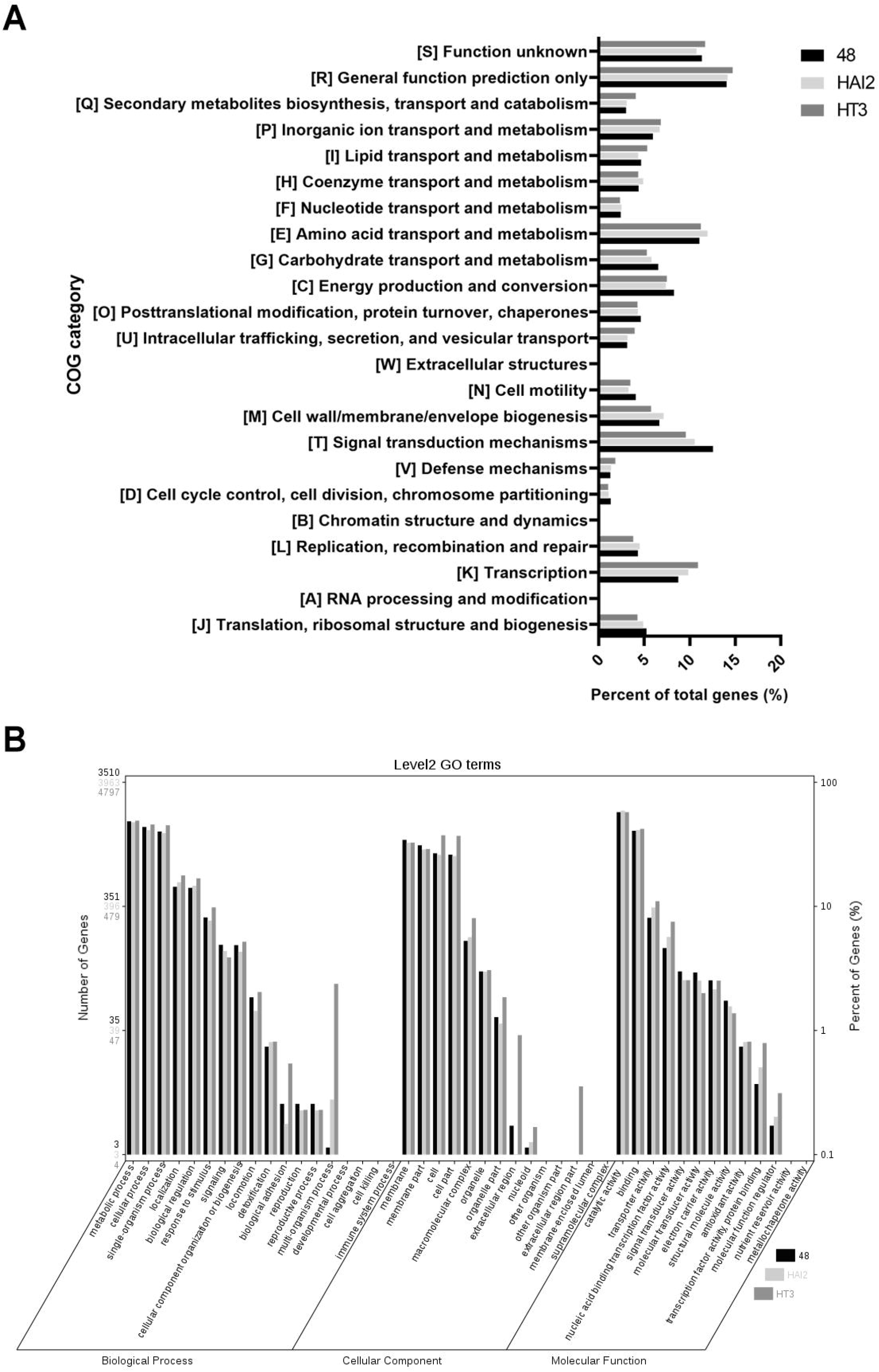
Functional protein classification of *Pseudomonas* strains. (A) Clusters of Orthologous Groups (COG) classification of the protein coding genes. (B) Gene Ontology (GO) functional categories of the protein coding genes.

We then annotated these protein sequences to GO database. For each strain, the most proteins are attributed to GO terms related with catalytic activity, binding, metabolic process, cellular process, and membrane (Figure 3b). Similar to the COG annotation, protein distributions in GO terms show few differences among these *Pseudomonas* strains. These results reinforce the conserved functions of core proteins in *Pseudomonas*.

### Metabolism-related pathway of AFB1-degrading *Pseudomonas* strains

The number of genes enriched in the metabolic and synthetic pathways is 1243 for strain 48, which is fewer than those of strains HAI2 (1323) and HT3 (1411). However, there is no difference in the gene abundance of pathways related to most basic metabolic functions, like carbohydrate metabolism, lipid metabolism, and nucleotide metabolism (Table S2). The two AFB1-degrading *Pseudomonas* strains harbor more genes involved in the metabolism of amino acids than strain 48 (Table S2), indicating a diversity in the nutrition uptake between AFB1-degrading and non-degrading *Pseudomonas* species. Furthermore, the number of genes belonged to the pathway of xenobiotics biodegradation and metabolism in AFB1-degrading *Pseudomonas* strains is obviously larger than that in strain 48 (Table S2). This implies a better metabolic ability in the two strains.

### AFB1 degradation-related enzymes

Previous studies have showed that F420H2 dependent reductases, multicopper oxidase, cytochrome P450, and peroxidase might be involved in the AFB1 degradation of bacteria. Thus, we searched gene families related to these enzymes in the genome sequences. Accordingly, all tested strains were found to harbor one multicopper oxidase (Figure 4a). Though homologous in protein sequences, the multicopper oxidases in strains HT3 is more diverse to those of strains 48 and HAI2 (BLASTP identity cut-off 70%, Figure 4b). HT3 harbor two cytochrome P450 genes, but homologous sequences could not be found in other two strains (Figure 4b). Moreover, the numbers of peroxidase in HT3 (5) and strain HAI2 (7) are higher than strain 48 (2). Notably, the two peroxidases in strain 48 are conserved in all tested strains (Figure 4b), which should be considered as housekeeping genes of genus *Pseudomonas*. In other words, the two AFB1-degrading *Pseudomonas* strains harbor additional peroxidases compared to strain 48. However, F420H2 dependent reductases was absent in all these strains. Together, there are much more AFB1 degradation-related enzymes in *Pseudomonas* strains that have the capability to degrade AFB1. Within them, HT3 might possess different degradation mechanism to that possessed by strain HAI2.

**Figure 4.**
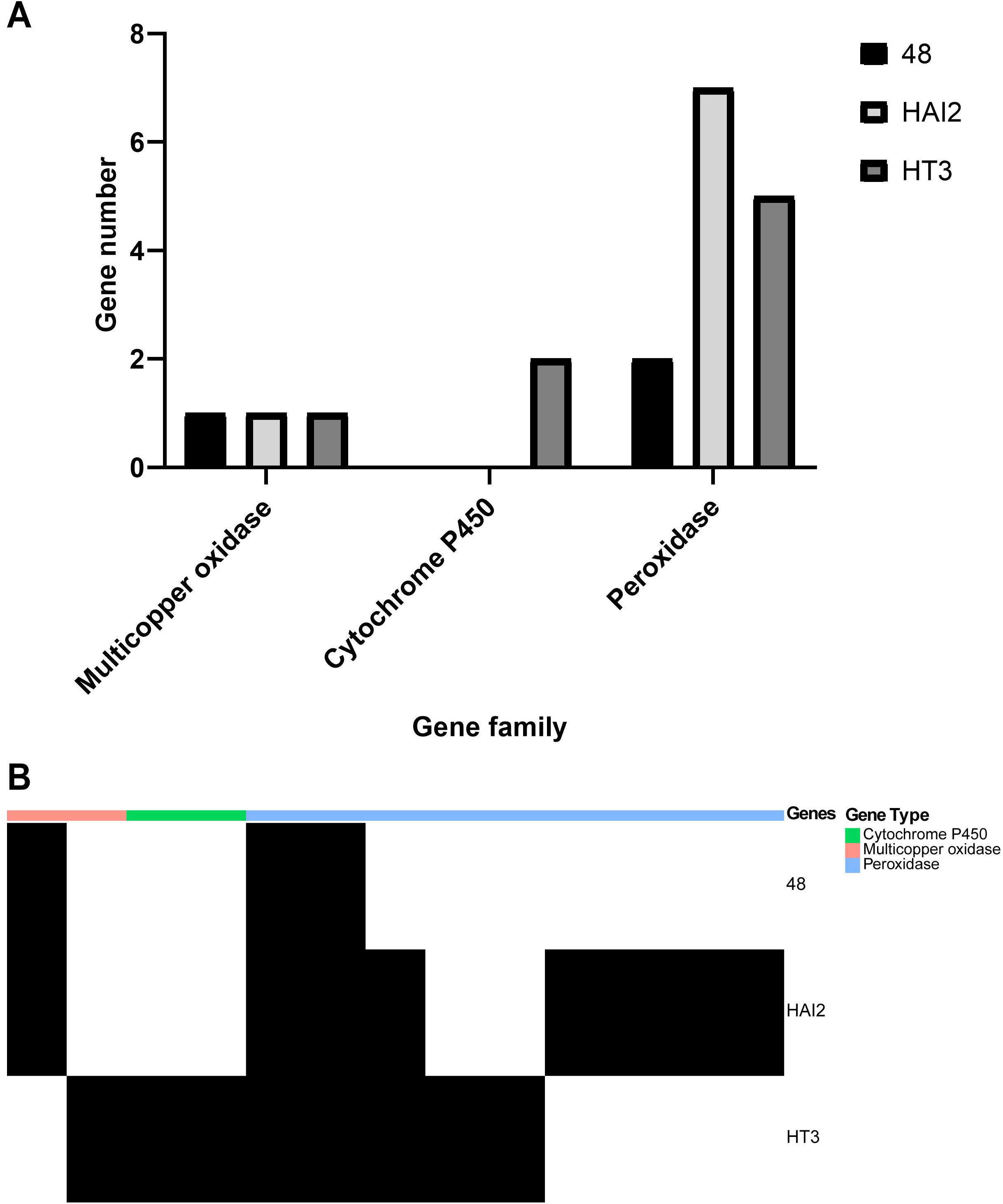
Potential AFB1-degradation enzymes in *Pseudomonas* strains. (A) The number of genes related to three enzyme families in *Pseudomonas* strains. (B) A heatmap for the distribution of corresponding genes in in *Pseudomonas* strains.

### Core proteins of AFB1-degrading *Pseudomonas* strains

We realized that part of the differences observed above might due to the intrinsic genomic diversity between different *Pseudomonas* species. To exclude the false positive and find out potential genes that are exactly involved in AFB1 degradation, we applied a pan-genome analysis on multiple bacterial strains from these four *Pseudomonas* species. The analysis revealed a pan-genome consisting of 16,838 protein-coding genes for these *Pseudomonas* strains (Figure 5a). Within the pan-genome, 1,436 core genes (present in all genomes) were identified. Besides, 8,336 accessory genes (present in some, but not all strains) and 7,066 unique genes (unique to individual strain) were determined (Figure 5a).

**Figure 5.**
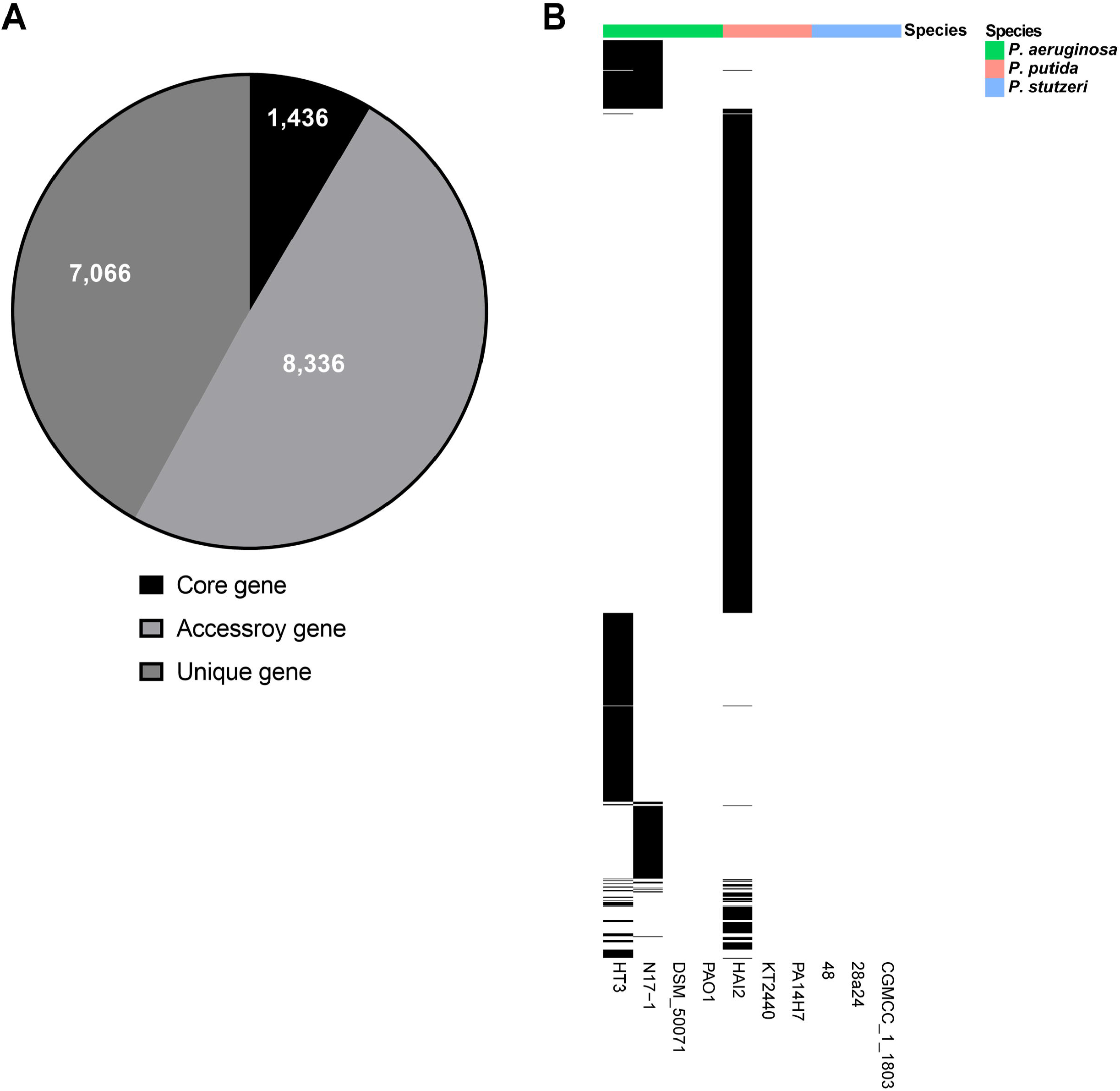
Pan-genome wide analysis of AFB1-degrading *Pseudomonas* strains and non-degrading *Pseudomonas* strains. (A) The pan-genome distribution of *Pseudomonas* strains that was constructed from the genome sequences of 10 strains. (B) A heatmap for the distribution of genes that are specific to AFB1-degrading *Pseudomonas* strains.

It’s obvious that the genes contributing to AFB1 degradation could only be found in accessory and unique genes. We assumed that these genes are present in a least one of the AFB1-degrading *Pseudomonas* strains (including a previous reported strain N17-1), and are absent from all other strains. According to this assumption, we obtained 1,528 genes, of which 1,414 were unique genes, and the less 114 were found in more than one AFB1-degrading strain (Figure 5b). Notably, none of these genes is present in all three AFB1-degrading strains, implying that these strains might utilize different mechanisms to degrade AFB1.

The function of the above genes was analyzed. Forty-five of these genes can’t get any annotation, and 413 are annotated as hypothetical or uncharacterized proteins (Table S3). GO annotation revealed that 347 genes are attributed to the functional category of catalytic activity (Figure 6a), within which 65 genes are related to oxidoreductase activity. However, none of these oxidoreductase activity-related genes is shared by all three AFB1-degrading strains, and only one gene (*viuB*, annotated as siderophore-interacting protein) is shared by two of the AFB1-degrading strains (*P. aeruginosa* HT3 and N17-1) (Figure 6b). These findings indicate that different *Pseudomonas* strains might earned the ability to degrade AFB1 by independently acquiring different catalytic proteins rather than sharing some core proteins.

**Figure 6.**
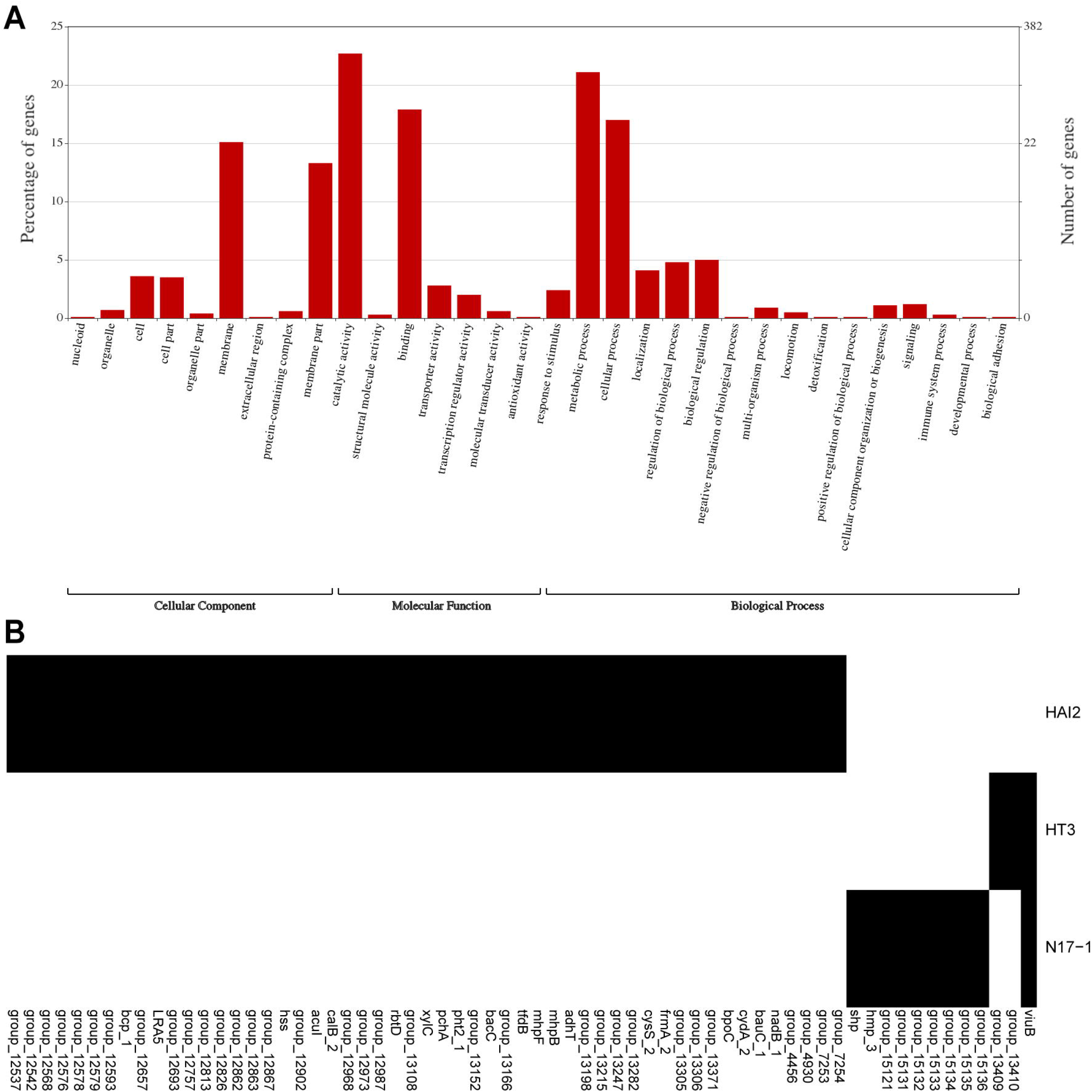
Analysis of genes that are specific to AFB1-degrading *Pseudomonas* strains. (A) Gene Ontology (GO) functional categories of these genes. (B) A heatmap for the distribution of genes that are related to oxidoreductase activity.

## Discussion

Degradation of AFB1 by bacterial genus *Pseudomonas* has been reported for several species, including *P. aeruginosa*, *P. putida*, *P. fluorescens*, and *P. anguilliseptica* (Sangare et al., 2014; Samuel et al., 2014; Adebo et al., 2016; Yang et al., 2017). Our study also presents that a *P. putida* strain and a *P. aeruginosa* strain harbor strong capability to degrade AFB1. This suggests that enzymes contributing to AFB1 degradation might exist in some *Pseudomonas* strains. To give an insight into the genes encoding for these enzymes, we applied whole genome sequencing on the two AFB1-degrading strains and compared them to a non-degrading strain.

The basic functional classification is relatively consistent across these *Pseudomonas* strains. Actually, the genome-wide distribution of COG categories seems to be conserved in this bacterial genus. For example, in *P. aeruginosa* strains F9676 and DN1, *P. alcaliphila* strain JAB1, and *Pseudomonas* sp. strain W15Feb9B, the definite functional categories containing the largest number of genes are amino acid transport and metabolism, signal transduction mechanisms, transcription (Shi et al., 2015; Chauhan et al., 2016; Dong et al., 2017; Ridl et al., 2018), which are also top enriched categories in our tested strains. Thus, the genomic features responsible for AFB1 degradation should not be inherent elements of the genomes of genus *Pseudomonas*. Those AFB1-degrading strains tend to obtain this capacity through some specific horizontal gene transfers.

Some families of enzymes that are previously reported to degrade AFB1 can be found in these *Pseudomonas* strains, including multicopper oxidase, cytochrome P450, and peroxidase. Some of the corresponding genes are conserved in all strains including non-degrading strain 48, and might has no correlation with AFB1 degradation. A significant fact is that strains HAI2 and HT3 harbor much more cytochrome P450 and peroxidase than strain 48. The extra genes in these two strains might contribute to their capacity to degrade AFB1. However, only one of these genes is shared by the two AFB1-degrading strains. Similar observation is also found in the gene sets that are specific present in AFB1-degrading strains. Though we included one more previous published AFB1-degrading strain (Sangare et al., 2014) in our subsequent analysis, the number of specific genes shared in AFB1-degrading strains is still relatively low (Figure 4). Actually, the degradation efficiency of these *Pseudomonas* strains was different, which might be an indicator for the different metabolic mechanisms possessed by these strains.

Interestingly, an oxidoreductase activity-related gene, *viuB*, is shared by AFB1-degrading strains HT3 and N17-1 but absent from all other *Pseudomonas* strains. This gene is commonly known to play a role in iron acquisition of bacteria (Butterton and Calderwood, 1994; Santhanagopalan and Rodriguez, 2012). Some of the catalytic reaction for AFB1 involve the utilization of metal ion, however, it is still unknown whether the uptake of iron by *viuB* might have association to catalyze AFB1. Further study will be conduct to explore the function of *viuB*. Besides, we notice the gene *bpoC* (annotated as α/ β hydrolase) that is named as putative non-heme bromoperoxidase and presents the peroxidase activity (Johnston et al., 2010). Since this gene is unique to strain HAI2, its potential role in AFB1 degradation of this strain deserves a further investigation.

In summary, we have obtained two *Pseudomonas* strains that present strong ability to degrade AFB1. The genome characterizations of these two strains have been clarified, and genes encoding potential catalytic enzymes are identified.

