## supplemental Table 1 for "The genomic context for aflatoxin B1-degrading *Pseudomonas* strains"

Table S1. Reference *Pseudomonas* genomes used in this study

| Species | Strain | Genbank accession No. |
|---|---|---|
| Pseudomonas aeruginosa | N17-1 | GCA_001606045.1_ASM160604v1 |
| Pseudomonas aeruginosa | PAO1 | GCA_000006765.1_ASM676v1 |
| Pseudomonas aeruginosa | DSM_50071 | GCA_001045685.1_ASM104568v1 |
| Pseudomonas putida | KT2440 | GCA_000007565.2_ASM756v2 |
| Pseudomonas putida | PA14H7 | GCA_000800615.1_PseuPA14H7 |
| Pseudomonas stutzeri | 28a24 | GCA_000590475.1_ASM59047v1 |
| Pseudomonas stutzeri | CGMCC_1_1803 | GCA_000219605.1_ASM21960v1 |
