## supplemental Table 2 for "The genomic context for aflatoxin B1-degrading *Pseudomonas* strains"

Table S2. The number of genes involved in KEGG pathways

| KEGG Pathway - level 1 | KEGG Pathway - level 2 | 48 | HAI2 | HT3 |
|---|---|---|---|---|
| Metabolism | Carbohydrate metabolism | 286 | 280 | 289 |
| | Energy metabolism | 147 | 153 | 164 |
| | Lipid metabolism | 78 | 69 | 77 |
| | Nucleotide metabolism | 95 | 93 | 92 |
| | Amino acid metabolism | 262 | 287 | 304 |
| | Metabolism of other amino acids | 49 | 55 | 69 |
| | Glycan biosynthesis and metabolism | 39 | 47 | 47 |
| | Metabolism of cofactors and vitamins | 154 | 172 | 178 |
| | Metabolism of terpenoids and polyketides | 34 | 28 | 42 |
| | Biosynthesis of other secondary metabolites | 43 | 46 | 52 |
| | Xenobiotics biodegradation and metabolism | 56 | 93 | 97 |
| Genetic Information Processing | Transcription | 4 | 5 | 58 |
| | Translation | 88 | 86 | 31 |
| | Folding, sorting and degradation | 44 | 48 | 46 |
| | Replication and repair | 78 | 77 | 73 |
| Environmental Information Processing | Membrane transport | 116 | 173 | 184 |
| | Signal transduction | 110 | 135 | 163 |
| Cellular Processes | Transport and catabolism | 27 | 9 | 7 |
| | Cell growth and death | 10 | 20 | 27 |
| | Cellular community - prokaryotes | 76 | 121 | 193 |
| | Cell motility | 44 | 51 | 52 |
| Organismal Systems | Immune system | 6 | 6 | 6 |
| | Endocrine system | 19 | 13 | 17 |
| | Circulatory system | 5 | 3 | 3 |
| | Digestive system | 3 | 1 | 1 |
| | Excretory system | 1 | 0 | 1 |
| | Nervous system | 5 | 6 | 6 |
| | Development and regeneration | 1 | 2 | 1 |
| | Aging | 14 | 12 | 12 |
| | Environmental adaptation | 13 | 12 | 14 |
