## supplemental Table 3 for "The genomic context for aflatoxin B1-degrading *Pseudomonas* strains"

Table S3. The annotation of genes specificially present in ABF1-degrading strains

| Gene ID | Description | #GO | GO IDs | GO Names |
|---|---|---|---|---|
| group_12521 | Uncharacterised protein | 1 | C:GO:0016021 | C:integral component of membrane |
| group_12522 | type II toxin-antitoxin system HipA family toxin | 2 | F:GO:0016301; P:GO:0016310 | F:kinase activity; P:phosphorylation |
| group_12523 | helix-turn-helix domain-containing protein | 1 | F:GO:0003677 | F:DNA binding |
| group_12524 | methyl-accepting chemotaxis protein | 2 | P:GO:0007165; C:GO:0016021 | P:signal transduction; C:integral component of membrane |
| group_12525 | hypothetical protein | | | |
| group_12526 | DUF2559 family protein | | | |
| group_12527 | cell filamentation protein Fic | 1 | P:GO:0051301 | P:cell division |
| group_12528 | Uncharacterised protein | | | |
| group_12529 | glycosyltransferase family 1 protein | 1 | F:GO:0016757 | F:transferase activity, transferring glycosyl groups |
| group_12530 | ABC transporter | 2 | C:GO:0043190; P:GO:0055085 | C:ATP-binding cassette (ABC) transporter complex; P:transmembrane transport |
| group_12531 | ABC transporter ATP-binding protein | 2 | F:GO:0005524; F:GO:0016887 | F:ATP binding; F:ATPase activity |
| group_12532 | SAM-dependent methyltransferase | 2 | F:GO:0008168; P:GO:0032259 | F:methyltransferase activity; P:methylation |
| group_12533 | glycosyl transferase family 1 | 1 | F:GO:0016740 | F:transferase activity |
| group_12534 | glycosyl transferase family 1 | 1 | F:GO:0016740 | F:transferase activity |
| group_12535 | glycosyl transferase family 1 | 2 | F:GO:0009011; F:GO:0033201 | F:starch synthase activity; F:alpha-1,4-glucan synthase activity |
| group_296 | NAD-dependent dehydratase | 2 | F:GO:0003978; F:GO:0050662 | F:UDP-glucose 4-epimerase activity; F:coenzyme binding |
| group_12536 | acyltransferase | 2 | C:GO:0016021; F:GO:0016747 | component of membrane; F:transferase activity, transferring acyl groups other than amino-acyl groups |
| group_12537 | dTDP-4-dehydrorhamnose reductase | 3 | F:GO:0008831; P:GO:0019305; P:GO:0055114 | ıydrorhamnose reductase activity; P:dTDP-rhamnose biosynthetic process; P:oxidation-reduction process |
| group_1790 | dTDP-glucose 4,6-dehydratase | 2 | F:GO:0008460; P:GO:0009225 | F:dTDP-glucose 4,6-dehydratase activity; P:nucleotide-sugar metabolic process |
| group_12538 | Uncharacterised protein | 1 | F:GO:0016740 | F:transferase activity |
| group_12539 | capsule polysaccharide biosynthesis protein | 2 | P:GO:0000271; P:GO:0015774 | P:polysaccharide biosynthetic process; P:polysaccharide transport |
| group_12540 | glycosyltransferase involved in cell wall biosynthesis | 1 | F:GO:0016740 | F:transferase activity |
| group_12541 | glycosyl transferase | 1 | F:GO:0016740 | F:transferase activity |
| group_12542 | glycosyl transferase | 6 | :0005623; F:GO:0009055; F:GO:0015035; F:GO:0016740; P:GO:0022900; P:GO:00 | ity; F:protein disulfide oxidoreductase activity; F:transferase activity; P:electron transport chain; P:cell redox homeostasis |
| group_12543 | DUF4214 domain-containing protein | 1 | F:GO:0005509 | F:calcium ion binding |
| group_12544 | ligase | 2 | C:GO:0016021; F:GO:0016874 | C:integral component of membrane; F:ligase activity |
| group_4290 | Uncharacterised protein | | | |
| group_55 | DUF3077 domain-containing protein | | | |
| group_663 | 3-deoxy-8-phosphooctulonate synthase | 3 | C:GO:0005737; F:GO:0008676; P:GO:0019294 | deoxy-8-phosphooctulonate synthase activity; P:keto-3-deoxy-D-manno-octulosonic acid biosynthetic process |
| group_12545 | lipopolysaccharide kinase (Kdo/WaaP) family protein | 2 | F:GO:0016301; P:GO:0016310 | F:kinase activity; P:phosphorylation |
| group_4293 | hypothetical protein | | | |
| group_16826 | ---NA--- | | | |
| group_12546 | type 1 fimbrial protein | 2 | P:GO:0007155; C:GO:0009289 | P:cell adhesion; C:pilus |
| fimD_1 | ferrous iron transporter B | 1 | C:GO:0016020 | C:membrane |
| group_12548 | RHS repeat-associated core domain-containing protein | 2 | C:GO:0016020; C:GO:0016021 | C:membrane; C:integral component of membrane |
| group_12549 | bisphenol A degradation protein | | | |
| group_12550 | bisphenol A degradation protein | | | |
| group_3022 | Imidazoleglycerol-phosphate synthase | | | |
| group_6673 | Uncharacterised protein | | | |
| group_3023 | hypothetical protein | | | |
| group_3024 | Imidazoleglycerol-phosphate synthase | | | |
| group_6674 | ---NA--- | | | |
| group_3025 | hypothetical protein | | | |
| group_12551 | hypothetical protein | | | |
| group_12552 | sugar-binding protein | | | |
| group_12553 | Uncharacterised protein | | | |
| group_12554 | autotransporter outer membrane beta-barrel domain-containing protein | 1 | C:GO:0019867 | C:outer membrane |
| group_12555 | hypothetical protein | | | |
| lpfB | phytochrome sensor protein | 1 | P:GO:0016043 | P:cellular component organization |
| group_12557 | type 1 fimbrial protein | 2 | P:GO:0007155; C:GO:0009289 | P:cell adhesion; C:pilus |
| group_12558 | hypothetical protein C7314_4102 | | | |
| group_12559 | hypothetical protein CCOS191_3632 | | | |
| group_12560 | type IV secretion protein Rhs | | | |
| group_12561 | phospholipase | 1 | F:GO:0003824 | F:catalytic activity |
| group_12562 | sel1 repeat family protein | | | |
| group_12563 | MULTISPECIES: hypothetical protein | 1 | C:GO:0016021 | C:integral component of membrane |
| group_12564 | DUF2628 domain-containing protein | 1 | C:GO:0016021 | C:integral component of membrane |
| group_12565 | type IV secretion protein Rhs | 1 | F:GO:0016787 | F:hydrolase activity |
| group_12566 | 2-dehydro-3-deoxyglucarate aldolase | 2 | F:GO:0008672; F:GO:0046872 | F:2-dehydro-3-deoxyglucarate aldolase activity; F:metal ion binding |
| gudP_1 | MFS transporter | 3 | C:GO:0016021; F:GO:0022857; P:GO:0055085 | egral component of membrane; F:transmembrane transporter activity; P:transmembrane transport |
| group_12568 | 2-hydroxyacid dehydrogenase | 3 | F:GO:0030267; F:GO:0051287; P:GO:0055114 | F:glyoxylate reductase (NADP) activity; F:NAD binding; P:oxidation-reduction process |
| tauE | anion permease | 2 | C:GO:0005886; C:GO:0016021 | C:plasma membrane; C:integral component of membrane |
| group_12570 | LacI family DNA-binding transcriptional regulator | 2 | F:GO:0003677; P:GO:0006355 | F:DNA binding; P:regulation of transcription, DNA-templated |
| yidK | solute:sodium symporter family transporter | 3 | C:GO:0016021; F:GO:0022857; P:GO:0055085 | egral component of membrane; F:transmembrane transporter activity; P:transmembrane transport |
| yihQ | alpha-glucosidase | 3 | P:GO:0005975; F:GO:0030246; F:GO:0061634 | :carbohydrate metabolic process; F:carbohydrate binding; F:alpha-D-xyloside xylohydrolase |
| group_12573 | porin | 3 | F:GO:0015288; C:GO:0016020; P:GO:0055085 | F:porin activity; C:membrane; P:transmembrane transport |
| group_12574 | aldose 1-epimerase | 3 | F:GO:0004034; P:GO:0005975; F:GO:0030246 | F:aldose 1-epimerase activity; P:carbohydrate metabolic process; F:carbohydrate binding |
| group_12575 | SMP-30/gluconolactonase/LRE family protein | 2 | F:GO:0004341; F:GO:0046872 | F:gluconolactonase activity; F:metal ion binding |
| group_12576 | 3-oxoacyl-ACP reductase | 5 | F:GO:0004316; F:GO:0047838; P:GO:0055114; F:GO:0102131; F:GO:0102132 | hydrogenase (NAD) activity; P:oxidation-reduction process; F:3-oxo-glutaryl-[acp] methyl ester reductase activity; F:3-oxo-pimeloyl-[acp] methyl ester reductase activity |
| group_12577 | dihydroxy-acid dehydratase | 4 | F:GO:0004160; C:GO:0016021; F:GO:0046872; F:GO:0051536 | dehydratase activity; C:integral component of membrane; F:metal ion binding; F:iron-sulfur cluster binding |
| group_12578 | NAD-dependent succinate-semialdehyde dehydrogenase | 3 | F:GO:0009013; P:GO:0009450; P:GO:0055114 | le dehydrogenase [NAD(P)+] activity; P:gamma-aminobutyric acid catabolic process; P:oxidation-reduction process |
| group_12579 | alcohol dehydrogenase | 6 | 0005506; F:GO:0009055; C:GO:0016020; F:GO:0016614; F:GO:0020037; P:GO:00 | activity; C:membrane; F:oxidoreductase activity, acting on CH-OH group of donors; F:heme binding; P:electron transport chain |
| group_12580 | peptidase M15 | | | |
| group_12581 | hypothetical protein EGN69_19955 | | | |
| group_12583 | Uncharacterised protein | | | |
| group_12584 | histidine kinase | 3 | P:GO:0000160; F:GO:0016301; P:GO:0016310 | P:phosphorelay signal transduction system; F:kinase activity; P:phosphorylation |
| group_12585 | DUF2491 family protein | | | |
| group_12586 | putative membrane protein | 1 | C:GO:0016021 | C:integral component of membrane |
| group_12587 | DUF1190 domain-containing protein | | | |
| ygiC | glutathionylspermidine synthase family protein | 1 | F:GO:0016874 | F:ligase activity |
| group_12589 | putative SOS response-associated peptidase YedK | 2 | P:GO:0006508; F:GO:0008233 | P:proteolysis; F:peptidase activity |
| group_1855 | arsenical efflux pump membrane protein ArsB | 5 | C:GO:0005886; F:GO:0015105; P:GO:0015700; C:GO:0016021; P:GO:0046685 | embrane transporter activity; P:arsenite transport; C:integral component of membrane; P:response to arsenic-containing substance |
| group_12590 | iron transporter | 1 | C:GO:0016021 | C:integral component of membrane |
| group_12591 | PepSY domain-containing protein | 1 | C:GO:0016021 | C:integral component of membrane |
| group_12592 | DUF3325 domain-containing protein | 1 | C:GO:0016021 | C:integral component of membrane |
| group_12593 | biliverdin-producing heme oxygenase | 2 | F:GO:0004392; P:GO:0006788 | F:heme oxygenase (decyclizing) activity; P:heme oxidation |
| pspA | histidine phosphatase family protein | | | |
| group_12595 | ---NA--- | | | |
| group_12601 | MULTISPECIES: hypothetical protein | | | |
| group_12605 | MULTISPECIES: hypothetical protein | 1 | C:GO:0016020 | C:membrane |
| group_12606 | MULTISPECIES: hypothetical protein | | | |
| group_12607 | ---NA--- | | | |
| group_12608 | putative membrane protein | 1 | C:GO:0016021 | C:integral component of membrane |
| group_12616 | hypothetical protein | 1 | F:GO:0016853 | F:isomerase activity |
| group_12617 | response regulator | 1 | P:GO:0000160 | P:phosphorelay signal transduction system |
| group_12618 | hypothetical protein | | | |
| group_12619 | hypothetical protein | | | |
| group_12622 | ---NA--- | | | |
| group_12623 | Uncharacterised protein | | | |
| group_12624 | MFS transporter | 3 | C:GO:0016021; F:GO:0022857; P:GO:0055085 | egral component of membrane; F:transmembrane transporter activity; P:transmembrane transport |
| group_12625 | DUF4440 domain-containing protein | 5 | F:GO:0004683; F:GO:0005516; P:GO:0006468; F:GO:0016301; P:GO:0016310 | t protein kinase activity; F:calmodulin binding; P:protein phosphorylation; F:kinase activity; P:phosphorylation |
| group_12626 | ---NA--- | | | |
| group_232 | metal-binding protein | | | |
| group_12627 | Uncharacterised protein | 1 | C:GO:0016021 | C:integral component of membrane |
| group_12628 | transcriptional regulator | 2 | F:GO:0003677; P:GO:0045892 | F:DNA binding; P:negative regulation of transcription, DNA-templated |
| group_12629 | RHS repeat-associated core domain-containing protein | 2 | C:GO:0016020; C:GO:0016021 | C:membrane; C:integral component of membrane |
| group_12630 | AraC family transcriptional regulator | 3 | F:GO:0003700; P:GO:0006355; F:GO:0043565 | ranscription factor activity; P:regulation of transcription, DNA-templated; F:sequence-specific DNA binding |
| group_12631 | Uncharacterised protein | | | |
| group_12632 | autotransporter outer membrane beta-barrel domain-containing protein | 3 | F:GO:0004725; C:GO:0019867; P:GO:0035335 | rotein tyrosine phosphatase activity; C:outer membrane; P:peptidyl-tyrosine dephosphorylation |
| group_12633 | ---NA--- | | | |
| lutR_2 | GntR family transcriptional regulator | 3 | F:GO:0003677; F:GO:0003700; P:GO:0006355 | binding; F:DNA-binding transcription factor activity; P:regulation of transcription, DNA-templated |
| group_12635 | hypothetical protein | | | |
| bcp_1 | pseudoazurin | 3 | F:GO:0005507; F:GO:0009055; P:GO:0022900 | F:copper ion binding; F:electron transfer activity; P:electron transport chain |
| group_12637 | Uncharacterised protein | | | |
| group_12638 | Uncharacterised protein | | | |
| group_12639 | iron-containing redox enzyme family protein | 2 | F:GO:0008168; P:GO:0032259 | F:methyltransferase activity; P:methylation |
| group_12640 | Uncharacterised protein | | | |
| group_12641 | amidase | 3 | F:GO:0004040; F:GO:0016740; F:GO:0050537 | F:amidase activity; F:transferase activity; F:mandelamide amidase activity |
| group_12642 | acyl-CoA synthetase | 1 | F:GO:0016874 | F:ligase activity |
| group_12643 | acyl carrier protein | | | |
| group_12644 | Uncharacterised protein | | | |
| group_12645 | prenyltransferase | 1 | F:GO:0016740 | F:transferase activity |
| group_12646 | coenzyme F390 synthetase | 1 | F:GO:0016874 | F:ligase activity |
| group_12647 | sigma-54-dependent Fis family transcriptional regulator | 4 | F:GO:0005524; P:GO:0006355; F:GO:0008134; F:GO:0043565 | regulation of transcription, DNA-templated; F:transcription factor binding; F:sequence-specific DNA binding |
| group_12648 | conjugal transfer protein TraX | 1 | C:GO:0016021 | C:integral component of membrane |
| group_12649 | Uncharacterised protein | 1 | C:GO:0016021 | C:integral component of membrane |
| group_12650 | Uncharacterised protein | | | |
| group_12651 | Uncharacterised protein | | | |
| group_12652 | hypothetical protein | | | |
| group_12653 | Uncharacterised protein | | | |
| group_12654 | MULTISPECIES: hypothetical protein | 1 | C:GO:0016021 | C:integral component of membrane |
| group_12655 | DNA-binding protein | 5 | F:GO:0003677; F:GO:0003700; P:GO:0006355; F:GO:0008483; F:GO:0030170 | scription factor activity; P:regulation of transcription, DNA-templated; F:transaminase activity; F:pyridoxal phosphate binding |
| group_12656 | GNAT family N-acetyltransferase | 1 | F:GO:0008080 | F:N-acetyltransferase activity |
| group_12657 | NAD(P)H dehydrogenase | 5 | F:GO:0009055; F:GO:0010181; F:GO:0016652; F:GO:0016661; P:GO:0022900 | ctivity, acting on NAD(P)H, NAD(P) as acceptor; F:oxidoreductase activity, acting on other nitrogenous compounds as donors; P:electron transport chain |
| group_12658 | DUF2599 domain-containing protein | 1 | F:GO:0016787 | F:hydrolase activity |
| group_12659 | DUF2599 domain-containing protein | 1 | F:GO:0016787 | F:hydrolase activity |
| group_12660 | DUF2599 domain-containing protein | | | |
| group_12661 | heme utilization protein | | | |
| group_12662 | cobalamin biosynthesis protein CobE | 1 | P:GO:0009236 | P:cobalamin biosynthetic process |
| group_12663 | aminotransferase class I/II-fold pyridoxal phosphate-dependent enzyme | 3 | F:GO:0008483; P:GO:0009058; F:GO:0030170 | F:transaminase activity; P:biosynthetic process; F:pyridoxal phosphate binding |
| group_12665 | cytochrome oxidase I | | | |
| group_12666 | hypothetical protein | | | |
| group_12667 | sugar phosphate isomerase/epimerase | 1 | F:GO:0016853 | F:isomerase activity |
| LRA5 | 3-oxoacyl-ACP reductase | 2 | F:GO:0016491; P:GO:0055114 | F:oxidoreductase activity; P:oxidation-reduction process |
| gudP_3 | MFS transporter | 3 | C:GO:0016021; F:GO:0022857; P:GO:0055085 | egral component of membrane; F:transmembrane transporter activity; P:transmembrane transport |
| group_12670 | 2-pyrone-4,6-dicarboxylate hydrolase | 2 | F:GO:0018732; F:GO:0102998 | F:sulfolactone hydrolase activity; F:4-sulfomuconolactone hydrolase activity |
| group_12671 | LysR family transcriptional regulator | 3 | F:GO:0003677; F:GO:0003700; P:GO:0006355 | binding; F:DNA-binding transcription factor activity; P:regulation of transcription, DNA-templated |
| group_12672 | DUF3053 domain-containing protein | | | |
| group_12673 | Uncharacterised protein | | | |
| group_12674 | formate/nitrite transporter | 3 | C:GO:0016021; F:GO:0022857; P:GO:0055085 | egral component of membrane; F:transmembrane transporter activity; P:transmembrane transport |
| group_12675 | histidine kinase | 2 | F:GO:0004673; P:GO:0018106 | F:protein histidine kinase activity; P:peptidyl-histidine phosphorylation |
| group_12676 | membrane protein | 1 | C:GO:0016021 | C:integral component of membrane |
| group_12677 | fluoride efflux transporter CrcB | 4 | C:GO:0005887; F:GO:0015103; P:GO:0015698; P:GO:0034220 | nembrane; F:inorganic anion transmembrane transporter activity; P:inorganic anion transport; P:ion transmembrane transport |
| group_12678 | LuxR family transcriptional regulator | 2 | F:GO:0003677; P:GO:0006355 | F:DNA binding; P:regulation of transcription, DNA-templated |
| group_12679 | malonate decarboxylase subunit epsilon | 1 | F:GO:0004314 | F:[acyl-carrier-protein] S-malonyltransferase activity |
| group_12680 | malonate decarboxylase holo-ACP synthase | 1 | F:GO:0016779 | F:nucleotidyltransferase activity |
| group_12681 | triphosphoribosyl-dephospho-CoA synthase | 4 | F:GO:0005524; P:GO:0016310; F:GO:0016757; F:GO:0046917 | orylation; F:transferase activity, transferring glycosyl groups; F:triphosphoribosyl-dephospho-CoA synthase activity |
| group_12682 | sulfatase | 3 | F:GO:0008484; C:GO:0016021; F:GO:0016772 | se activity; C:integral component of membrane; F:transferase activity, transferring phosphorus-containing groups |
| group_12683 | twin-arginine translocation signal domain-containing protein | 1 | C:GO:0016021 | C:integral component of membrane |
| group_12684 | ---NA--- | | | |
| group_12685 | leucine-rich repeat domain-containing protein | | | |
| group_12686 | UDP-N-acetylglucosamine 2-epimerase (non-hydrolyzing) | 1 | F:GO:0008761 | F:UDP-N-acetylglucosamine 2-epimerase activity |
| group_12687 | delta-60 repeat domain-containing protein | | | |
| hup | HU family DNA-binding protein | 2 | F:GO:0003677; P:GO:0030261 | F:DNA binding; P:chromosome condensation |
| group_12689 | E3 ligase-like protein (putative virulence factor) | 3 | F:GO:0004842; C:GO:0016020; P:GO:0016567 | F:ubiquitin-protein transferase activity; C:membrane; P:protein ubiquitination |
| group_12690 | GntR family transcriptional regulator | 5 | F:GO:0003677; F:GO:0003700; P:GO:0006355; F:GO:0008483; F:GO:0030170 | scription factor activity; P:regulation of transcription, DNA-templated; F:transaminase activity; F:pyridoxal phosphate binding |
| group_12691 | Uncharacterised protein | | | |
| group_12692 | His-Xaa-Ser system protein HxsD | | | |
| group_12693 | His-Xaa-Ser system radical SAM maturase HxsB | 3 | F:GO:0016491; F:GO:0051536; P:GO:0055114 | F:oxidoreductase activity; F:iron-sulfur cluster binding; P:oxidation-reduction process |
| group_12694 | His-Xaa-Ser system radical SAM maturase HxsC | 2 | F:GO:0003824; F:GO:0051536 | F:catalytic activity; F:iron-sulfur cluster binding |
| group_12695 | His-Xaa-Ser repeat protein HxsA | | | |
| group_12696 | hypothetical protein | | | |
| group_12697 | hypothetical protein | | | |
| group_12699 | hypothetical protein | 1 | C:GO:0016021 | C:integral component of membrane |
| group_12700 | chromate transporter | 3 | F:GO:0015109; P:GO:0015703; C:GO:0016021 | nate transmembrane transporter activity; P:chromate transport; C:integral component of membrane |
| group_12701 | Uncharacterised protein | 1 | C:GO:0016021 | C:integral component of membrane |
| group_12702 | universal stress protein | | | |
| group_12703 | hypothetical protein | | | |
| group_12704 | universal stress protein | | | |
| group_12705 | ---NA--- | | | |
| group_12706 | lipoprotein | 2 | F:GO:0003674; P:GO:0008150 | F:molecular_function; P:biological_process |
| lprI | DUF1311 domain-containing protein | | | |
| group_12708 | serine hydrolase | 1 | F:GO:0016787 | F:hydrolase activity |
| group_12709 | DUF4303 domain-containing protein | | | |
| group_12710 | hypothetical protein | | | |
| group_12711 | Uncharacterised protein | | | |
| group_12712 | Possible hemagglutinin | | | |
| group_12713 | hypothetical protein YSA_03352 | | | |
| group_12714 | Uncharacterised protein | | | |
| group_12716 | DUF3077 domain-containing protein | | | |
| group_12715 | Uncharacterised protein | | | |
| group_12717 | DUF3077 domain-containing protein | | | |
| group_12718 | nucleotide pyrophosphohydrolase | 3 | P:GO:0009143; F:GO:0046872; F:GO:0047429 | triphosphate catabolic process; F:metal ion binding; F:nucleoside-triphosphate diphosphatase activity |
| group_12719 | DUF2075 domain-containing protein | 1 | F:GO:0005524 | F:ATP binding |
| group_111 | Uncharacterised protein | | | |

| | | | | |
|---|---|---|---|---|
| group_12720 | hypothetical protein | 1 | C:GO:0016021 | C:integral component of membrane |
| group_12721 | adenylyl-sulfate kinase | 6 | :0000103; F:GO:0004020; F:GO:0005524; C:GO:0016020; P:GO:0016310; P:GO:00 | nylylsulfate kinase activity; F:ATP binding; C:membrane; P:phosphorylation; P:hydrogen sulfide biosynthetic process |
| group_12722 | TolC family protein | 2 | F:GO:0015562; P:GO:0055085 | F:efflux transmembrane transporter activity; P:transmembrane transport |
| group_12723 | DUF4347 domain-containing protein | 2 | F:GO:0004519; P:GO:0090305 | F:endonuclease activity; P:nucleic acid phosphodiester bond hydrolysis |
| group_12724 | phage tail protein | | | |
| group_12725 | N-acetyltransferase | 2 | C:GO:0005840; F:GO:0008080 | C:ribosome; F:N-acetyltransferase activity |
| group_12726 | sulfotransferase family protein | 1 | F:GO:0016740 | F:transferase activity |
| group_12727 | aspartyl beta-hydroxylase | 1 | P:GO:0018193 | P:peptidyl-amino acid modification |
| group_12728 | peptidase M50 | 3 | F:GO:0004222; P:GO:0006508; C:GO:0016021 | F:metalloendopeptidase activity; P:proteolysis; C:integral component of membrane |
| group_12729 | multidrug transporter | | | |
| group_12730 | efflux RND transporter periplasmic adaptor subunit | 3 | C:GO:0005886; F:GO:0022857; P:GO:0055085 | C:plasma membrane; F:transmembrane transporter activity; P:transmembrane transport |
| gudP_4 | MFS transporter | 3 | C:GO:0016021; F:GO:0022857; P:GO:0055085 | egral component of membrane; F:transmembrane transporter activity; P:transmembrane transport |
| group_12732 | hypothetical protein | | | |
| group_239 | ATP-dependent helicase HrpB | 3 | F:GO:0003676; F:GO:0004386; F:GO:0005524 | F:nucleic acid binding; F:helicase activity; F:ATP binding |
| group_12733 | MULTISPECIES: hypothetical protein | | | |
| group_12734 | integrase | 3 | F:GO:0003677; P:GO:0006310; P:GO:0015074 | F:DNA binding; P:DNA recombination; P:DNA integration |
| group_12735 | MULTISPECIES: hypothetical protein | | | |
| group_12736 | integrase | 3 | F:GO:0003677; P:GO:0006310; P:GO:0015074 | F:DNA binding; P:DNA recombination; P:DNA integration |
| group_12737 | Uncharacterised protein | 1 | C:GO:0016021 | C:integral component of membrane |
| group_12738 | zinc chelation protein SecC | 1 | F:GO:0005524 | F:ATP binding |
| group_12739 | Uncharacterised protein | | | |
| group_12740 | hypothetical protein ALQ23_200213 | | | |
| group_12741 | metallophosphoesterase | 1 | F:GO:0016787 | F:hydrolase activity |
| group_12742 | transposase | 3 | F:GO:0004803; P:GO:0006313; F:GO:0043565 | F:transposase activity; P:transposition, DNA-mediated; F:sequence-specific DNA binding |
| group_3030 | transposase | | | |
| group_12743 | DUF3732 domain-containing protein | 4 | P:GO:0006508; F:GO:0008237; F:GO:0008241; C:GO:0016020 | P:proteolysis; F:metallopeptidase activity; F:peptidyl-dipeptidase activity; C:membrane |
| group_12744 | Uncharacterised protein | | | |
| group_12745 | Uncharacterised protein | | | |
| group_12746 | hypothetical protein | | | |
| group_12747 | SH3 domain-containing protein | | | |
| group_3038 | transcriptional regulator | 2 | F:GO:0003677; P:GO:0006355 | F:DNA binding; P:regulation of transcription, DNA-templated |
| group_12748 | phosphotransferase | 2 | F:GO:0005524; F:GO:0016740 | F:ATP binding; F:transferase activity |
| group_12749 | ---NA--- | | | |
| group_12750 | serine/threonine protein phosphatase | 1 | F:GO:0016787 | F:hydrolase activity |
| group_12751 | Uncharacterised protein | | | |
| group_12752 | hypothetical protein | | | |
| group_12753 | Fis family transcriptional regulator | 1 | F:GO:0003677 | F:DNA binding |
| group_12754 | Permuted papain-like amidase enzyme, YaeF/YiiX, C92 family | 1 | F:GO:0016787 | F:hydrolase activity |
| group_241 | ATP-dependent helicase HrpB | 3 | F:GO:0003676; F:GO:0004386; F:GO:0005524 | F:nucleic acid binding; F:helicase activity; F:ATP binding |
| group_12755 | HNH endonuclease | | | |
| group_12756 | XRE family transcriptional regulator | 1 | F:GO:0003677 | F:DNA binding |
| group_117 | Uncharacterised protein | | | |
| group_12757 | TfdA family taurine catabolism dioxygenase TauD | 3 | F:GO:0005506; F:GO:0051213; P:GO:0055114 | F:iron ion binding; F:dioxygenase activity; P:oxidation-reduction process |
| group_12758 | NAD-dependent DNA ligase LigB | 4 | F:GO:0003911; P:GO:0006260; P:GO:0006266; P:GO:0006281 | F:DNA ligase (NAD+) activity; P:DNA replication; P:DNA ligation; P:DNA repair |
| yqjC_2 | DUF1090 domain-containing protein | 2 | F:GO:0008168; P:GO:0032259 | F:methyltransferase activity; P:methylation |
| group_12760 | ---NA--- | | | |
| group_12761 | Uncharacterised protein | | | |
| group_12762 | hypothetical protein | | | |
| group_12763 | E3 ligase-like protein (putative virulence factor) | 2 | F:GO:0004842; P:GO:0016567 | F:ubiquitin-protein transferase activity; P:protein ubiquitination |
| group_12764 | ---NA--- | | | |
| group_12765 | hypothetical protein | | | |
| group_4292 | Uncharacterised protein | | | |
| group_58 | DUF3077 domain-containing protein | | | |
| group_12766 | PhoPQ-activated pathogenicity protein | | | |
| group_12767 | integrase | 4 | F:GO:0003677; F:GO:0004803; P:GO:0006313; P:GO:0015074 | :DNA binding; F:transposase activity; P:transposition, DNA-mediated; P:DNA integration |
| group_12768 | Protein of uncharacterised function (DUF3077) | | | |
| group_6907 | DUF3077 domain-containing protein | | | |
| group_12769 | Uncharacterised protein | | | |
| group_6908 | DUF3077 domain-containing protein | | | |
| group_12770 | Uncharacterised protein | | | |
| group_12771 | ynthetase catalytic domain-containing protein (RelA/SpoT-type nucleotidyltra | 1 | P:GO:0015969 | P:guanosine tetraphosphate metabolic process |
| group_12772 | tail assembly protein | 1 | C:GO:0016021 | C:integral component of membrane |
| group_12773 | hypothetical protein | | | |
| group_12774 | hypothetical protein | | | |
| group_12775 | phage major capsid protein | | | |
| group_12776 | DNA-binding protein | 1 | F:GO:0003677 | F:DNA binding |
| group_12777 | integrase | 3 | F:GO:0003677; P:GO:0006310; P:GO:0015074 | F:DNA binding; P:DNA recombination; P:DNA integration |
| group_12778 | Uncharacterised protein | | | |
| group_12779 | AAA family ATPase | | | |
| group_12780 | virulence-associated protein E | 3 | F:GO:0003677; P:GO:0006260; F:GO:0008270 | F:DNA binding; P:DNA replication; F:zinc ion binding |
| group_12781 | hypothetical protein | | | |
| group_12782 | Uncharacterised protein | | | |
| group_12783 | hypothetical protein SAMN05660967_03408 | | | |
| group_12784 | AlpA family phage regulatory protein | | | |
| group_12785 | Uncharacterised protein | | | |
| intS | integrase | 3 | F:GO:0003677; P:GO:0006310; P:GO:0015074 | F:DNA binding; P:DNA recombination; P:DNA integration |
| group_12787 | membrane protein | 1 | C:GO:0016021 | C:integral component of membrane |
| group_12788 | ---NA--- | | | |
| group_12789 | S-type Pyocin | 2 | P:GO:0009987; P:GO:0044419 | P:cellular process; P:interspecies interaction between organisms |
| imm2 | Pyocin-S2 immunity protein | 2 | F:GO:0015643; P:GO:0030153 | F:toxic substance binding; P:bacteriocin immunity |
| group_12791 | PHP-associated family protein | 1 | F:GO:0003824 | F:catalytic activity |
| group_12792 | hdig domain protein | | | |
| group_12793 | HD domain-containing protein | 1 | F:GO:0016787 | F:hydrolase activity |
| pphA | phosphoenolpyruvate mutase | 1 | F:GO:0050188 | F:phosphoenolpyruvate mutase activity |
| group_12795 | MFS transporter | 3 | C:GO:0016020; C:GO:0016021; P:GO:0055085 | C:membrane; C:integral component of membrane; P:transmembrane transport |
| group_12796 | Heterokaryon incompatibility protein Het-C | | | |
| group_12797 | type II secretion system protein GspL | 1 | C:GO:0016021 | C:integral component of membrane |
| group_12798 | type II secretion system protein GspM | 1 | C:GO:0016021 | C:integral component of membrane |
| group_12799 | general secretion pathway protein GspN | | | |
| group_12800 | ShlB/FhaC/HecB family hemolysin secretion/activation protein | 1 | P:GO:0015031 | P:protein transport |
| group_12801 | histidine kinase | 2 | F:GO:0016301; P:GO:0016310 | F:kinase activity; P:phosphorylation |
| group_56 | DUF3077 domain-containing protein | | | |
| group_4291 | Uncharacterised protein | | | |
| group_12802 | ---NA--- | | | |
| group_12803 | hypothetical protein | | | |
| group_12804 | phosphoethanolamine transferase | 3 | F:GO:0008484; C:GO:0016021; F:GO:0016772 | ase activity; C:integral component of membrane; F:transferase activity, transferring phosphorus-containing groups |
| group_12805 | DNA-binding response regulator | 3 | P:GO:0000160; F:GO:0003677; P:GO:0006355 | horelay signal transduction system; F:DNA binding; P:regulation of transcription, DNA-templated |
| group_12806 | phosphatase PAP2 family protein | 1 | C:GO:0016021 | C:integral component of membrane |
| group_12807 | lipoprotein, putative | | | |
| ais | histidine phosphatase family protein | 1 | C:GO:0016021 | C:integral component of membrane |
| group_12809 | PAP2 family protein | 1 | C:GO:0016021 | C:integral component of membrane |
| group_12810 | hypothetical protein DI560_24990 | | | |
| group_12811 | isochorismatase | 1 | F:GO:0050127 | F:N-carbamoylsarcosine amidase activity |
| group_12812 | LysR family transcriptional regulator | 3 | F:GO:0003677; F:GO:0003700; P:GO:0006355 | binding; F:DNA-binding transcription factor activity; P:regulation of transcription, DNA-templated |
| group_12813 | NADP-dependent 3-hydroxy acid dehydrogenase YdfG | 2 | F:GO:0016491; P:GO:0055114 | F:oxidoreductase activity; P:oxidation-reduction process |
| group_12814 | MULTISPECIES: hypothetical protein | | | |
| gloA_1 | lactoylglutathione lyase | 4 | F:GO:0004462; C:GO:0005737; P:GO:0019243; F:GO:0046872 | activity; C:cytoplasm; P:methylglyoxal catabolic process to D-lactate via S-lactoyl-glutathione; F:metal ion binding |
| group_12816 | methyltransferase | 2 | F:GO:0008168; P:GO:0032259 | F:methyltransferase activity; P:methylation |
| group_12817 | MULTISPECIES: hypothetical protein | | | |
| group_12818 | lipoprotein | | | |
| group_93 | arsenate reductase | | | |
| group_12819 | Translation initiation factor 2 | 2 | F:GO:0003743; P:GO:0006413 | F:translation initiation factor activity; P:translational initiation |
| group_12820 | formylglycine-generating enzyme family protein | 2 | F:GO:0016301; P:GO:0016310 | F:kinase activity; P:phosphorylation |
| group_12821 | amino acid adenylation domain-containing protein | 2 | F:GO:0003824; F:GO:0031177 | F:catalytic activity; F:phosphopantetheine binding |
| group_12822 | amino acid adenylation domain-containing protein | 3 | P:GO:0009058; F:GO:0016788; F:GO:0031177 | iosynthetic process; F:hydrolase activity, acting on ester bonds; F:phosphopantetheine binding |
| group_12823 | lipase | 1 | F:GO:0004806 | F:triglyceride lipase activity |
| group_12824 | TonB-dependent siderophore receptor | 4 | C:GO:0009279; F:GO:0015344; F:GO:0038023; P:GO:0044718 | erophore uptake transmembrane transporter activity; F:signaling receptor activity; P:siderophore transmembrane transport |
| tycB | non-ribosomal peptide synthetase | 2 | F:GO:0003824; F:GO:0031177 | F:catalytic activity; F:phosphopantetheine binding |
| group_12826 | non-ribosomal peptide synthetase | 3 | F:GO:0016491; F:GO:0031177; P:GO:0055114 | F:oxidoreductase activity; F:phosphopantetheine binding; P:oxidation-reduction process |
| group_12827 | hypothetical protein WR25_08626 | | | |
| group_12828 | glutamyl-tRNA amidotransferase | 1 | F:GO:0016740 | F:transferase activity |
| group_12829 | glutamyl-tRNA amidotransferase | 1 | F:GO:0016740 | F:transferase activity |
| group_12830 | EamA family transporter | 1 | C:GO:0016021 | C:integral component of membrane |
| group_12831 | hypothetical protein YSA_08300 | | | |
| group_12832 | hypothetical protein | 2 | C:GO:0016020; C:GO:0016021 | C:membrane; C:integral component of membrane |
| group_12833 | leucine-rich repeat-containing protein typical subtype | 3 | F:GO:0004842; C:GO:0016021; P:GO:0016567 | iquitin-protein transferase activity; C:integral component of membrane; P:protein ubiquitination |
| group_12834 | mannosyltransferase | 1 | F:GO:0016757 | F:transferase activity, transferring glycosyl groups |
| group_12835 | mannosyltransferase | 1 | F:GO:0016757 | F:transferase activity, transferring glycosyl groups |
| group_12836 | ---NA--- | | | |
| group_12837 | Hpt domain-containing protein | 3 | P:GO:0000160; F:GO:0016301; P:GO:0016310 | P:phosphorelay signal transduction system; F:kinase activity; P:phosphorylation |
| group_12838 | flagellar hook protein FliD | 5 | C:GO:0005576; P:GO:0007155; C:GO:0009421; C:GO:0009424; P:GO:0071973 | on; C:bacterial-type flagellum filament cap; C:bacterial-type flagellum hook; P:bacterial-type flagellum-dependent cell motility |
| group_12839 | flagellar protein FlaG | | | |
| group_12840 | flagellin | 4 | F:GO:0005198; C:GO:0005576; C:GO:0009420; P:GO:0071973 | ivity; C:extracellular region; C:bacterial-type flagellum filament; P:bacterial-type flagellum-dependent cell motility |
| group_12841 | hypothetical protein | | | |
| group_12842 | DUF115 domain-containing protein | | | |
| pseI | pseudaminic acid synthase | 3 | P:GO:0016051; F:GO:0047444; F:GO:0050462 | ynthetic process; F:N-acylneuraminate-9-phosphate synthase activity; F:N-acetylneuraminate synthase activity |
| murG_1 | UDP-2,4-diacetamido-2,4,6-trideoxy-beta-L-altropyranose hydrolase | 3 | F:GO:0008080; F:GO:0016758; F:GO:0016787 | acetyltransferase activity; F:transferase activity, transferring hexosyl groups; F:hydrolase activity |
| neuA | pseudaminic acid cytidylyltransferase | 1 | F:GO:0008781 | F:N-acylneuraminate cytidylyltransferase activity |
| group_12846 | UDP-4-amino-4,6-dideoxy-N-acetyl-beta-L-altrosamine transaminase | 1 | F:GO:0008483 | F:transaminase activity |
| pseB | UDP-N-acetylglucosamine 4,6-dehydratase (inverting) | 2 | F:GO:0008483; F:GO:0016829 | F:transaminase activity; F:lyase activity |
| group_12848 | glycosyl transferase family 2 | 1 | F:GO:0016740 | F:transferase activity |
| group_12849 | cephalosporin hydroxylase | 2 | F:GO:0008168; P:GO:0008610 | F:methyltransferase activity; P:lipid biosynthetic process |
| group_12850 | methyltransferase domain-containing protein | 2 | F:GO:0008168; P:GO:0032259 | F:methyltransferase activity; P:methylation |
| tll | NAD(P)-dependent oxidoreductase | 2 | F:GO:0003824; F:GO:0050662 | F:catalytic activity; F:coenzyme binding |
| group_12852 | SAM-dependent methyltransferase | 2 | F:GO:0008168; P:GO:0032259 | F:methyltransferase activity; P:methylation |
| rfbC | dTDP-4-dehydrorhamnose 3,5-epimerase | 1 | F:GO:0008830 | F:dTDP-4-dehydrorhamnose 3,5-epimerase activity |
| rfbG | CDP-glucose 4,6-dehydratase | 1 | F:GO:0047733 | F:CDP-glucose 4,6-dehydratase activity |
| rfbF | glucose-1-phosphate cytidylyltransferase | 2 | P:GO:0009058; F:GO:0047343 | P:biosynthetic process; F:glucose-1-phosphate cytidylyltransferase activity |
| group_12856 | glycosyl transferase family 2 | 1 | F:GO:0016740 | F:transferase activity |
| group_12857 | flagellar hook-associated protein 3 | 4 | F:GO:0005198; C:GO:0009420; C:GO:0009424; P:GO:0071973 | ; C:bacterial-type flagellum filament; C:bacterial-type flagellum hook; P:bacterial-type flagellum-dependent cell motility |
| group_12858 | flagellar hook-associated protein FlgK | 4 | F:GO:0005198; C:GO:0009424; P:GO:0044780; P:GO:0071973 | C:bacterial-type flagellum hook; P:bacterial-type flagellum assembly; P:bacterial-type flagellum-dependent cell motility |
| group_12859 | Uncharacterised protein | | | |
| group_12860 | flagellar biosynthesis protein FlgD | 1 | P:GO:0044781 | P:bacterial-type flagellum organization |
| group_12861 | dTDP-6-deoxy-L-hexose 3-O-methyltransferase | 2 | F:GO:0008168; P:GO:0032259 | F:methyltransferase activity; P:methylation |
| group_12862 | hypothetical protein | 2 | F:GO:0016491; P:GO:0055114 | F:oxidoreductase activity; P:oxidation-reduction process |
| group_12863 | phytanoyl-CoA dioxygenase PhyH | 2 | F:GO:0051213; P:GO:0055114 | F:dioxygenase activity; P:oxidation-reduction process |
| group_12864 | sugar O-acyltransferase | 1 | F:GO:0016746 | F:transferase activity, transferring acyl groups |
| group_12865 | hypothetical protein | | | |
| group_12866 | dTDP-6-deoxy-L-hexose 3-O-methyltransferase | 2 | F:GO:0008168; P:GO:0032259 | F:methyltransferase activity; P:methylation |
| group_12867 | phytanoyl-CoA dioxygenase | 2 | F:GO:0051213; P:GO:0055114 | F:dioxygenase activity; P:oxidation-reduction process |
| ubiG_3 | class I SAM-dependent methyltransferase | 2 | F:GO:0008168; P:GO:0032259 | F:methyltransferase activity; P:methylation |
| group_12869 | WbqC-like family protein | | | |
| group_12870 | DegT/DnrJ/EryC1/StrS family aminotransferase | 1 | F:GO:0008483 | F:transaminase activity |
| group_12871 | glycosyl transferase | 1 | F:GO:0016740 | F:transferase activity |
| group_12872 | type VI secretion system protein | 1 | C:GO:0016020 | C:membrane |
| group_1425 | type VI secretion protein VgrG | | | |
| group_12873 | hypothetical protein | | | |
| group_12874 | MULTISPECIES: hypothetical protein | 1 | C:GO:0016020 | C:membrane |
| group_6956 | Uncharacterised protein | 2 | C:GO:0016020; C:GO:0016021 | C:membrane; C:integral component of membrane |
| group_4654 | putative Zn-binding protein involved in type VI secretion | | | |
| group_6957 | Uncharacterised protein | 2 | C:GO:0016020; C:GO:0016021 | C:membrane; C:integral component of membrane |
| group_4655 | Uncharacterized conserved protein | | | |
| group_12875 | hypothetical protein N430_01247 | 2 | C:GO:0016020; C:GO:0016021 | C:membrane; C:integral component of membrane |
| group_4656 | putative Zn-binding protein involved in type VI secretion | | | |
| group_12876 | RHS repeat-associated core domain-containing protein | | | |
| group_12877 | ---NA--- | | | |
| group_12878 | hypothetical protein | | | |
| imm_1 | colicin immunity protein | 2 | F:GO:0015643; P:GO:0030153 | F:toxic substance binding; P:bacteriocin immunity |
| col | S-type pyocin | 6 | :0004519; F:GO:0005102; P:GO:0009405; P:GO:0009617; P:GO:0019835; P:GO:00 | ling receptor binding; P:pathogenesis; P:response to bacterium; P:cytolysis; P:nucleic acid phosphodiester bond hydrolysis |
| imm_2 | bacteriocin immunity protein | 3 | F:GO:0005515; F:GO:0015643; P:GO:0030153 | F:protein binding; F:toxic substance binding; P:bacteriocin immunity |
| group_12882 | pyocin | 4 | F:GO:0005488; P:GO:0009987; F:GO:0016787; P:GO:0044419 | binding; P:cellular process; F:hydrolase activity; P:interspecies interaction between organisms |
| group_12883 | hypothetical protein | | | |
| group_12884 | hypothetical protein | | | |
| group_12885 | ---NA--- | | | |
| group_12886 | EamA family transporter | 1 | C:GO:0016021 | C:integral component of membrane |
| group_12887 | Uncharacterised protein | | | |
| hss | homospermidine synthase | 2 | F:GO:0016491; P:GO:0055114 | F:oxidoreductase activity; P:oxidation-reduction process |
| glpQ | glycerophosphodiester phosphodiesterase | 2 | P:GO:0006629; F:GO:0008889 | P:lipid metabolic process; F:glycerophosphodiester phosphodiesterase activity |
| nadB_1 | L-aspartate oxidase | 5 | C:GO:0005737; F:GO:0008734; P:GO:0009435; F:GO:0044318; P:GO:0055114 | idase activity; P:NAD biosynthetic process; F:L-aspartate:fumarate oxidoreductase activity; P:oxidation-reduction process |
| group_12890 | integrase | 3 | F:GO:0003677; P:GO:0006310; P:GO:0015074 | F:DNA binding; P:DNA recombination; P:DNA integration |
| group_12891 | hypothetical protein IEC33019_0915 | 1 | C:GO:0016021 | C:integral component of membrane |
| group_12892 | hypothetical protein | | | |
| group_12893 | hypothetical protein DI560_25180 | | | |
| group_12894 | hypothetical protein | | | |
| group_12895 | MULTISPECIES: hypothetical protein | | | |
| group_12896 | MULTISPECIES: hypothetical protein | | | |
| group_12897 | Rha family transcriptional regulator | | | |

| | | | | |
|---|---|---|---|---|
| group_12898 | Uncharacterised protein | | | |
| group_12899 | leucine-rich repeat-containing protein typical subtype | 2 | F:GO:0004842; P:GO:0016567 | F:ubiquitin-protein transferase activity; P:protein ubiquitination |
| group_12900 | Uncharacterised protein | 1 | C:GO:0016021 | C:integral component of membrane |
| group_12901 | NADH:quinone oxidoreductase | 1 | C:GO:0016021 | C:integral component of membrane |
| group_12902 | electron transporter RnfG | 5 | C:GO:0005886; F:GO:0009055; F:GO:0010181; C:GO:0016021; P:GO:0022900 | e; F:electron transfer activity; F:FMN binding; C:integral component of membrane; P:electron transport chain |
| rnfA | electron transporter RnfA | 1 | C:GO:0016021 | C:integral component of membrane |
| group_12904 | Uncharacterised protein | | | |
| group_12905 | hypothetical protein | | | |
| group_12906 | hypothetical protein | | | |
| group_12907 | integrase | 3 | F:GO:0003677; P:GO:0006310; P:GO:0015074 | F:DNA binding; P:DNA recombination; P:DNA integration |
| group_12908 | integrase | 3 | F:GO:0003677; P:GO:0006310; P:GO:0015074 | F:DNA binding; P:DNA recombination; P:DNA integration |
| group_12909 | Peptidase inhibitor I78 family protein | | | |
| mntH | Mn2+/Fe2+ transporter | 4 | C:GO:0005886; C:GO:0016021; P:GO:0030001; F:GO:0046873 | ie; C:integral component of membrane; P:metal ion transport; F:metal ion transmembrane transporter activity |
| group_12911 | DUF2790 domain-containing protein | | | |
| group_12912 | DUF4034 domain-containing protein | | | |
| group_12913 | hypothetical protein C7318_0443 | | | |
| group_12914 | hypothetical protein | 2 | P:GO:0006629; F:GO:0008081 | P:lipid metabolic process; F:phosphoric diester hydrolase activity |
| group_12915 | OmpA family protein | 1 | C:GO:0016021 | C:integral component of membrane |
| group_12916 | Uncharacterised protein | 1 | C:GO:0016021 | C:integral component of membrane |
| group_12917 | Uncharacterised protein | | | |
| group_12918 | Uncharacterised protein | 1 | C:GO:0016020 | C:membrane |
| imm_3 | bacteriocin immunity protein | 2 | F:GO:0015643; P:GO:0030153 | F:toxic substance binding; P:bacteriocin immunity |
| imm_4 | bacteriocin immunity protein | 2 | F:GO:0015643; P:GO:0030153 | F:toxic substance binding; P:bacteriocin immunity |
| group_12921 | magnesium transporter | 5 | C:GO:0005886; F:GO:0015095; C:GO:0016021; F:GO:0046872; P:GO:1903830 | ısmembrane transporter activity; C:integral component of membrane; F:metal ion binding; P:magnesium ion transmembrane transport |
| group_12922 | insulinase family protein | 2 | F:GO:0003824; F:GO:0046872 | F:catalytic activity; F:metal ion binding |
| group_12923 | insulinase family protein | 2 | F:GO:0016787; F:GO:0046872 | F:hydrolase activity; F:metal ion binding |
| group_12925 | pilus assembly protein PilM | | | |
| group_12926 | fimbrial protein | 1 | C:GO:0016021 | C:integral component of membrane |
| group_12927 | type IV pili biogenesis protein | 1 | C:GO:0016021 | C:integral component of membrane |
| group_12928 | ---NA--- | | | |
| group_12929 | Uncharacterised protein | | | |
| group_12930 | chorismate mutase | 2 | F:GO:0016853; P:GO:0046417 | F:isomerase activity; P:chorismate metabolic process |
| group_12931 | DUF3829 domain-containing protein | | | |
| hscC | molecular chaperone HscC | 1 | F:GO:0005524 | F:ATP binding |
| group_12933 | J domain-containing protein | 1 | C:GO:0016020 | C:membrane |
| group_12934 | CoA ester lyase | 2 | F:GO:0016829; F:GO:0046872 | F:lyase activity; F:metal ion binding |
| group_12935 | TRAP transporter small permease | 1 | C:GO:0016021 | C:integral component of membrane |
| group_12936 | Uncharacterised protein | | | |
| group_12937 | RNA polymerase subunit sigma-70 | 2 | C:GO:0005886; C:GO:0016021 | C:plasma membrane; C:integral component of membrane |
| mrdA_2 | penicillin-binding protein 2 | 8 | :0006508; P:GO:0008360; F:GO:0008658; F:GO:0009002; P:GO:0009252; F:GO:00 | cillin binding; F:serine-type D-Ala-D-Ala carboxypeptidase activity; P:peptidoglycan biosynthetic process; F:transferase activity, transferring glycosyl groups; P:cell wall organization |
| group_12939 | spermidine synthase | 3 | F:GO:0004766; P:GO:0006596; C:GO:0016021 | midine synthase activity; P:polyamine biosynthetic process; C:integral component of membrane |
| group_12940 | arylsulfatase | 1 | F:GO:0004065 | F:arylsulfatase activity |
| group_12941 | haloacid dehalogenase-like hydrolase | 1 | F:GO:0016787 | F:hydrolase activity |
| group_12942 | protein BatD | 1 | C:GO:0016021 | C:integral component of membrane |
| group_12943 | VWA domain-containing protein | 1 | C:GO:0016021 | C:integral component of membrane |
| group_12944 | VWA domain-containing protein | 1 | C:GO:0016021 | C:integral component of membrane |
| group_12945 | DUF4381 domain-containing protein | 1 | C:GO:0016021 | C:integral component of membrane |
| group_12946 | DUF58 domain-containing protein | | | |
| group_12947 | MoxR family ATPase | 2 | F:GO:0005524; F:GO:0016887 | F:ATP binding; F:ATPase activity |
| group_12948 | tetratricopeptide repeat protein | 2 | C:GO:0016021; F:GO:0042802 | C:integral component of membrane; F:identical protein binding |
| group_12949 | transporter | | | |
| group_12950 | Uncharacterised protein | 1 | C:GO:0016021 | C:integral component of membrane |
| group_12951 | DUF3313 domain-containing protein | | | |
| group_12952 | aromatic hydrocarbon degradation protein | | | |
| caiT_2 | BCCT family transporter | 5 | C:GO:0005886; C:GO:0016021; F:GO:0022857; P:GO:0055085; P:GO:0071705 | component of membrane; F:transmembrane transporter activity; P:transmembrane transport; P:nitrogen compound transport |
| bauC_1 | methylmalonate-semialdehyde dehydrogenase (CoA acylating) | 2 | F:GO:0004491; P:GO:0055114 | ıethylmalonate-semialdehyde dehydrogenase (acylating) activity; P:oxidation-reduction process |
| dddD | carnitine dehydratase | 1 | F:GO:0008410 | F:CoA-transferase activity |
| group_12955 | LysR family transcriptional regulator | 3 | F:GO:0003677; F:GO:0003700; P:GO:0006355 | ɔinding; F:DNA-binding transcription factor activity; P:regulation of transcription, DNA-templated |
| acuR | TetR family transcriptional regulator | 1 | F:GO:0003677 | F:DNA binding |
| acuI | alcohol dehydrogenase | 2 | F:GO:0016491; P:GO:0055114 | F:oxidoreductase activity; P:oxidation-reduction process |
| group_12958 | LysR family transcriptional regulator | 3 | F:GO:0003677; F:GO:0003700; P:GO:0006355 | ɔinding; F:DNA-binding transcription factor activity; P:regulation of transcription, DNA-templated |
| group_12959 | amidohydrolase family protein | 1 | F:GO:0016810 | F:hydrolase activity, acting on carbon-nitrogen (but not peptide) bonds |
| group_12960 | EamA-like transporter family protein | 1 | C:GO:0016021 | C:integral component of membrane |
| group_12961 | hypothetical protein | 1 | C:GO:0016021 | C:integral component of membrane |
| group_12962 | GNAT family N-acetyltransferase | 1 | F:GO:0008080 | F:N-acetyltransferase activity |
| group_12963 | NADP oxidoreductase | | | |
| group_12964 | LysR family transcriptional regulator | 3 | F:GO:0003677; F:GO:0003700; P:GO:0006355 | ɔinding; F:DNA-binding transcription factor activity; P:regulation of transcription, DNA-templated |
| group_12965 | TetR family transcriptional regulator | 1 | F:GO:0003677 | F:DNA binding |
| calB_2 | coniferyl aldehyde dehydrogenase | 3 | P:GO:0006081; F:GO:0016620; P:GO:0055114 | :oxidoreductase activity, acting on the aldehyde or oxo group of donors, NAD or NADP as acceptor; P:oxidation-reduction process |
| group_12967 | NAD-dependent dehydratase | 2 | F:GO:0003824; F:GO:0050662 | F:catalytic activity; F:coenzyme binding |
| group_12968 | NAD(P)H-binding protein | 2 | F:GO:0016491; P:GO:0055114 | F:oxidoreductase activity; P:oxidation-reduction process |
| group_12969 | ---NA--- | | | |
| group_12970 | Uncharacterised protein | 1 | C:GO:0016021 | C:integral component of membrane |
| group_12971 | ---NA--- | | | |
| group_112 | Uncharacterised protein | | | |
| group_48 | DUF3077 domain-containing protein | | | |
| group_12972 | entry exclusion lipoprotein TrbK | | | |
| group_113 | Uncharacterised protein | | | |
| group_53 | DUF3077 domain-containing protein | | | |
| group_12973 | FAD-binding oxidoreductase | 3 | F:GO:0016899; P:GO:0055114; F:GO:0071949 | vity, acting on the CH-OH group of donors, oxygen as acceptor; P:oxidation-reduction process; F:FAD binding |
| group_12974 | TonB-dependent receptor | 2 | C:GO:0009279; C:GO:0016021 | C:cell outer membrane; C:integral component of membrane |
| group_12975 | DUF1302 domain-containing protein | | | |
| group_12976 | thioesterase | 1 | F:GO:0016787 | F:hydrolase activity |
| group_12977 | DUF202 domain-containing protein | 1 | C:GO:0016021 | C:integral component of membrane |
| group_12978 | DUF202 domain-containing protein | 1 | C:GO:0016021 | C:integral component of membrane |
| group_12979 | DUF1329 domain-containing protein | | | |
| group_12980 | cell division protein | 2 | C:GO:0016021; P:GO:0051301 | C:integral component of membrane; P:cell division |
| group_12981 | TIGR01777 family protein | 4 | F:GO:0003824; C:GO:0016021; F:GO:0050662; P:GO:0051301 | :catalytic activity; C:integral component of membrane; F:coenzyme binding; P:cell division |
| group_12982 | alpha/beta hydrolase | 4 | F:GO:0005524; F:GO:0016301; P:GO:0016310; F:GO:0016787 | F:ATP binding; F:kinase activity; P:phosphorylation; F:hydrolase activity |
| group_12983 | MFS transporter | 2 | C:GO:0016021; P:GO:0055085 | C:integral component of membrane; P:transmembrane transport |
| group_12984 | LysR family transcriptional regulator | 3 | F:GO:0003677; F:GO:0003700; P:GO:0006355 | ɔinding; F:DNA-binding transcription factor activity; P:regulation of transcription, DNA-templated |
| group_12985 | DUF2878 domain-containing protein | 1 | C:GO:0016021 | C:integral component of membrane |
| group_12986 | periplasmic protein | 1 | F:GO:0016872 | F:intramolecular lyase activity |
| group_12987 | UDP-glucose/GDP-mannose dehydrogenase family protein | 4 | P:GO:0000271; F:GO:0003979; F:GO:0051287; P:GO:0055114 | iosynthetic process; F:UDP-glucose 6-dehydrogenase activity; F:NAD binding; P:oxidation-reduction process |
| group_12988 | polysaccharide lyase family 7 protein | 1 | F:GO:0016829 | F:lyase activity |
| group_12989 | hypothetical protein | | | |
| group_12990 | hypothetical protein | | | |
| group_3050 | hypothetical protein | | | |
| group_12991 | peptidase S24 | 5 | F:GO:0003677; P:GO:0006281; P:GO:0006355; P:GO:0009432; F:GO:0016787 | ıng; P:DNA repair; P:regulation of transcription, DNA-templated; P:SOS response; F:hydrolase activity |
| group_12992 | Putative SOS response-associated peptidase YedK | 2 | P:GO:0006508; F:GO:0008233 | P:proteolysis; F:peptidase activity |
| group_12993 | GGDEF domain-containing protein | 2 | C:GO:0016021; F:GO:0052621 | C:integral component of membrane; F:diguanylate cyclase activity |
| group_12994 | ---NA--- | | | |
| group_12995 | Uncharacterised protein | | | |
| group_12996 | hypothetical protein | | | |
| group_12997 | type VI secretion system secreted protein VgrG | 2 | C:GO:0016020; C:GO:0016021 | C:membrane; C:integral component of membrane |
| group_12998 | signal peptide protein | | | |
| group_12999 | DUF2169 domain-containing protein | | | |
| group_13000 | DUF4150 domain-containing protein | | | |
| group_13001 | filamentous hemagglutinin | | | |
| group_13002 | GAD-like domain protein | | | |
| group_13003 | nuclease | | | |
| group_2243 | type IV secretion protein Rhs | 2 | C:GO:0016021; F:GO:0016787 | C:integral component of membrane; F:hydrolase activity |
| group_13004 | Uncharacterised protein | 2 | F:GO:0016158; P:GO:0016311 | F:3-phytase activity; P:dephosphorylation |
| group_13005 | hypothetical protein | | | |
| group_2244 | type IV secretion protein Rhs | 2 | C:GO:0016021; F:GO:0016787 | C:integral component of membrane; F:hydrolase activity |
| group_13006 | ---NA--- | | | |
| group_13008 | peptide transporter | 2 | C:GO:0016021; P:GO:0055085 | C:integral component of membrane; P:transmembrane transport |
| group_109 | Uncharacterised protein | | | |
| group_47 | DUF3077 domain-containing protein | | | |
| group_13009 | NUDIX domain-containing protein | 1 | F:GO:0016787 | F:hydrolase activity |
| group_13010 | Uncharacterised protein | 1 | C:GO:0016021 | C:integral component of membrane |
| group_13011 | helicase | 3 | F:GO:0003676; F:GO:0004386; F:GO:0005524 | F:nucleic acid binding; F:helicase activity; F:ATP binding |
| group_13012 | transposase | 4 | F:GO:0003677; F:GO:0004803; P:GO:0006313; P:GO:0015074 | ':DNA binding; F:transposase activity; P:transposition, DNA-mediated; P:DNA integration |
| group_110 | Uncharacterised protein | | | |
| group_46 | DUF3077 domain-containing protein | | | |
| group_13013 | camphor resistance protein CrcB | 4 | C:GO:0005887; F:GO:0015103; P:GO:0015698; P:GO:0034220 | nembrane; F:inorganic anion transmembrane transporter activity; P:inorganic anion transport; P:ion transmembrane transport |
| group_13014 | type I secretion C-terminal target domain-containing protein | 3 | F:GO:0005509; P:GO:0007229; C:GO:0016021 | lcium ion binding; P:integrin-mediated signaling pathway; C:integral component of membrane |
| group_13015 | hypothetical protein | | | |
| group_13016 | Uncharacterised protein | | | |
| group_13017 | Uncharacterised protein | | | |
| group_13018 | RNA-directed DNA polymerase | 1 | F:GO:0016779 | F:nucleotidyltransferase activity |
| group_13019 | hypothetical protein | 2 | C:GO:0016020; C:GO:0016021 | C:membrane; C:integral component of membrane |
| group_13020 | HNH endonuclease | | | |
| group_3062 | integrase | 3 | F:GO:0003677; P:GO:0006310; P:GO:0015074 | F:DNA binding; P:DNA recombination; P:DNA integration |
| group_3063 | integrase | 3 | F:GO:0003677; P:GO:0006310; P:GO:0015074 | F:DNA binding; P:DNA recombination; P:DNA integration |
| group_13021 | type II toxin-antitoxin system RelE/ParE family toxin | | | |
| group_13023 | ---NA--- | | | |
| group_13024 | Uncharacterised protein | | | |
| group_13025 | hypothetical protein | | | |
| group_13026 | peptidase M72 family protein | 2 | P:GO:0006508; F:GO:0008237 | P:proteolysis; F:metallopeptidase activity |
| group_13027 | RHS repeat-associated core domain-containing protein | 2 | C:GO:0016020; C:GO:0016021 | C:membrane; C:integral component of membrane |
| group_13028 | ---NA--- | | | |
| group_13029 | protein of unknown function | | | |
| group_13030 | RHS repeat-associated core domain-containing protein | | | |
| group_57 | DUF3077 domain-containing protein | | | |
| group_13031 | RHS repeat-associated core domain-containing protein | | | |
| group_13032 | MULTISPECIES: hypothetical protein | | | |
| group_13033 | RHS repeat-associated core domain-containing protein | | | |
| group_13034 | cell envelope integrity protein CreD | 1 | C:GO:0016021 | C:integral component of membrane |
| group_13035 | type III effector 1 | | | |
| group_13036 | Uncharacterised protein | 1 | C:GO:0016021 | C:integral component of membrane |
| group_13037 | ComF family protein | 2 | P:GO:0009116; F:GO:0016757 | P:nucleoside metabolic process; F:transferase activity, transferring glycosyl groups |
| group_13038 | pyrroloquinoline quinone biosynthesis protein PqqF | 4 | F:GO:0004222; P:GO:0006508; F:GO:0008270; P:GO:0018189 | opeptidase activity; P:proteolysis; F:zinc ion binding; P:pyrroloquinoline quinone biosynthetic process |
| group_13039 | ATP-binding protein | 1 | F:GO:0005524 | F:ATP binding |
| group_20 | transposase | 3 | F:GO:0003677; F:GO:0004803; P:GO:0006313 | F:DNA binding; F:transposase activity; P:transposition, DNA-mediated |
| group_19 | IS3 family transposase | 3 | F:GO:0003676; P:GO:0015074; P:GO:0032196 | F:nucleic acid binding; P:DNA integration; P:transposition |
| group_13040 | Abi family protein | | | |
| group_13041 | Uncharacterised protein | | | |
| group_13042 | P-type DNA transfer protein VirB5 | | | |
| group_13043 | Uncharacterised protein | | | |
| group_13044 | ribonuclease T2 | 3 | F:GO:0003723; F:GO:0033897; P:GO:0090502 | A binding; F:ribonuclease T2 activity; P:RNA phosphodiester bond hydrolysis, endonucleolytic |
| group_13045 | MFS transporter | 2 | C:GO:0016021; P:GO:0055085 | C:integral component of membrane; P:transmembrane transport |
| group_13046 | hybrid sensor histidine kinase/response regulator | 3 | F:GO:0000155; P:GO:0000160; P:GO:0023014 | ɔr kinase activity; P:phosphorelay signal transduction system; P:signal transduction by protein phosphorylation |
| group_13047 | hybrid sensor histidine kinase/response regulator | 4 | F:GO:0000155; P:GO:0000160; F:GO:0000166; P:GO:0023014 | tivity; P:phosphorelay signal transduction system; F:nucleotide binding; P:signal transduction by protein phosphorylation |
| group_13048 | CSS domain containing protein | 1 | C:GO:0016021 | C:integral component of membrane |
| group_13049 | diguanylate cyclase | 1 | C:GO:0016021 | C:integral component of membrane |
| group_13050 | Uncharacterised protein | | | |
| group_13052 | AraC family transcriptional regulator | 3 | F:GO:0003700; P:GO:0006355; F:GO:0043565 | ranscription factor activity; P:regulation of transcription, DNA-templated; F:sequence-specific DNA binding |
| group_4930 | NAD(P)/FAD-dependent oxidoreductase | 2 | F:GO:0016491; P:GO:0055114 | F:oxidoreductase activity; P:oxidation-reduction process |
| group_13053 | Uncharacterised protein | | | |
| group_13054 | hypothetical protein | | | |
| group_13055 | methyl-accepting chemotaxis protein | 4 | F:GO:0004888; P:GO:0006935; P:GO:0007165; C:GO:0016021 | ane signaling receptor activity; P:chemotaxis; P:signal transduction; C:integral component of membrane |
| group_13056 | hybrid sensor histidine kinase/response regulator | 3 | F:GO:0000155; P:GO:0000160; P:GO:0023014 | ɔr kinase activity; P:phosphorelay signal transduction system; P:signal transduction by protein phosphorylation |
| group_13057 | ArtI protein | | | |
| group_13058 | ferrous iron transport protein A | 1 | F:GO:0046914 | F:transition metal ion binding |
| group_13059 | ferrous iron transporter B | 5 | F:GO:0005525; F:GO:0015093; C:GO:0016021; P:GO:0034755; P:GO:0072511 | ane transporter activity; C:integral component of membrane; P:iron ion transmembrane transport; P:divalent inorganic cation transport |
| group_13060 | stress-induced protein | | | |
| group_4952 | WYL domain-containing protein | | | |
| group_13061 | N-acetyltransferase | 2 | C:GO:0005840; F:GO:0008080 | C:ribosome; F:N-acetyltransferase activity |
| group_13062 | Uncharacterised protein | | | |
| group_13063 | DUF2063 domain-containing protein | 1 | F:GO:0003677 | F:DNA binding |
| group_13064 | hypothetical protein | | | |
| ydjE | MFS transporter | 3 | C:GO:0016021; F:GO:0022857; P:GO:0055085 | egral component of membrane; F:transmembrane transporter activity; P:transmembrane transport |
| group_13066 | nuclear transport factor 2 family protein | | | |
| group_13067 | ethanolamine utilization protein EutN | 1 | P:GO:0006865 | P:amino acid transport |
| group_13068 | branched-chain amino acid ABC transporter permease | 4 | C:GO:0005886; C:GO:0016021; F:GO:0022857; P:GO:0055085 | rane; C:integral component of membrane; F:transmembrane transporter activity; P:transmembrane transport |
| btuD_8 | ABC transporter | 6 | 0005524; C:GO:0005886; C:GO:0016021; F:GO:0016887; F:GO:0022857; P:GO:00 | ne; C:integral component of membrane; F:ATPase activity; F:transmembrane transporter activity; P:transmembrane transport |
| codA_2 | amidohydrolase | 2 | F:GO:0004131; F:GO:0102480 | F:cytosine deaminase activity; F:5-fluorocytosine deaminase activity |
| group_13071 | GntR family transcriptional regulator | 3 | F:GO:0003677; F:GO:0003700; P:GO:0006355 | ɔinding; F:DNA-binding transcription factor activity; P:regulation of transcription, DNA-templated |
| group_13072 | Gamma-glutamyltransferase | 2 | F:GO:0102953; F:GO:0103068 | cin A gamma-glutamyl transpeptidase activity; F:leukotriene C4 gamma-glutamyl transferase activity |
| group_13073 | LuxR family transcriptional regulator | 2 | F:GO:0003677; P:GO:0006355 | F:DNA binding; P:regulation of transcription, DNA-templated |
| group_13074 | membrane protein, TIGR04086 family | 1 | C:GO:0016021 | C:integral component of membrane |
| group_13075 | CsbD family protein | | | |
| group_13076 | TIGR02450 family Trp-rich protein | | | |
| group_13077 | hypothetical protein | | | |
| group_13078 | GABA permease | 4 | P:GO:0006865; C:GO:0016021; F:GO:0022857; P:GO:0055085 | sport; C:integral component of membrane; F:transmembrane transporter activity; P:transmembrane transport |
| group_13079 | cupin | | | |

| | | | | |
|---|---|---|---|---|
| group_13080 | LysR family transcriptional regulator | 3 | F:GO:0003677; F:GO:0003700; P:GO:0006355 | pinding; F:DNA-binding transcription factor activity; P:regulation of transcription, DNA-templated |
| group_13081 | aminotransferase class III-fold pyridoxal phosphate-dependent enzyme | 3 | F:GO:0008483; F:GO:0030170; F:GO:0042286 | nase activity; F:pyridoxal phosphate binding; F:glutamate-1-semialdehyde 2,1-aminomutase activity |
| group_13082 | phosphodiesterase | 2 | F:GO:0016787; F:GO:0046872 | F:hydrolase activity; F:metal ion binding |
| group_13083 | phage protein | | | |
| group_13084 | phage tail protein I | | | |
| group_13085 | phage tail protein | | | |
| group_13086 | pyocin R2_PP, tail length determination protein | | | |
| group_13087 | phage tail protein | | | |
| group_13088 | phage tail protein | | | |
| group_13089 | late control protein | | | |
| group_13090 | lysis protein | 2 | P:GO:0019076; P:GO:0019835 | P:viral release from host cell; P:cytolysis |
| group_13091 | ---NA--- | | | |
| talB | transaldolase | 4 | F:GO:0004801; C:GO:0005737; P:GO:0005975; P:GO:0006098 | raldehyde-3-phosphate glyceronetransferase activity; C:cytoplasm; P:carbohydrate metabolic process; P:pentose-phosphate shunt |
| xylB_1 | FGGY-family carbohydrate kinase | 2 | F:GO:0004856; P:GO:0046835 | F:xylulokinase activity; P:carbohydrate phosphorylation |
| purR_1 | LacI family transcriptional regulator | 2 | F:GO:0003677; P:GO:0006355 | F:DNA binding; P:regulation of transcription, DNA-templated |
| rbtD | NAD(P)-dependent oxidoreductase | 2 | F:GO:0050255; P:GO:0055114 | F:ribitol 2-dehydrogenase activity; P:oxidation-reduction process |
| group_13096 | 5-carboxymethyl-2-hydroxymuconate isomerase | 2 | F:GO:0008704; P:GO:0019439 | boxymethyl-2-hydroxymuconate delta-isomerase activity; P:aromatic compound catabolic process |
| group_13097 | DUF2892 domain-containing protein | | | |
| group_13098 | ---NA--- | | | |
| group_13099 | DUF4265 domain-containing protein | | | |
| quiC | 3-dehydroshikimate dehydratase | 1 | F:GO:0046565 | F:3-dehydroshikimate dehydratase activity |
| hisC_6 | GntR family transcriptional regulator | 5 | F:GO:0003677; F:GO:0003700; P:GO:0006355; F:GO:0008483; F:GO:0030170 | scription factor activity; P:regulation of transcription, DNA-templated; F:transaminase activity; F:pyridoxal phosphate binding |
| group_13102 | transporter | 2 | C:GO:0016021; P:GO:0055085 | C:integral component of membrane; P:transmembrane transport |
| group_13103 | transcriptional regulator | 1 | F:GO:0003677 | F:DNA binding |
| group_13104 | hypothetical protein | | | |
| group_13105 | hypothetical protein | | | |
| group_13106 | hypothetical protein CBL13_02901 | | | |
| proA_2 | RraA family protein | 3 | F:GO:0008168; P:GO:0032259; F:GO:0047443 | ethyltransferase activity; P:methylation; F:4-hydroxy-4-methyl-2-oxoglutarate aldolase activity |
| group_13108 | hydroxyacid dehydrogenase | 4 | F:GO:0004617; F:GO:0016618; F:GO:0051287; P:GO:0055114 | dehydrogenase activity; F:hydroxypyruvate reductase activity; F:NAD binding; P:oxidation-reduction process |
| proP_5 | MFS transporter | 3 | C:GO:0016021; F:GO:0022857; P:GO:0055085 | egral component of membrane; F:transmembrane transporter activity; P:transmembrane transport |
| group_13110 | helix-turn-helix domain-containing protein | 2 | F:GO:0003677; P:GO:0006355 | F:DNA binding; P:regulation of transcription, DNA-templated |
| group_13111 | methyl-accepting chemotaxis protein | 5 | F:GO:0004888; P:GO:0006935; P:GO:0007165; C:GO:0016020; C:GO:0016021 | naling receptor activity; P:chemotaxis; P:signal transduction; C:membrane; C:integral component of membrane |
| group_13112 | hypothetical protein | | | |
| xylC | benzaldehyde dehydrogenase | 2 | F:GO:0018479; P:GO:0055114 | F:benzaldehyde dehydrogenase (NAD+) activity; P:oxidation-reduction process |
| acoR_4 | sigma-54-dependent Fis family transcriptional regulator | 2 | F:GO:0000166; F:GO:0003677 | F:nucleotide binding; F:DNA binding |
| group_13115 | ABC transporter ATP-binding protein | 2 | F:GO:0000166; C:GO:0016020 | F:nucleotide binding; C:membrane |
| group_13116 | FAD-dependent oxidoreductase | 1 | F:GO:0071949 | F:FAD binding |
| group_13117 | quinohemoprotein amine dehydrogenase subunit alpha | 1 | F:GO:0005488 | F:binding |
| group_13118 | quinohemoprotein amine dehydrogenase maturation protein | 3 | F:GO:0003824; F:GO:0046872; F:GO:0051539 | F:catalytic activity; F:metal ion binding; F:4 iron, 4 sulfur cluster binding |
| group_13119 | quinohemoprotein amine dehydrogenase subunit beta | | | |
| apr | subtilisin | 2 | F:GO:0004252; P:GO:0006508 | F:serine-type endopeptidase activity; P:proteolysis |
| hemL_3 | glutamate-1-semialdehyde 2,1-aminomutase | 2 | F:GO:0008483; F:GO:0030170 | F:transaminase activity; F:pyridoxal phosphate binding |
| group_13122 | LysR family transcriptional regulator | 3 | F:GO:0003677; F:GO:0003700; P:GO:0006355 | pinding; F:DNA-binding transcription factor activity; P:regulation of transcription, DNA-templated |
| pchA | aldehyde dehydrogenase | 2 | F:GO:0016620; P:GO:0055114 | ivity, acting on the aldehyde or oxo group of donors, NAD or NADP as acceptor; P:oxidation-reduction process |
| group_13124 | APC family permease | 3 | C:GO:0016021; F:GO:0022857; P:GO:0055085 | egral component of membrane; F:transmembrane transporter activity; P:transmembrane transport |
| group_13125 | hypothetical protein | | | |
| group_13126 | APC family permease | 3 | C:GO:0016021; F:GO:0022857; P:GO:0055085 | egral component of membrane; F:transmembrane transporter activity; P:transmembrane transport |
| group_13127 | aspartate aminotransferase family protein | 2 | F:GO:0008483; F:GO:0030170 | F:transaminase activity; F:pyridoxal phosphate binding |
| group_13128 | LysR family transcriptional regulator | 3 | F:GO:0003677; F:GO:0003700; P:GO:0006355 | pinding; F:DNA-binding transcription factor activity; P:regulation of transcription, DNA-templated |
| pht2_1 | oxidoreductase | 8 | 0009055; C:GO:0016021; F:GO:0018620; P:GO:0022900; P:GO:0032259; F:GO:00 | mponent of membrane; F:phthalate 4,5-dioxygenase activity; P:electron transport chain; P:methylation; F:metal ion binding; F:2 iron, 2 sulfur cluster binding |
| group_13130 | MarR family transcriptional regulator | 3 | F:GO:0003677; F:GO:0003700; P:GO:0006355 | pinding; F:DNA-binding transcription factor activity; P:regulation of transcription, DNA-templated |
| group_13131 | TetR/AcrR family transcriptional regulator | 1 | F:GO:0003677 | F:DNA binding |
| group_13132 | MBL fold metallo-hydrolase | 1 | F:GO:0016787 | F:hydrolase activity |
| group_13133 | ester cyclase | 1 | P:GO:0030638 | P:polyketide metabolic process |
| group_13134 | cyanate hydratase | 3 | F:GO:0003677; F:GO:0008824; P:GO:0009439 | F:DNA binding; F:cyanate hydratase activity; P:cyanate metabolic process |
| mcpU_3 | methyl-accepting chemotaxis protein | 1 | C:GO:0016020 | C:membrane |
| group_13136 | hypothetical protein | | | |
| group_13137 | hypothetical protein PMI38_00840 | | | |
| group_13138 | DUF2188 domain-containing protein | | | |
| group_13139 | response regulator | 2 | P:GO:0000160; F:GO:0003677 | P:phosphorelay signal transduction system; F:DNA binding |
| group_13140 | response regulator | 1 | P:GO:0000160 | P:phosphorelay signal transduction system |
| group_13141 | low affinity iron permease family protein | 2 | C:GO:0016021; P:GO:0055085 | C:integral component of membrane; P:transmembrane transport |
| group_13142 | peptidase M42 | 3 | F:GO:0004177; P:GO:0006508; F:GO:0046872 | F:aminopeptidase activity; P:proteolysis; F:metal ion binding |
| group_13143 | DUF421 domain-containing protein | 1 | C:GO:0016021 | C:integral component of membrane |
| group_13144 | hypothetical protein | 1 | C:GO:0016021 | C:integral component of membrane |
| group_4988 | CinA family protein | | | |
| group_13145 | ---NA--- | | | |
| group_4456 | NAD(P)-dependent oxidoreductase | 2 | F:GO:0047936; P:GO:0055114 | F:glucose 1-dehydrogenase [NAD(P)] activity; P:oxidation-reduction process |
| group_13146 | hypothetical protein WR25_00526 | 2 | C:GO:0016020; C:GO:0016021 | C:membrane; C:integral component of membrane |
| group_13147 | hypothetical protein | | | |
| group_13148 | hypothetical protein | | | |
| yciF | ferritin-like domain-containing protein | | | |
| group_3051 | MULTISPECIES: hypothetical protein | | | |
| group_13150 | ShlB/FhaC/HecB family hemolysin secretion/activation protein | | | |
| glmU_1 | transferase | 2 | F:GO:0003977; F:GO:0019134 | -acetylglucosamine diphosphorylase activity; F:glucosamine-1-phosphate N-acetyltransferase activity |
| group_13152 | cytochrome-c oxidase | 2 | F:GO:0016627; P:GO:0055114 | oxidoreductase activity, acting on the CH-CH group of donors; P:oxidation-reduction process |
| group_13153 | ---NA--- | | | |
| group_13154 | hypothetical protein | | | |
| group_13155 | molecular chaperone Tir | | | |
| group_13157 | RHS repeat-associated core domain-containing protein | 2 | C:GO:0016020; C:GO:0016021 | C:membrane; C:integral component of membrane |
| group_13158 | peptide ABC transporter substrate-binding protein | 1 | C:GO:0016021 | C:integral component of membrane |
| group_13159 | TonB-dependent siderophore receptor | 4 | C:GO:0009279; F:GO:0015344; F:GO:0038023; P:GO:0044718 | erophore uptake transmembrane transporter activity; F:signaling receptor activity; P:siderophore transmembrane transport |
| group_13160 | Fic family protein | | | |
| group_3069 | DUF2955 domain-containing protein | 1 | C:GO:0016021 | C:integral component of membrane |
| group_3070 | DUF2955 domain-containing protein | 1 | C:GO:0016021 | C:integral component of membrane |
| group_13161 | MULTISPECIES: hypothetical protein | | | |
| group_13162 | DEAD/DEAH box helicase | 4 | F:GO:0003677; F:GO:0004386; F:GO:0005524; P:GO:0006139 | pinding; F:helicase activity; F:ATP binding; P:nucleobase-containing compound metabolic process |
| group_13163 | lipocalin-like domain-containing protein | | | |
| bacC | SDR family oxidoreductase | 4 | F:GO:0004316; P:GO:0055114; F:GO:0102131; F:GO:0102132 | tivity; P:oxidation-reduction process; F:3-oxo-glutaryl-[acp] methyl ester reductase activity; F:3-oxo-pimeloyl-[acp] methyl ester reductase activity |
| group_13165 | LysR family transcriptional regulator | 3 | F:GO:0003677; F:GO:0003700; P:GO:0006355 | pinding; F:DNA-binding transcription factor activity; P:regulation of transcription, DNA-templated |
| group_13166 | alcohol dehydrogenase | 3 | C:GO:0016020; F:GO:0016491; P:GO:0055114 | C:membrane; F:oxidoreductase activity; P:oxidation-reduction process |
| group_13167 | AraC family transcriptional regulator | 3 | F:GO:0003700; P:GO:0006355; F:GO:0043565 | ranscription factor activity; P:regulation of transcription, DNA-templated; F:sequence-specific DNA binding |
| group_13168 | hypothetical protein | | | |
| group_13169 | multidrug transporter | 5 | C:GO:0005886; C:GO:0009279; F:GO:0015562; C:GO:0016021; P:GO:0055085 | er membrane; F:efflux transmembrane transporter activity; C:integral component of membrane; P:transmembrane transport |
| group_13170 | multidrug efflux RND transporter permease subunit | 5 | C:GO:0005886; F:GO:0015562; C:GO:0016021; P:GO:0042908; P:GO:0055085 | ransmembrane transporter activity; C:integral component of membrane; P:xenobiotic transport; P:transmembrane transport |
| srpA_2 | efflux RND transporter periplasmic adaptor subunit | 3 | C:GO:0005886; F:GO:0022857; P:GO:0055085 | C:plasma membrane; F:transmembrane transporter activity; P:transmembrane transport |
| ttgV | transcriptional regulator | 2 | F:GO:0003677; P:GO:0006355 | F:DNA binding; P:regulation of transcription, DNA-templated |
| srpR | TetR family transcriptional regulator | 1 | F:GO:0003677 | F:DNA binding |
| group_1939 | acetolactate synthase large subunit | 3 | F:GO:0000287; F:GO:0003984; F:GO:0030976 | magnesium ion binding; F:acetolactate synthase activity; F:thiamine pyrophosphate binding |
| tfdB | 2,4-dichlorophenol 6-monooxygenase | 3 | F:GO:0004497; P:GO:0055114; F:GO:0071949 | F:monooxygenase activity; P:oxidation-reduction process; F:FAD binding |
| group_13175 | outer membrane porin, OprD family | 1 | C:GO:0016021 | C:integral component of membrane |
| mhpT_1 | 3-(3-hydroxy-phenyl)propionate transporter MhpT | 3 | C:GO:0016021; F:GO:0022857; P:GO:0055085 | egral component of membrane; F:transmembrane transporter activity; P:transmembrane transport |
| group_13177 | 4-hydroxy-2-oxovalerate aldolase | 3 | F:GO:0008701; P:GO:0019439; F:GO:0030145 | xy-2-oxovalerate aldolase activity; P:aromatic compound catabolic process; F:manganese ion binding |
| mhpF | acetaldehyde dehydrogenase | 4 | F:GO:0008774; P:GO:0019439; F:GO:0051287; P:GO:0055114 | enase (acetylating) activity; P:aromatic compound catabolic process; F:NAD binding; P:oxidation-reduction process |
| mhpD | 2-keto-4-pentenoate hydratase | 3 | F:GO:0008684; P:GO:0019380; F:GO:0030145 | pent-4-enoate hydratase activity; P:3-phenylpropionate catabolic process; F:manganese ion binding |
| mhpC | 2-hydroxy-6-oxo-6-phenylhexa-2,4-dienoate hydrolase | 5 | C:GO:0005737; F:GO:0018771; P:GO:0019380; F:GO:0042803; F:GO:0052823 | ase activity; P:3-phenylpropionate catabolic process; F:protein homodimerization activity; F:2-hydroxy-6-oxonona-2,4,7-trienedioate hydrolase activity |
| mhpB | 3-(2,3-dihydroxyphenyl)propionate dioxygenase | 4 | F:GO:0008198; P:GO:0019380; F:GO:0047070; P:GO:0055114 | -phenylpropionate catabolic process; F:3-carboxyethylcatechol 2,3-dioxygenase activity; P:oxidation-reduction process |
| group_13182 | GntR family transcriptional regulator | 3 | F:GO:0003677; F:GO:0003700; P:GO:0006355 | pinding; F:DNA-binding transcription factor activity; P:regulation of transcription, DNA-templated |
| group_13183 | transporter | | | |
| group_13184 | cupin domain-containing protein | | | |
| adhT | alcohol dehydrogenase | 3 | C:GO:0016021; F:GO:0016491; P:GO:0055114 | :integral component of membrane; F:oxidoreductase activity; P:oxidation-reduction process |
| lspA_1 | lipoprotein signal peptidase | 4 | F:GO:0004190; C:GO:0005886; P:GO:0006508; C:GO:0016021 | -type endopeptidase activity; C:plasma membrane; P:proteolysis; C:integral component of membrane |
| group_13187 | peptidase S8 | 2 | F:GO:0004252; P:GO:0006508 | F:serine-type endopeptidase activity; P:proteolysis |
| group_13188 | peptidase S8 and S53, subtilisin, kexin, sedolisin | 2 | F:GO:0004252; P:GO:0006508 | F:serine-type endopeptidase activity; P:proteolysis |
| group_13189 | AAA family ATPase | 1 | F:GO:0005524 | F:ATP binding |
| group_13190 | diguanylate cyclase | 3 | F:GO:0003677; F:GO:0004386; F:GO:0005524 | F:DNA binding; F:helicase activity; F:ATP binding |
| group_13191 | hypothetical protein | | | |
| group_13192 | XRE family transcriptional regulator | 1 | F:GO:0003677 | F:DNA binding |
| group_13193 | Uncharacterised protein | | | |
| group_13194 | integrase family protein | 3 | F:GO:0003677; P:GO:0006310; P:GO:0015074 | F:DNA binding; P:DNA recombination; P:DNA integration |
| group_13195 | site-specific integrase | 3 | F:GO:0003677; P:GO:0006310; P:GO:0015074 | F:DNA binding; P:DNA recombination; P:DNA integration |
| group_13196 | integrase | 3 | F:GO:0003677; P:GO:0006310; P:GO:0015074 | F:DNA binding; P:DNA recombination; P:DNA integration |
| group_13197 | ---NA--- | | | |
| group_13198 | short-chain dehydrogenase | 2 | F:GO:0018509; P:GO:0055114 | F:cis-2,3-dihydrobiphenyl-2,3-diol dehydrogenase activity; P:oxidation-reduction process |
| group_13199 | hypothetical protein | | | |
| group_13200 | hypothetical protein | | | |
| group_13201 | methyl-accepting chemotaxis protein | 2 | P:GO:0007165; C:GO:0016021 | P:signal transduction; C:integral component of membrane |
| group_13202 | TonB-dependent siderophore receptor | 4 | C:GO:0009279; F:GO:0015344; F:GO:0038023; P:GO:0044718 | erophore uptake transmembrane transporter activity; F:signaling receptor activity; P:siderophore transmembrane transport |
| group_13203 | LysR family transcriptional regulator | 3 | F:GO:0003677; F:GO:0003700; P:GO:0006355 | pinding; F:DNA-binding transcription factor activity; P:regulation of transcription, DNA-templated |
| group_13204 | Uncharacterised protein | | | |
| group_13205 | peptidase S8 | 2 | F:GO:0004252; P:GO:0006508 | F:serine-type endopeptidase activity; P:proteolysis |
| group_3072 | TonB-dependent siderophore receptor | 5 | C:GO:0009279; F:GO:0015344; C:GO:0016021; F:GO:0038023; P:GO:0044718 | nsmembrane transporter activity; C:integral component of membrane; F:signaling receptor activity; P:siderophore transmembrane transport |
| group_13206 | sugar ABC transporter substrate-binding protein | | | |
| group_13207 | TetR family transcriptional regulator | 1 | F:GO:0003677 | F:DNA binding |
| group_13208 | amidohydrolase | 1 | F:GO:0016810 | F:hydrolase activity, acting on carbon-nitrogen (but not peptide) bonds |
| group_13209 | MFS transporter | 3 | C:GO:0016021; F:GO:0022857; P:GO:0055085 | egral component of membrane; F:transmembrane transporter activity; P:transmembrane transport |
| group_13210 | DUF1737 domain-containing protein | | | |
| hchA_1 | type 1 glutamine amidotransferase domain-containing protein | 4 | P:GO:0006508; P:GO:0006541; F:GO:0008233; F:GO:0016740 | P:proteolysis; P:glutamine metabolic process; F:peptidase activity; F:transferase activity |
| group_13212 | LysR family transcriptional regulator | 3 | F:GO:0003677; F:GO:0003700; P:GO:0006355 | pinding; F:DNA-binding transcription factor activity; P:regulation of transcription, DNA-templated |
| pucK | permease | 4 | P:GO:0015851; C:GO:0016021; F:GO:0022857; P:GO:0055085 | sport; C:integral component of membrane; F:transmembrane transporter activity; P:transmembrane transport |
| group_13214 | Cytosine/adenosine deaminase | 1 | F:GO:0008892 | F:guanine deaminase activity |
| group_13215 | aldo/keto reductase | 2 | F:GO:0016491; P:GO:0055114 | F:oxidoreductase activity; P:oxidation-reduction process |
| group_13216 | Uncharacterised protein | | | |
| group_13217 | DUF393 domain-containing protein | | | |
| ttuB_3 | MFS transporter | 2 | C:GO:0016021; P:GO:0055085 | C:integral component of membrane; P:transmembrane transport |
| group_13219 | FadR family transcriptional regulator | 3 | F:GO:0003677; F:GO:0003700; P:GO:0006355 | pinding; F:DNA-binding transcription factor activity; P:regulation of transcription, DNA-templated |
| group_13220 | hypothetical protein | | | |
| group_13221 | hypothetical protein O164_12795 | | | |
| group_13222 | hypothetical protein EOP15_00105 | | | |
| group_13223 | putative lipoprotein | | | |
| group_13224 | MULTISPECIES: hypothetical protein | 1 | C:GO:0016021 | C:integral component of membrane |
| group_13225 | hybrid sensor histidine kinase/response regulator | 3 | F:GO:0000155; P:GO:0000160; P:GO:0023014 | or kinase activity; P:phosphorelay signal transduction system; P:signal transduction by protein phosphorylation |
| xerC_6 | Integrase | 3 | F:GO:0003677; P:GO:0006310; P:GO:0015074 | F:DNA binding; P:DNA recombination; P:DNA integration |
| pinR | resolvase | 4 | F:GO:0000150; F:GO:0003677; P:GO:0006310; P:GO:0006355 | nase activity; F:DNA binding; P:DNA recombination; P:regulation of transcription, DNA-templated |
| group_13228 | hypothetical protein | | | |
| group_13229 | ---NA--- | | | |
| group_13230 | KilA-N domain-containing protein | | | |
| group_13231 | hypothetical protein I1A_000717 | | | |
| group_13232 | hypothetical protein | | | |
| group_13233 | protein TadG | 1 | C:GO:0016021 | C:integral component of membrane |
| group_13234 | PAS domain-containing sensor histidine kinase | 5 | F:GO:0000155; P:GO:0000160; F:GO:0005524; P:GO:0018106; P:GO:0023014 | orelay signal transduction system; F:ATP binding; P:peptidyl-histidine phosphorylation; P:signal transduction by protein phosphorylation |
| group_13235 | pilus assembly protein | 1 | C:GO:0016021 | C:integral component of membrane |
| group_13236 | prepilin peptidase | 3 | F:GO:0004190; P:GO:0006508; C:GO:0016021 | F:aspartic-type endopeptidase activity; P:proteolysis; C:integral component of membrane |
| group_13237 | DUF3613 domain-containing protein | | | |
| group_13238 | tetratricopeptide repeat protein | | | |
| group_13239 | type II secretion system protein F | 1 | C:GO:0016021 | C:integral component of membrane |
| group_13240 | type II secretion system protein F | 1 | C:GO:0016021 | C:integral component of membrane |
| group_13241 | secretin | 1 | P:GO:0009306 | P:protein secretion |
| group_13242 | Flp pilus assembly protein CpaB | | | |
| group_13243 | Flp family type IVb pilin | 1 | C:GO:0016021 | C:integral component of membrane |
| group_13244 | glucosyltransferase GtrII-like protein | 1 | C:GO:0016020 | C:membrane |
| yfdG | GtrA family protein | 2 | P:GO:0000271; C:GO:0016021 | P:polysaccharide biosynthetic process; C:integral component of membrane |
| yfdH | glycosyltransferase | 4 | F:GO:0004525; C:GO:0016021; F:GO:0016757; P:GO:0090502 | nponent of membrane; F:transferase activity, transferring glycosyl groups; P:RNA phosphodiester bond hydrolysis, endonucleolytic |
| group_13247 | SDR family NAD(P)-dependent oxidoreductase | 2 | F:GO:0016616; P:GO:0055114 | activity, acting on the CH-OH group of donors, NAD or NADP as acceptor; P:oxidation-reduction process |
| group_13248 | Uncharacterised protein | | | |
| group_13249 | hypothetical protein | | | |
| group_13250 | ABC transporter ATP-binding protein | 2 | F:GO:0005524; F:GO:0016887 | F:ATP binding; F:ATPase activity |
| group_13251 | ABC transporter permease | 3 | C:GO:0005886; C:GO:0016021; F:GO:0016787 | C:plasma membrane; C:integral component of membrane; F:hydrolase activity |
| group_13252 | TolC family protein | 2 | F:GO:0015562; P:GO:0055085 | F:efflux transmembrane transporter activity; P:transmembrane transport |
| group_13253 | HlyD family efflux transporter periplasmic adaptor subunit | | | |
| group_13254 | four helix bundle protein | 1 | C:GO:0005840 | C:ribosome |
| group_13260 | Uncharacterised protein | | | |
| group_13261 | Uncharacterised protein | 3 | C:GO:0005840; P:GO:0006508; F:GO:0008237 | C:ribosome; P:proteolysis; F:metallopeptidase activity |
| group_13262 | S-type pyocin domain-containing protein | 3 | P:GO:0009405; F:GO:0015643; P:GO:0030153 | P:pathogenesis; F:toxic substance binding; P:bacteriocin immunity |
| group_13263 | hypothetical protein | 1 | C:GO:0016021 | C:integral component of membrane |
| group_13264 | abortive phage resistance protein | | | |
| group_13265 | site-specific DNA-methyltransferase | 4 | F:GO:0003677; F:GO:0008170; F:GO:0009007; P:GO:0032775 | ltransferase activity; F:site-specific DNA-methyltransferase (adenine-specific) activity; P:DNA methylation on adenine |
| group_13266 | RNA helicase | 3 | F:GO:0003677; F:GO:0005524; F:GO:0016787 | F:DNA binding; F:ATP binding; F:hydrolase activity |
| group_7235 | transcriptional regulator | | | |
| group_13267 | hypothetical protein | | | |
| group_13268 | cobalamin biosynthesis protein CobW | | | |
| group_13269 | DUF3800 domain-containing protein | | | |
| group_13273 | AP endonuclease | 3 | F:GO:0004519; F:GO:0016853; P:GO:0090305 | :endonuclease activity; F:isomerase activity; P:nucleic acid phosphodiester bond hydrolysis |
| group_13274 | Uncharacterised protein | 1 | C:GO:0016021 | C:integral component of membrane |
| group_13275 | FUSC family protein | 4 | C:GO:0005886; C:GO:0016021; F:GO:0022857; P:GO:0055085 | rane; C:integral component of membrane; F:transmembrane transporter activity; P:transmembrane transport |
| group_7253 | zinc-binding alcohol dehydrogenase family protein | 3 | F:GO:0008270; F:GO:0016491; P:GO:0055114 | F:zinc ion binding; F:oxidoreductase activity; P:oxidation-reduction process |
| group_13276 | effector protein | 1 | C:GO:0016021 | C:integral component of membrane |

| | | | | |
|---|---|---|---|---|
| group_13277 | effector protein | 1 | C:GO:0016021 | C:integral component of membrane |
| group_13278 | hybrid sensor histidine kinase/response regulator | 3 | F:GO:0000155; P:GO:0000160; P:GO:0023014 | or kinase activity; P:phosphorelay signal transduction system; P:signal transduction by protein phosphorylation |
| group_13279 | membrane integrity-associated transporter subunit PqiC | | | |
| group_13280 | Uncharacterised protein | | | |
| group_13281 | Uncharacterised protein | | | |
| group_13282 | VOC family virulence protein | 3 | F:GO:0016829; F:GO:0051213; P:GO:0055114 | F:lyase activity; F:dioxygenase activity; P:oxidation-reduction process |
| group_13283 | NUDIX domain-containing protein | 1 | F:GO:0016787 | F:hydrolase activity |
| group_13284 | TonB-dependent siderophore receptor | 4 | C:GO:0009279; F:GO:0015344; F:GO:0038023; P:GO:0044718 | erophore uptake transmembrane transporter activity; F:signaling receptor activity; P:siderophore transmembrane transport |
| group_13285 | ABC transporter ATP-binding protein | 2 | F:GO:0005524; F:GO:0016887 | F:ATP binding; F:ATPase activity |
| group_13286 | iron ABC transporter substrate-binding protein | | | |
| group_13287 | DUF2474 domain-containing protein | 1 | C:GO:0016021 | C:integral component of membrane |
| cydA_2 | cytochrome ubiquinol oxidase subunit I | 6 | 0005886; F:GO:0009055; C:GO:0016021; P:GO:0019646; F:GO:0046872; C:GO:0( | fer activity; C:integral component of membrane; P:aerobic electron transport chain; F:metal ion binding; C:cytochrome complex |
| group_13288 | DUF2946 domain-containing protein | 1 | C:GO:0016021 | C:integral component of membrane |
| group_1839 | LysR family transcriptional regulator | 3 | F:GO:0003677; F:GO:0003700; P:GO:0006355 | binding; F:DNA-binding transcription factor activity; P:regulation of transcription, DNA-templated |
| group_13289 | copper uptake system-associated protein | | | |
| znuC_3 | manganese ABC transporter ATP-binding protein | 2 | F:GO:0005524; F:GO:0016887 | F:ATP binding; F:ATPase activity |
| hpf_2 | metal ABC transporter substrate-binding protein | 3 | P:GO:0007155; P:GO:0030001; F:GO:0046872 | P:cell adhesion; P:metal ion transport; F:metal ion binding |
| group_13293 | hypothetical protein A9C11_17800 | 2 | C:GO:0016020; C:GO:0016021 | C:membrane; C:integral component of membrane |
| thrS_2 | threonine--tRNA ligase | 7 | ); F:GO:0004829; F:GO:0005524; C:GO:0005737; P:GO:0006435; F:GO:0016829; I | NA ligase activity; F:ATP binding; C:cytoplasm; P:threonyl-tRNA aminoacylation; F:lyase activity; F:metal ion binding |
| group_13295 | 6-carboxy-5,6,7,8-tetrahydropterin synthase | | | |
| cysS_2 | cysteine--tRNA ligase | 7 | 7; F:GO:0005524; C:GO:0005737; P:GO:0006423; F:GO:0008270; F:GO:0016491; | inding; C:cytoplasm; P:cysteinyl-tRNA aminoacylation; F:zinc ion binding; F:oxidoreductase activity; P:oxidation-reduction process |
| group_13297 | GTP-binding protein | | | |
| zur_2 | transcriptional repressor | 3 | F:GO:0003677; F:GO:0003700; P:GO:0006355 | binding; F:DNA-binding transcription factor activity; P:regulation of transcription, DNA-templated |
| group_13299 | N-acetylmuramoyl-L-alanine amidase | 2 | F:GO:0008745; P:GO:0009253 | F:N-acetylmuramoyl-L-alanine amidase activity; P:peptidoglycan catabolic process |
| cdhR_8 | AraC family transcriptional regulator | 3 | F:GO:0003700; P:GO:0006355; F:GO:0043565 | ranscription factor activity; P:regulation of transcription, DNA-templated; F:sequence-specific DNA binding |
| frmA_2 | S-(hydroxymethyl)glutathione dehydrogenase/class III alcohol dehydrogenase | 3 | P:GO:0006069; F:GO:0008270; F:GO:0051903 | thanol oxidation; F:zinc ion binding; F:S-(hydroxymethyl)glutathione dehydrogenase activity |
| group_13302 | peptidase | | | |
| group_13303 | hybrid sensor histidine kinase/response regulator | 3 | P:GO:0007165; F:GO:0016301; P:GO:0016310 | P:signal transduction; F:kinase activity; P:phosphorylation |
| group_13304 | circadian clock protein KaiC | 1 | F:GO:0005524 | F:ATP binding |
| pncC_3 | damage-inducible protein CinA | 1 | F:GO:0016787 | F:hydrolase activity |
| group_7254 | zinc-binding alcohol dehydrogenase family protein | 3 | F:GO:0008270; F:GO:0016491; P:GO:0055114 | F:zinc ion binding; F:oxidoreductase activity; P:oxidation-reduction process |
| group_13305 | alkylhydroperoxidase | 4 | F:GO:0004601; F:GO:0051920; P:GO:0055114; P:GO:0098869 | se activity; F:peroxiredoxin activity; P:oxidation-reduction process; P:cellular oxidant detoxification |
| group_13306 | AraC family transcriptional regulator | 2 | F:GO:0016491; P:GO:0055114 | F:oxidoreductase activity; P:oxidation-reduction process |
| cdhR_9 | GlxA family transcriptional regulator | 3 | F:GO:0003700; P:GO:0006355; F:GO:0043565 | ranscription factor activity; P:regulation of transcription, DNA-templated; F:sequence-specific DNA binding |
| group_13308 | transcriptional regulator | 1 | F:GO:0003677 | F:DNA binding |
| dxs_2 | transketolase | 1 | F:GO:0004802 | F:transketolase activity |
| group_13310 | EAL domain-containing protein | | | |
| nikE_2 | nickel import ATP-binding protein NikE | 6 | 0005524; C:GO:0005886; F:GO:0015413; F:GO:0016151; F:GO:0016887; P:GO:00 | e-coupled nickel transmembrane transporter activity; F:nickel cation binding; F:ATPase activity; P:nickel cation transmembrane transport |
| group_13311 | LysR family transcriptional regulator | 3 | F:GO:0003677; F:GO:0003700; P:GO:0006355 | binding; F:DNA-binding transcription factor activity; P:regulation of transcription, DNA-templated |
| hchA_2 | protein deglycase HchA | 2 | F:GO:0036524; P:GO:0036525 | F:protein deglycase activity; P:protein deglycation |
| group_13313 | ---NA--- | | | |
| group_13314 | Uncharacterised protein | | | |
| group_13315 | hypothetical protein | | | |
| group_13316 | Uncharacterised protein | | | |
| group_13317 | E3 ligase-like protein (putative virulence factor) | 2 | F:GO:0004842; P:GO:0016567 | F:ubiquitin-protein transferase activity; P:protein ubiquitination |
| group_13318 | TonB-dependent siderophore receptor | 5 | C:GO:0009279; F:GO:0015344; C:GO:0016021; F:GO:0038023; P:GO:0044718 | nsmembrane transporter activity; C:integral component of membrane; F:signaling receptor activity; P:siderophore transmembrane transport |
| group_13319 | chemotaxis protein | 4 | F:GO:0004888; P:GO:0006935; P:GO:0007165; C:GO:0016020 | transmembrane signaling receptor activity; P:chemotaxis; P:signal transduction; C:membrane |
| group_13320 | sensor domain-containing phosphodiesterase | | | |
| group_13321 | Uncharacterised protein | | | |
| group_13323 | membrane protein | 1 | C:GO:0016021 | C:integral component of membrane |
| group_13324 | Uncharacterised protein | 1 | C:GO:0016021 | C:integral component of membrane |
| group_13325 | putative membrane protein | 1 | C:GO:0016021 | C:integral component of membrane |
| group_13326 | type IV secretion protein Rhs | | | |
| group_13327 | hypothetical protein | | | |
| group_13328 | DUF1795 domain-containing protein | | | |
| rhsA | type IV secretion protein Rhs | 3 | F:GO:0004519; C:GO:0016021; P:GO:0090305 | iclease activity; C:integral component of membrane; P:nucleic acid phosphodiester bond hydrolysis |
| group_13329 | cation:proton antiporter | | | |
| group_269 | type IV secretion protein Rhs | 3 | F:GO:0004519; C:GO:0016021; P:GO:0090305 | iclease activity; C:integral component of membrane; P:nucleic acid phosphodiester bond hydrolysis |
| group_270 | type IV secretion protein Rhs | 3 | F:GO:0004519; C:GO:0016021; P:GO:0090305 | iclease activity; C:integral component of membrane; P:nucleic acid phosphodiester bond hydrolysis |
| group_13330 | cytoplasmic protein | | | |
| group_268 | type IV secretion protein Rhs | 1 | C:GO:0016021 | C:integral component of membrane |
| group_13331 | hypothetical protein | 2 | C:GO:0016020; C:GO:0016021 | C:membrane; C:integral component of membrane |
| group_13332 | hypothetical protein | | | |
| rhsB | sugar-binding protein | 1 | C:GO:0016021 | C:integral component of membrane |
| group_13333 | MULTISPECIES: hypothetical protein | | | |
| group_13334 | putative transmembrane protein | 1 | C:GO:0016021 | C:integral component of membrane |
| group_50 | DUF3077 domain-containing protein | | | |
| group_107 | Uncharacterised protein | | | |
| group_13335 | IclR family transcriptional regulator | 2 | F:GO:0003677; P:GO:0006355 | F:DNA binding; P:regulation of transcription, DNA-templated |
| group_13336 | NUDIX domain-containing protein | 1 | F:GO:0016787 | F:hydrolase activity |
| group_13337 | response regulator | 3 | F:GO:0000155; P:GO:0000160; P:GO:0023014 | or kinase activity; P:phosphorelay signal transduction system; P:signal transduction by protein phosphorylation |
| group_13338 | NUDIX domain-containing protein | 1 | F:GO:0016787 | F:hydrolase activity |
| group_1867 | IS66 family transposase | | | |
| group_13339 | DNA mismatch repair protein MutT | 1 | F:GO:0016787 | F:hydrolase activity |
| group_13340 | universal stress protein UspA | | | |
| group_13341 | SH3 domain-containing protein | | | |
| group_13342 | Uncharacterised protein | | | |
| group_3039 | transcriptional regulator | 2 | F:GO:0003677; P:GO:0006355 | F:DNA binding; P:regulation of transcription, DNA-templated |
| group_13343 | hypothetical protein | | | |
| group_13344 | hypothetical protein | | | |
| group_13345 | DUF4062 domain-containing protein | | | |
| group_13346 | AAA family ATPase | | | |
| group_13347 | hypothetical protein DCO48_21420 | | | |
| group_13348 | hypothetical protein DCO48_21425 | | | |
| group_13349 | ---NA--- | | | |
| group_13350 | AAA family ATPase | | | |
| group_13351 | hypothetical protein | | | |
| group_13352 | hypothetical protein | | | |
| group_13353 | integrase | 3 | F:GO:0003677; P:GO:0006310; P:GO:0015074 | F:DNA binding; P:DNA recombination; P:DNA integration |
| group_13354 | integrase | 3 | F:GO:0003677; P:GO:0006310; P:GO:0015074 | F:DNA binding; P:DNA recombination; P:DNA integration |
| group_13355 | integrase | 3 | F:GO:0003677; P:GO:0006310; P:GO:0015074 | F:DNA binding; P:DNA recombination; P:DNA integration |
| group_13356 | type IV pili biogenesis protein FimT | 2 | C:GO:0015627; P:GO:0015628 | :type II protein secretion system complex; P:protein secretion by the type II secretion system |
| group_13357 | pilus assembly protein PilW | 1 | C:GO:0016021 | C:integral component of membrane |
| group_13358 | type IV pilus assembly protein PilX | | | |
| group_3064 | integrase | 3 | F:GO:0003677; P:GO:0006310; P:GO:0015074 | F:DNA binding; P:DNA recombination; P:DNA integration |
| group_13359 | hypothetical protein | | | |
| group_13360 | ATP-binding protein | 3 | F:GO:0005524; F:GO:0016301; P:GO:0016310 | F:ATP binding; F:kinase activity; P:phosphorylation |
| group_13364 | heat-shock protein HtpX | | | |
| group_13365 | MULTISPECIES: hypothetical protein | | | |
| group_102 | Copper resistance protein B precursor | | | |
| group_13366 | hypothetical protein IEC33019_0010 | | | |
| group_5184 | TniQ protein | | | |
| group_13367 | hypothetical protein | | | |
| group_136 | serine hydroxymethyltransferase | 7 | 2; C:GO:0005737; F:GO:0008168; P:GO:0019264; F:GO:0030170; P:GO:0032259; | lmethyltransferase activity; P:glycine biosynthetic process from serine; F:pyridoxal phosphate binding; P:methylation; P:tetrahydrofolate interconversion |
| group_13368 | MFS transporter | 4 | C:GO:0005886; C:GO:0016021; F:GO:0042910; P:GO:1990961 | mbrane; F:xenobiotic transmembrane transporter activity; P:xenobiotic detoxification by transmembrane export across the plasma membrane |
| group_13369 | thermostable hemolysin | | | |
| group_13370 | AMP-dependent synthetase | 1 | F:GO:0016874 | F:ligase activity |
| group_13371 | taurine catabolism dioxygenase TauD | 2 | F:GO:0051213; P:GO:0055114 | F:dioxygenase activity; P:oxidation-reduction process |
| group_13372 | LysR family transcriptional regulator | 3 | F:GO:0003677; F:GO:0003700; P:GO:0006355 | binding; F:DNA-binding transcription factor activity; P:regulation of transcription, DNA-templated |
| group_13373 | MarR family transcriptional regulator | 3 | F:GO:0003677; F:GO:0003700; P:GO:0006355 | binding; F:DNA-binding transcription factor activity; P:regulation of transcription, DNA-templated |
| bpoC | alpha/beta hydrolase | 4 | F:GO:0004601; F:GO:0016787; P:GO:0055114; P:GO:0098869 | idase activity; F:hydrolase activity; P:oxidation-reduction process; P:cellular oxidant detoxification |
| group_13375 | MFS transporter | 2 | C:GO:0016021; P:GO:0055085 | C:integral component of membrane; P:transmembrane transport |
| group_13376 | hypothetical protein | | | |
| group_13377 | hypothetical protein BN2475_140048 | | | |
| group_13378 | hypothetical protein BN2475_140049 | | | |
| group_13379 | Lyzozyme M1 (1,4-beta-N-acetylmuramidase), GH25 family | 2 | P:GO:0009057; F:GO:0016787 | P:macromolecule catabolic process; F:hydrolase activity |
| group_13380 | curculin domain-containing protein | 4 | F:GO:0004252; P:GO:0006508; F:GO:0008237; F:GO:0030246 | e-type endopeptidase activity; P:proteolysis; F:metallopeptidase activity; F:carbohydrate binding |
| group_13381 | Hypothetical protein (plasmid) | | | |
| group_7327 | Uncharacterised protein | | | |
| group_13398 | Uncharacterised protein | | | |
| group_7328 | Uncharacterised protein | | | |
| recD2 | Nuclease-related domain-containing protein | 6 | 0004386; F:GO:0004672; F:GO:0005524; P:GO:0006468; C:GO:0016020; C:GO:00 | tein kinase activity; F:ATP binding; P:protein phosphorylation; C:membrane; C:integral component of membrane |
| group_13400 | hypothetical protein | | | |
| group_13401 | integrase | 3 | F:GO:0003677; P:GO:0006310; P:GO:0015074 | F:DNA binding; P:DNA recombination; P:DNA integration |
| group_13402 | MULTISPECIES: hypothetical protein | | | |
| group_13403 | Uncharacterised protein | | | |
| group_13404 | hypothetical protein | | | |
| group_3076 | Uncharacterised protein | | | |
| group_3075 | Uncharacterised protein | | | |
| group_5203 | transposase | | | |
| group_13405 | transposase | 3 | F:GO:0004803; P:GO:0006313; F:GO:0043565 | F:transposase activity; P:transposition, DNA-mediated; F:sequence-specific DNA binding |
| group_3037 | transposase | | | |
| group_1874 | IS66 family transposase | | | |
| group_3085 | group II intron reverse transcriptase/maturase | 3 | F:GO:0003723; F:GO:0003964; P:GO:0006278 | binding; F:RNA-directed DNA polymerase activity; P:RNA-dependent DNA biosynthetic process |
| group_13406 | IS256 family transposase | 3 | F:GO:0003677; F:GO:0004803; P:GO:0006313 | F:DNA binding; F:transposase activity; P:transposition, DNA-mediated |
| group_3208 | ---NA--- | | | |
| group_3209 | ---NA--- | | | |
| group_13407 | Uncharacterised protein | | | |
| group_13409 | Rieske (2Fe-2S) protein | 4 | F:GO:0004497; F:GO:0051213; F:GO:0051537; P:GO:0055114 | enase activity; F:dioxygenase activity; F:2 iron, 2 sulfur cluster binding; P:oxidation-reduction process |
| group_13410 | oxidoreductase | 7 | 3; F:GO:0009055; P:GO:0022900; P:GO:0032259; F:GO:0046872; F:GO:0051213; | sfer activity; P:electron transport chain; P:methylation; F:metal ion binding; F:dioxygenase activity; F:2 iron, 2 sulfur cluster binding |
| group_13411 | helix-turn-helix transcriptional regulator | 2 | F:GO:0003677; P:GO:0006355 | F:DNA binding; P:regulation of transcription, DNA-templated |
| group_4100 | usher CupB3 | 4 | P:GO:0009297; F:GO:0015473; C:GO:0016020; P:GO:0055085 | P:pilus assembly; F:fimbrial usher porin activity; C:membrane; P:transmembrane transport |
| group_13412 | hypothetical protein | | | |
| group_13413 | transcriptional regulator | | | |
| group_13414 | AlpA family transcriptional regulator | | | |
| group_13415 | inovirus Gp2 family protein | | | |
| group_13416 | NERD domain-containing protein | | | |
| group_13417 | plasmid pRiA4b ORF-3 family protein | | | |
| group_1175 | WYL domain-containing protein | | | |
| group_13418 | tRNA anti-like protein | 1 | C:GO:0016021 | C:integral component of membrane |
| group_13419 | transcriptional regulator | 1 | F:GO:0004386 | F:helicase activity |
| group_13420 | chromosome partitioning protein | 1 | F:GO:0016787 | F:hydrolase activity |
| group_13421 | DUF1156 domain-containing protein | | | |
| group_13422 | hypothetical protein | | | |
| group_13423 | DUF499 domain-containing protein | 1 | F:GO:0005524 | F:ATP binding |
| group_13424 | Uncharacterised protein | | | |
| group_13425 | Uncharacterised protein | | | |
| group_13426 | transcriptional regulator | 1 | F:GO:0003677 | F:DNA binding |
| group_13427 | hypothetical protein | | | |
| group_13428 | Zn-dependent peptidase ImmA, M78 family | 1 | F:GO:0003677 | F:DNA binding |
| group_13429 | DNA-binding protein | 1 | F:GO:0003677 | F:DNA binding |
| group_13430 | integrase | 3 | F:GO:0003677; P:GO:0006310; P:GO:0015074 | F:DNA binding; P:DNA recombination; P:DNA integration |
| group_13431 | hypothetical protein ALP65_03490 | | | |
| group_13433 | DUF2007 domain-containing protein | | | |
| group_13435 | integrase | 3 | F:GO:0003677; P:GO:0006310; P:GO:0015074 | F:DNA binding; P:DNA recombination; P:DNA integration |
| repA | replication protein A | | | |
| group_13437 | ABC transporter ATPase | | | |
| group_13438 | Uncharacterised protein | 1 | C:GO:0016021 | C:integral component of membrane |
| group_13439 | P-type conjugative transfer protein TrbL | 2 | C:GO:0016021; P:GO:0030255 | C:integral component of membrane; P:protein secretion by the type IV secretion system |
| group_13440 | P-type conjugative transfer protein TrbJ | | | |
| group_13441 | Uncharacterised protein | | | |
| group_13442 | conjugal transfer protein TraI | | | |
| group_13443 | Uncharacterised protein | | | |
| group_13444 | Uncharacterised protein | | | |
| group_13445 | Competence protein | | | |
| group_13446 | AAA family ATPase | | | |
| group_13447 | integrase | 3 | F:GO:0003677; P:GO:0006310; P:GO:0015074 | F:DNA binding; P:DNA recombination; P:DNA integration |
| group_13448 | MULTISPECIES: hypothetical protein | | | |
| group_13449 | DNA-binding protein | 1 | F:GO:0003677 | F:DNA binding |
| group_13450 | AlpA family transcriptional regulator | | | |
| group_13451 | subtilase family protein | 2 | F:GO:0004252; P:GO:0006508 | F:serine-type endopeptidase activity; P:proteolysis |
| group_13452 | ATPase | 3 | F:GO:0005524; P:GO:0006508; F:GO:0008237 | F:ATP binding; P:proteolysis; F:metallopeptidase activity |
| group_13453 | antirestriction protein ArdA | | | |
| group_13454 | class I SAM-dependent methyltransferase | 2 | F:GO:0008168; P:GO:0032259 | F:methyltransferase activity; P:methylation |
| group_13455 | Uncharacterised protein | | | |
| group_13456 | hypothetical protein | | | |
| group_13457 | ---NA--- | | | |
| group_13458 | Uncharacterised protein | | | |
| group_13459 | HNH endonuclease | 2 | F:GO:0004519; P:GO:0090305 | F:endonuclease activity; P:nucleic acid phosphodiester bond hydrolysis |
| group_13460 | restriction endonuclease subunit S | 2 | F:GO:0004519; P:GO:0090305 | F:endonuclease activity; P:nucleic acid phosphodiester bond hydrolysis |
| group_13461 | nuclease | 3 | F:GO:0003676; F:GO:0004519; P:GO:0090305 | nucleic acid binding; F:endonuclease activity; P:nucleic acid phosphodiester bond hydrolysis |
| group_13462 | chemotaxis protein | 1 | C:GO:0016021 | C:integral component of membrane |
| group_13463 | chemotaxis protein MotB | 2 | C:GO:0016021; P:GO:0016539 | C:integral component of membrane; P:intein-mediated protein splicing |
| group_13464 | Uncharacterised protein | | | |
| group_13465 | N-formylmethionyl-tRNA deformylase | 2 | F:GO:0004386; F:GO:0005524 | F:helicase activity; F:ATP binding |
| group_13466 | Iron-binding zinc finger CDGSH type | 2 | C:GO:0043231; F:GO:0051537 | C:intracellular membrane-bounded organelle; F:2 iron, 2 sulfur cluster binding |
| group_13468 | hypothetical protein Q003_01486 | | | |

| | | | | |
|---|---|---|---|---|
| imm | bacteriocin immunity protein | 2 | F:GO:0015643; P:GO:0030153 | F:toxic substance binding; P:bacteriocin immunity |
| group_13470 | S-type Pyocin | 6 | :0004519; F:GO:0005102; P:GO:0009405; P:GO:0009617; P:GO:0019835; P:GO:00 | ling receptor binding; P:pathogenesis; P:response to bacterium; P:cytolysis; P:nucleic acid phosphodiester bond hydrolysis |
| group_2772 | DUF2235 domain-containing protein | | | |
| group_13475 | Uncharacterised protein | | | |
| group_13476 | MULTISPECIES: hypothetical protein | | | |
| group_7339 | MULTISPECIES: hypothetical protein | | | |
| group_30 | transposase | 3 | F:GO:0003677; F:GO:0004803; P:GO:0006313 | F:DNA binding; F:transposase activity; P:transposition, DNA-mediated |
| group_13492 | O-antigen chain length regulator | 2 | P:GO:0009103; C:GO:0016021 | P:lipopolysaccharide biosynthetic process; C:integral component of membrane |
| group_13493 | flippase | 1 | C:GO:0016021 | C:integral component of membrane |
| group_13494 | hypothetical protein | 1 | C:GO:0016021 | C:integral component of membrane |
| group_13495 | glycosyltransferase family 2 protein | | | |
| group_13496 | glycosyltransferase family 4 protein | | | |
| capD | UDP-glucose 4-epimerase | 2 | F:GO:0003978; P:GO:0009103 | F:UDP-glucose 4-epimerase activity; P:lipopolysaccharide biosynthetic process |
| wbjC | capsular biosynthesis protein | 2 | F:GO:0016853; F:GO:0050662 | F:isomerase activity; F:coenzyme binding |
| group_13499 | hypothetical protein PA13_1001060 | | | |
| group_13500 | UDP-N-acetylglucosamine 2-epimerase (non-hydrolyzing) | 1 | F:GO:0008761 | F:UDP-N-acetylglucosamine 2-epimerase activity |
| group_13501 | putative glycosyl transferase | 2 | C:GO:0016021; F:GO:0016740 | C:integral component of membrane; F:transferase activity |
| group_13502 | NAD-dependent dehydratase | 2 | F:GO:0003978; F:GO:0050662 | F:UDP-glucose 4-epimerase activity; F:coenzyme binding |
| group_13503 | glycosyl transferase | 3 | F:GO:0008963; C:GO:0016021; F:GO:0036380 | ctivity; C:integral component of membrane; F:UDP-N-acetylglucosamine-undecaprenyl-phosphate N-acetylglucosaminephosphotransferase activity |
| group_1850 | rhomboid family intramembrane serine protease | 3 | F:GO:0004252; P:GO:0006508; C:GO:0016021 | F:serine-type endopeptidase activity; P:proteolysis; C:integral component of membrane |
| group_13507 | Uncharacterised protein | | | |
| group_7345 | R body protein RebB-like protein | | | |
| group_7346 | R body protein RebB-like protein | | | |
| group_13509 | Mu-like prophage I protein | | | |
| group_13510 | Uncharacterised protein | | | |
| group_13511 | Uncharacterised protein | | | |
| group_13512 | hypothetical protein | | | |
| group_13513 | Uncharacterised protein | | | |
| group_13514 | RebB like protein | | | |
| group_13515 | RNA polymerase subunit sigma-70 | 4 | F:GO:0003677; F:GO:0003700; F:GO:0016987; P:GO:2000142 | binding transcription factor activity; F:sigma factor activity; P:regulation of DNA-templated transcription, initiation |
| group_13516 | transcriptional regulator | 2 | F:GO:0003677; P:GO:0006355 | F:DNA binding; P:regulation of transcription, DNA-templated |
| group_13517 | integrase | 3 | F:GO:0003677; P:GO:0006310; P:GO:0015074 | F:DNA binding; P:DNA recombination; P:DNA integration |
| group_13518 | Pyocin activator protein PrtN | 1 | P:GO:0006355 | P:regulation of transcription, DNA-templated |
| group_13519 | Uncharacterised protein | | | |
| group_13520 | hypothetical protein | | | |
| group_13521 | glycoside hydrolase | 1 | F:GO:0016787 | F:hydrolase activity |
| group_13522 | membrane protein | | | |
| group_13523 | TetR family transcriptional regulator | | | |
| group_13524 | two-component sensor | 1 | C:GO:0016021 | C:integral component of membrane |
| group_13525 | transcriptional regulator | 1 | P:GO:0006355 | P:regulation of transcription, DNA-templated |
| group_13526 | hypothetical protein | | | |
| group_13527 | DNA-binding protein | 1 | F:GO:0003677 | F:DNA binding |
| group_13528 | putative transcriptional regulator | | | |
| group_13529 | taurine dioxygenase | | | |
| group_13530 | peptidase S24 | 1 | F:GO:0003677 | F:DNA binding |
| group_13531 | putative zinc-binding dehydrogenase | | | |
| ybiI_1 | conjugal transfer protein TraR | 1 | F:GO:0008270 | F:zinc ion binding |
| group_13533 | conjugal transfer protein TraC | | | |
| group_13534 | MULTISPECIES: hypothetical protein | | | |
| group_13535 | pyocin R2, holin | 1 | C:GO:0016021 | C:integral component of membrane |
| group_13536 | DNA packaging protein | | | |
| group_13537 | terminase | | | |
| group_13538 | preprotein translocase subunit SecA | | | |
| group_13539 | phage portal protein | 2 | F:GO:0005198; P:GO:0019068 | F:structural molecule activity; P:virion assembly |
| group_13540 | hypothetical phage-related protein | 2 | P:GO:0006508; F:GO:0008233 | P:proteolysis; F:peptidase activity |
| group_13541 | head decoration protein | | | |
| group_13542 | capsid protein | | | |
| group_13543 | Putative DnaB-like replicative helicase | | | |
| group_13544 | Putative lipoprotein | | | |
| group_13545 | hypothetical protein | | | |
| group_13546 | hypothetical protein | | | |
| group_13547 | phage baseplate protein | | | |
| group_13548 | UDP-N-acetyl-D-mannosaminuronate dehydrogenase | | | |
| group_2192 | phage baseplate protein | | | |
| group_13549 | baseplate assembly protein | | | |
| group_13550 | phage tail protein I | | | |
| group_13551 | phage tail protein | | | |
| group_13552 | phage tail protein | | | |
| gpFI_1 | phage tail protein | | | |
| group_6282 | phage major tail tube protein | | | |
| group_13553 | phage tail assembly protein | | | |
| group_13554 | phage tail protein | | | |
| group_13555 | phage tail protein | | | |
| group_13556 | phage tail protein | | | |
| group_13557 | DNA primase | | | |
| group_2711 | glycoside hydrolase family 19 | 3 | F:GO:0004568; P:GO:0006032; P:GO:0016998 | F:chitinase activity; P:chitin catabolic process; P:cell wall macromolecule catabolic process |
| group_13558 | unknown | | | |
| group_13559 | hypothetical protein | | | |
| group_5255 | hypothetical protein | | | |
| group_13560 | DUF159 family protein | 2 | P:GO:0006508; F:GO:0008233 | P:proteolysis; F:peptidase activity |
| group_7348 | hypothetical protein | | | |
| aplIM | DNA (cytosine-5-)-methyltransferase | 2 | F:GO:0003886; P:GO:0090116 | F:DNA (cytosine-5-)-methyltransferase activity; P:C-5 methylation of cytosine |
| group_13562 | Eco29kI family restriction endonuclease | 2 | F:GO:0004519; P:GO:0090305 | F:endonuclease activity; P:nucleic acid phosphodiester bond hydrolysis |
| group_13563 | GIY-YIG nuclease family protein | 2 | F:GO:0004519; P:GO:0090305 | F:endonuclease activity; P:nucleic acid phosphodiester bond hydrolysis |
| group_13564 | HNH endonuclease | 3 | F:GO:0003676; F:GO:0004519; P:GO:0090305 | nucleic acid binding; F:endonuclease activity; P:nucleic acid phosphodiester bond hydrolysis |
| vsr | very short patch repair endonuclease | 3 | F:GO:0004519; P:GO:0006298; P:GO:0090305 | F:endonuclease activity; P:mismatch repair; P:nucleic acid phosphodiester bond hydrolysis |
| group_13566 | HNH endonuclease | 2 | F:GO:0004519; P:GO:0090305 | F:endonuclease activity; P:nucleic acid phosphodiester bond hydrolysis |
| group_13567 | WYL domain-containing protein | | | |
| group_13568 | nucleotide pyrophosphohydrolase | 3 | P:GO:0009143; F:GO:0046872; F:GO:0047429 | e triphosphate catabolic process; F:metal ion binding; F:nucleoside-triphosphate diphosphatase activity |
| group_4146 | RNA-splicing ligase RtcB | 3 | P:GO:0006396; F:GO:0008452; F:GO:0046872 | P:RNA processing; F:RNA ligase activity; F:metal ion binding |
| group_13569 | integrase | 3 | F:GO:0003677; P:GO:0006310; P:GO:0015074 | F:DNA binding; P:DNA recombination; P:DNA integration |
| group_13570 | DUF4224 domain-containing protein | | | |
| group_7351 | Uncharacterised protein | | | |
| group_13571 | hypothetical protein | | | |
| group_13572 | hypothetical protein | | | |
| group_13573 | hypothetical protein | | | |
| group_13574 | DUF1643 domain-containing protein | | | |
| group_7352 | Uncharacterised protein | | | |
| group_7353 | DUF4406 domain-containing protein | | | |
| group_13575 | cytosine methyltransferase | 2 | F:GO:0008168; P:GO:0032259 | F:methyltransferase activity; P:methylation |
| group_7354 | DNA cytosine methyltransferase | 2 | F:GO:0008168; P:GO:0032259 | F:methyltransferase activity; P:methylation |
| recF_2 | AAA family ATPase | | | |
| group_7356 | cell division protein FtsK | 1 | P:GO:0051301 | P:cell division |
| group_13576 | MULTISPECIES: hypothetical protein | | | |
| group_13577 | DNA recombination protein RecT | | | |
| group_13578 | PD-(D/E)XK nuclease family protein | | | |
| group_13579 | Uncharacterised protein | | | |
| group_13580 | hypothetical protein | | | |
| group_7357 | Uncharacterised protein | | | |
| group_13581 | MULTISPECIES: hypothetical protein | | | |
| group_7358 | Uncharacterised protein | | | |
| group_13582 | phage protein | | | |
| csrA_1 | carbon storage regulator | 6 | :0005737; P:GO:0006109; P:GO:0006402; P:GO:0045947; P:GO:0045948; F:GO:00 | P:mRNA catabolic process; P:negative regulation of translational initiation; P:positive regulation of translational initiation; F:mRNA 5'-UTR binding |
| group_13584 | hypothetical protein | | | |
| group_7359 | hypothetical protein | | | |
| group_7361 | hypothetical protein | | | |
| group_13585 | DUF1654 domain-containing protein | | | |
| group_13586 | heavy metal transporter | | | |
| lexA_3 | peptidase S24 | 1 | F:GO:0003677 | F:DNA binding |
| group_7363 | transcriptional regulator | 1 | F:GO:0003677 | F:DNA binding |
| group_13588 | hypothetical protein | | | |
| group_7364 | hypothetical protein | | | |
| group_13589 | Replication protein P | 4 | F:GO:0000287; P:GO:0006270; P:GO:0006281; P:GO:0006310 | magnesium ion binding; P:DNA replication initiation; P:DNA repair; P:DNA recombination |
| group_7365 | RusA family crossover junction endodeoxyribonuclease | 3 | F:GO:0000287; P:GO:0006281; P:GO:0006310 | F:magnesium ion binding; P:DNA repair; P:DNA recombination |
| group_7366 | hypothetical protein | | | |
| group_13590 | MULTISPECIES: hypothetical protein | | | |
| group_13591 | phage-related protein | 1 | C:GO:0016021 | C:integral component of membrane |
| group_13592 | phage holin family protein | 1 | C:GO:0016021 | C:integral component of membrane |
| group_13593 | terminase small subunit | | | |
| group_13594 | terminase | | | |
| group_13595 | DNA-binding protein | 1 | F:GO:0003677 | F:DNA binding |
| group_13596 | Phage head morphogenesis protein, SPP1 gp7 | | | |
| group_13597 | Uncharacterised protein | | | |
| group_13598 | hypothetical bacteriophage protein | | | |
| group_13599 | hypothetical protein | | | |
| group_13600 | hypothetical protein | | | |
| group_13601 | hypothetical protein | | | |
| group_13602 | Uncharacterised protein | | | |
| group_13603 | prophage protein | | | |
| group_13604 | hypothetical phage protein | | | |
| group_13605 | phage tail protein | | | |
| group_13606 | hypothetical protein | | | |
| group_13607 | phage tail tape measure protein | | | |
| group_13608 | phage tail protein | | | |
| group_13609 | phage minor tail protein L | | | |
| group_13610 | hydrolase Nlp/P60 | | | |
| group_13611 | hypothetical protein | | | |
| group_13612 | MULTISPECIES: hypothetical protein | | | |
| group_13613 | tail assembly protein | | | |
| group_13614 | phage tail protein | | | |
| group_7367 | Uncharacterised protein | | | |
| group_5258 | tail protein | | | |
| group_2712 | glycoside hydrolase family 19 | 3 | F:GO:0004568; P:GO:0006032; P:GO:0016998 | F:chitinase activity; P:chitin catabolic process; P:cell wall macromolecule catabolic process |
| group_7369 | hypothetical protein | | | |
| group_7371 | putative lipoprotein | | | |
| group_5257 | Phage protein | | | |
| group_7349 | hypothetical protein Q094_01836 | | | |
| group_13617 | Uncharacterised protein | | | |
| group_13620 | Uncharacterised protein | | | |
| group_7373 | recombinase XerC | 3 | F:GO:0003677; P:GO:0006310; P:GO:0015074 | F:DNA binding; P:DNA recombination; P:DNA integration |
| group_7374 | relaxase | | | |
| group_13621 | SIR2 family protein | | | |
| group_13622 | ATPase | 1 | F:GO:0005524 | F:ATP binding |
| group_13623 | ATP-binding protein | 1 | F:GO:0005524 | F:ATP binding |
| group_7375 | plasmid stabilization protein ParE | | | |
| group_7376 | addiction module antidote protein | 1 | P:GO:0006355 | P:regulation of transcription, DNA-templated |
| group_13624 | DUF3742 domain-containing protein | 1 | C:GO:0016021 | C:integral component of membrane |
| group_7377 | conjugal transfer protein TraG | 1 | C:GO:0016021 | C:integral component of membrane |
| group_7378 | putative membrane protein | 1 | C:GO:0016021 | C:integral component of membrane |
| group_13625 | integrating conjugative element protein | | | |
| group_7379 | integrating conjugative element protein | | | |
| group_13491 | MULTISPECIES: hypothetical protein | | | |
| group_7343 | Uncharacterised protein | | | |
| group_3087 | protein-disulfide isomerase | 3 | C:GO:0005623; F:GO:0016853; P:GO:0045454 | C:cell; F:isomerase activity; P:cell redox homeostasis |
| group_13626 | Uncharacterised protein | | | |
| group_15112 | conjugative transfer ATPase | 2 | F:GO:0003677; F:GO:0005524 | F:DNA binding; F:ATP binding |
| group_7380 | conjugal transfer protein | | | |
| group_13627 | integrating conjugative element protein | 1 | C:GO:0016021 | C:integral component of membrane |
| group_7381 | integrating conjugative element protein | | | |
| group_7382 | integrating conjugative element protein | 1 | C:GO:0016021 | C:integral component of membrane |
| group_13628 | conjugal transfer protein | 1 | C:GO:0016021 | C:integral component of membrane |
| group_7383 | integrating conjugative element membrane protein | 1 | C:GO:0016021 | C:integral component of membrane |
| group_7384 | integrating conjugative element protein | 1 | C:GO:0016021 | C:integral component of membrane |
| group_7385 | type III effector Hop protein | | | |
| group_7386 | Uncharacterised protein | | | |
| group_13629 | Uncharacterised protein | | | |
| addA | DNA helicase | 4 | F:GO:0003677; F:GO:0003678; F:GO:0005524; P:GO:0032508 | F:DNA binding; F:DNA helicase activity; F:ATP binding; P:DNA duplex unwinding |
| group_7388 | integrating conjugative element membrane protein | 1 | C:GO:0016021 | C:integral component of membrane |
| group_4041 | conjugative coupling factor TraD, PFGI-1 class | 1 | C:GO:0016021 | C:integral component of membrane |
| group_7389 | dTDP-glucose 4,6-dehydratase | | | |
| group_13630 | integrating conjugative element protein | | | |
| group_7390 | lytic transglycosylase | | | |
| group_7391 | integrating conjugative element protein | | | |
| group_15110 | methyl-accepting chemotaxis protein | | | |
| group_13631 | transposase | 2 | F:GO:0003676; P:GO:0015074 | F:nucleic acid binding; P:DNA integration |
| group_7392 | transposase | 3 | F:GO:0003677; F:GO:0004803; P:GO:0006313 | F:DNA binding; F:transposase activity; P:transposition, DNA-mediated |
| group_13632 | class I SAM-dependent methyltransferase | 3 | F:GO:0003676; F:GO:0008168; P:GO:0032259 | F:nucleic acid binding; F:methyltransferase activity; P:methylation |

| | | | | |
|---|---|---|---|---|
| group_7393 | Uncharacterised protein | | | |
| group_13633 | Uncharacterised protein | | | |
| group_13634 | Uncharacterised protein | | | |
| group_7394 | DUF3275 family protein | | | |
| group_13635 | Uncharacterised protein | | | |
| group_13636 | hypothetical protein | | | |
| group_13637 | Uncharacterised protein | | | |
| group_7395 | Uncharacterised protein | | | |
| group_7396 | Uncharacterised protein | | | |
| group_7397 | DUF3577 domain-containing protein | | | |
| group_13638 | putative membrane protein | 1 | C:GO:0016021 | C:integral component of membrane |
| group_13639 | Uncharacterised protein | 1 | C:GO:0016021 | C:integral component of membrane |
| group_7398 | CP43 (plasmid) | | | |
| group_7399 | pilus assembly protein | | | |
| group_13640 | pilus assembly protein PilV | 1 | C:GO:0016021 | C:integral component of membrane |
| pilT_3 | twitching motility protein PilT | | | |
| group_13642 | pilus assembly protein PilX | 1 | C:GO:0016021 | C:integral component of membrane |
| tcpE | type II secretion system protein F | 1 | C:GO:0016021 | C:integral component of membrane |
| group_13644 | type IV B pilus protein | | | |
| group_13645 | pilus assembly protein PilX | | | |
| group_7400 | pilus assembly protein | | | |
| group_13646 | PilN family type IVB pilus formation outer membrane protein | 3 | P:GO:0009297; P:GO:0009306; C:GO:0019867 | P:pilus assembly; P:protein secretion; C:outer membrane |
| group_7401 | pilus assembly protein PilL | | | |
| group_13647 | three-Cys-motif partner protein TcmP | | | |
| group_13648 | ABC transporter ATP-binding protein | 1 | F:GO:0005524 | F:ATP binding |
| rapA_3 | DEAD/DEAH box helicase | 2 | F:GO:0004386; F:GO:0005524 | F:helicase activity; F:ATP binding |
| group_7403 | MULTISPECIES: hypothetical protein | | | |
| topA_2 | DNA topoisomerase I | 4 | F:GO:0003677; F:GO:0003917; P:GO:0006265; F:GO:0046872 | topoisomerase type I (single strand cut, ATP-independent) activity; P:DNA topological change; F:metal ion binding |
| group_13649 | hypothetical protein | | | |
| group_13650 | cobalt-precorrin-6A synthase | | | |
| group_13651 | single-stranded DNA-binding protein | 2 | F:GO:0003697; P:GO:0006260 | F:single-stranded DNA binding; P:DNA replication |
| group_13652 | phage regulatory protein | | | |
| group_7405 | integrase | | | |
| group_7406 | integrating conjugative element protein | | | |
| group_13653 | WD40 repeat-containing protein | | | |
| group_7407 | hypothetical protein | | | |
| group_7408 | DUF2857 domain-containing protein | | | |
| group_7409 | transcriptional regulator | | | |
| group_7410 | DNA-binding protein | 2 | F:GO:0003677; P:GO:0006355 | F:DNA binding; P:regulation of transcription, DNA-templated |
| yejK_2 | nucleoid-associated protein YejK | 2 | C:GO:0005737; C:GO:0009295 | C:cytoplasm; C:nucleoid |
| group_13654 | cobyrinic acid a,c-diamide synthase | | | |
| group_7411 | hyphothetical protein CP13 | | | |
| group_7412 | CP12 | | | |
| group_7413 | CP11 | | | |
| group_7414 | hypothetical protein | | | |
| group_13655 | Uncharacterised protein | | | |
| dnaB_2 | replicative DNA helicase | 6 | 0003677; F:GO:0003678; F:GO:0005524; P:GO:0006269; P:GO:0032508; C:GO:19 | activity; F:ATP binding; P:DNA replication, synthesis of RNA primer; P:DNA duplex unwinding; C:primosome complex |
| group_7415 | conserved domain protein | | | |
| group_7416 | HNH endonuclease | 2 | F:GO:0004519; P:GO:0090305 | F:endonuclease activity; P:nucleic acid phosphodiester bond hydrolysis |
| group_15098 | DUF2786 domain-containing protein | | | |
| group_13656 | MULTISPECIES: hypothetical protein | | | |
| group_13657 | MULTISPECIES: hypothetical protein | | | |
| group_13658 | Uncharacterised protein | | | |
| group_13659 | Orf50 protein | | | |
| soj_3 | cobyrinic acid a,c-diamide synthase | | | |
| group_13662 | DUF262 domain-containing protein | | | |
| group_13663 | phage integrase | 3 | F:GO:0003677; P:GO:0006310; P:GO:0015074 | F:DNA binding; P:DNA recombination; P:DNA integration |
| group_13664 | conjugal transfer protein TraY | 1 | C:GO:0016021 | C:integral component of membrane |
| group_13665 | hypothetical protein | | | |
| group_13666 | integrase | 3 | F:GO:0003677; P:GO:0006310; P:GO:0015074 | F:DNA binding; P:DNA recombination; P:DNA integration |
| group_13668 | putative lipoprotein | | | |
| group_1807 | transposase | 3 | F:GO:0003677; F:GO:0004803; P:GO:0006313 | F:DNA binding; F:transposase activity; P:transposition, DNA-mediated |
| group_7419 | hypothetical protein RLJV_24115 | | | |
| group_13671 | Membrane protein involved in colicin uptake-like protein | | | |
| group_13672 | Uncharacterised protein | | | |
| group_13674 | integrase | 3 | F:GO:0003677; P:GO:0006310; P:GO:0015074 | F:DNA binding; P:DNA recombination; P:DNA integration |
| yhcG | DUF1016 domain-containing protein | | | |
| group_13676 | DNA repair protein RadC | 3 | P:GO:0006508; F:GO:0008237; F:GO:0046872 | P:proteolysis; F:metallopeptidase activity; F:metal ion binding |
| group_13677 | 2-keto-D-gluconate dehydrogenase | | | |
| group_13678 | restriction endonuclease | 4 | F:GO:0004519; F:GO:0005524; F:GO:0016887; P:GO:0090305 | uclease activity; F:ATP binding; F:ATPase activity; P:nucleic acid phosphodiester bond hydrolysis |
| group_13679 | putative metal-dependent hydrolase | 1 | F:GO:0016787 | F:hydrolase activity |
| group_13680 | type I restriction endonuclease subunit R | 5 | F:GO:0003677; F:GO:0005524; F:GO:0009035; P:GO:0009307; P:GO:0090305 | e I site-specific deoxyribonuclease activity; P:DNA restriction-modification system; P:nucleic acid phosphodiester bond hydrolysis |
| group_13681 | restriction endonuclease subunit S | 4 | F:GO:0003677; F:GO:0004519; P:GO:0006304; P:GO:0090305 | nding; F:endonuclease activity; P:DNA modification; P:nucleic acid phosphodiester bond hydrolysis |
| group_13682 | SAM-dependent DNA methyltransferase | 6 | 0003677; F:GO:0004519; F:GO:0004636; P:GO:0006306; F:GO:0008170; P:GO:00 | phoribosyl-ATP diphosphatase activity; P:DNA methylation; F:N-methyltransferase activity; P:nucleic acid phosphodiester bond hydrolysis |
| group_13683 | DNA protecting protein DprA | 1 | P:GO:0009294 | P:DNA mediated transformation |
| rhlB_2 | RecQ family ATP-dependent DNA helicase | 6 | 0003676; F:GO:0003678; F:GO:0005524; P:GO:0006310; P:GO:0009116; P:GO:00 | A helicase activity; F:ATP binding; P:DNA recombination; P:nucleoside metabolic process; P:DNA duplex unwinding |
| group_13685 | WYL domain-containing protein | 1 | F:GO:0003677 | F:DNA binding |
| group_13687 | hypothetical protein HMPREF1224_11165 | | | |
| group_13688 | ---NA--- | | | |
| group_13689 | hypothetical protein | | | |
| group_13690 | ATPase involved in DNA repair | | | |
| group_13691 | hypothetical protein | | | |
| group_13692 | hypothetical protein | | | |
| group_13693 | TonB-dependent siderophore receptor | | | |
| group_13694 | hypothetical protein | | | |
| group_13695 | Uncharacterised protein | | | |
| group_13696 | ead/Ea22-like family protein | | | |
| group_13697 | Uncharacterised protein | | | |
| group_13698 | hypothetical protein | | | |
| group_13699 | phage protein | 3 | F:GO:0004725; F:GO:0008138; P:GO:0035335 | sphatase activity; F:protein tyrosine/serine/threonine phosphatase activity; P:peptidyl-tyrosine dephosphorylation |
| group_13700 | putative membrane protein | 1 | C:GO:0016021 | C:integral component of membrane |
| group_13701 | restriction alleviation protein, Lar family | | | |
| group_13702 | hypothetical protein K652_25336 | | | |
| group_13703 | hypothetical protein | 1 | F:GO:0046872 | F:metal ion binding |
| group_13704 | hypothetical protein | | | |
| group_13705 | MULTISPECIES: hypothetical protein | | | |
| group_13706 | Uncharacterised protein | | | |
| group_13707 | hypothetical protein | | | |
| group_13708 | hypothetical protein | | | |
| group_13709 | siphovirus Gp157 family protein | | | |
| group_13710 | Uncharacterised protein | | | |
| group_13711 | putative membrane protein | 1 | C:GO:0016021 | C:integral component of membrane |
| group_13712 | putative membrane protein | 1 | C:GO:0016021 | C:integral component of membrane |
| group_13713 | hypothetical protein | | | |
| group_13714 | hypothetical protein | | | |
| group_13715 | Uncharacterised protein | | | |
| group_13716 | hypothetical protein | | | |
| group_13717 | hypothetical protein | | | |
| group_13718 | Uncharacterised protein | | | |
| group_7360 | Uncharacterised protein | | | |
| group_7362 | Uncharacterised protein | | | |
| group_13719 | zinc ribbon domain-containing protein | 1 | C:GO:0016021 | C:integral component of membrane |
| group_13720 | hypothetical protein | | | |
| group_13721 | NAD-dependent DNA ligase | 1 | F:GO:0016874 | F:ligase activity |
| group_13722 | peptidase | 4 | F:GO:0003677; P:GO:0006508; F:GO:0008236; C:GO:0016021 | NA binding; P:proteolysis; F:serine-type peptidase activity; C:integral component of membrane |
| group_13723 | Cro/Cl family transcriptional regulator | 2 | F:GO:0003677; P:GO:0006355 | F:DNA binding; P:regulation of transcription, DNA-templated |
| group_13724 | hypothetical protein Q026_03401 | | | |
| group_13725 | prophage PssSM-01 | | | |
| group_13726 | phage replication protein | 1 | P:GO:0006260 | P:DNA replication |
| dnaC | replicative DNA helicase | 6 | 0003677; F:GO:0003678; F:GO:0005524; P:GO:0006269; P:GO:0032508; C:GO:19 | activity; F:ATP binding; P:DNA replication, synthesis of RNA primer; P:DNA duplex unwinding; C:primosome complex |
| group_13728 | Uncharacterised protein | | | |
| group_13729 | hypothetical protein | | | |
| group_13730 | conjugal transfer protein TraR | 1 | F:GO:0008270 | F:zinc ion binding |
| group_13731 | MULTISPECIES: hypothetical protein | | | |
| group_13732 | Uncharacterised protein | | | |
| group_13733 | DUF1367 family protein | | | |
| group_13734 | hypothetical protein | | | |
| group_13735 | recombination protein NinG | | | |
| group_13736 | 5-methyltetrahydropteroyltriglutamate--homocysteine S-methyltransferase | 1 | C:GO:0016021 | C:integral component of membrane |
| group_13737 | putative sensor/response regulator hybrid | | | |
| group_13738 | ---NA--- | | | |
| group_13739 | ---NA--- | | | |
| group_13740 | ---NA--- | | | |
| group_13741 | hypothetical protein | 1 | C:GO:0016021 | C:integral component of membrane |
| group_13742 | Uncharacterised protein | | | |
| group_13743 | terminase small subunit | | | |
| group_13744 | terminase | | | |
| group_13745 | putative membrane protein | 1 | C:GO:0016021 | C:integral component of membrane |
| group_13746 | phage portal protein | | | |
| clpP_4 | peptidase | 2 | F:GO:0004252; P:GO:0006508 | F:serine-type endopeptidase activity; P:proteolysis |
| group_13748 | phage major capsid protein | | | |
| group_13749 | Uncharacterised protein | | | |
| group_13750 | phage gp6-like head-tail connector protein | | | |
| group_13751 | prophage antirepressor | | | |
| group_13752 | RyR domain | | | |
| group_13753 | glycosyltransferase family 9 protein | 1 | F:GO:0016740 | F:transferase activity |
| group_13754 | Uncharacterised protein | | | |
| group_13755 | HK97 gp10 family phage protein | | | |
| group_13756 | phage tail protein | | | |
| group_13757 | hypothetical protein | | | |
| group_13758 | phage tail tape measure protein | 1 | C:GO:0016021 | C:integral component of membrane |
| group_13759 | Uncharacterised protein | | | |
| group_13760 | phage minor tail protein L | | | |
| group_13761 | hydrolase Nlp/P60 | | | |
| group_13762 | tail assembly protein | 1 | C:GO:0016021 | C:integral component of membrane |
| group_13763 | host specificity protein | | | |
| group_7368 | Uncharacterised protein | | | |
| group_5259 | tail fiber protein | | | |
| group_2710 | glycoside hydrolase family 19 | | | |
| group_7370 | hypothetical protein | | | |
| group_7372 | hypothetical protein | | | |
| group_5256 | MULTISPECIES: hypothetical protein | | | |
| group_27 | IS3 family transposase | | | |
| group_13764 | MULTISPECIES: hypothetical protein | 1 | C:GO:0016021 | C:integral component of membrane |
| group_13765 | integrase | 3 | F:GO:0003677; P:GO:0006310; P:GO:0015074 | F:DNA binding; P:DNA recombination; P:DNA integration |
| group_1421 | transposase | 3 | F:GO:0003677; F:GO:0004803; P:GO:0006313 | F:DNA binding; F:transposase activity; P:transposition, DNA-mediated |
| group_1419 | IS3 family transposase | | | |
| group_13766 | MULTISPECIES: hypothetical protein | | | |
| phoA2 | protein disulfide reductase | | | |
| group_13768 | hemagglutinin | | | |
| group_29 | IS3 family transposase | 1 | P:GO:0032196 | P:transposition |
| group_193 | amino acid adenylation domain-containing protein | | | |
| pupA | Ferric-pseudobactin 358 receptor precursor | 4 | C:GO:0009279; F:GO:0015344; F:GO:0038023; P:GO:0044718 | erophore uptake transmembrane transporter activity; F:signaling receptor activity; P:siderophore transmembrane transport |
| group_13770 | cyclic peptide transporter | 7 | 4; P:GO:0015833; C:GO:0016021; F:GO:0016887; F:GO:0042626; P:GO:0055085; | lmbrane; F:ATPase activity; F:ATPase-coupled transmembrane transporter activity; P:transmembrane transport; F:peptide transmembrane transporter activity |
| group_13771 | MULTISPECIES: hypothetical protein | | | |
| group_13772 | Uncharacterised protein | | | |
| group_4172 | HEAT repeat domain-containing protein | | | |
| group_13777 | Uncharacterised protein | | | |
| group_13778 | MULTISPECIES: hypothetical protein | | | |
| group_13779 | putative replication-associated protein G2P | 1 | P:GO:0006260 | P:DNA replication |
| group_13780 | hypothetical protein | | | |
| group_13781 | hypothetical protein | | | |
| group_13782 | hypothetical protein | 1 | C:GO:0016021 | C:integral component of membrane |
| group_7429 | membrane protein | 1 | C:GO:0016021 | C:integral component of membrane |
| group_13783 | coat protein A of bacteriophage Pf1 | | | |
| group_7430 | DUF2523 domain-containing protein | 1 | C:GO:0016021 | C:integral component of membrane |
| group_13784 | membrane protein | 1 | C:GO:0016021 | C:integral component of membrane |
| group_28 | IS3 family transposase | 3 | F:GO:0003676; P:GO:0015074; P:GO:0032196 | F:nucleic acid binding; P:DNA integration; P:transposition |
| group_13785 | hypothetical protein L346_02215 | | | |
| group_13786 | hypothetical protein | | | |
| group_13787 | Uncharacterised protein | 1 | C:GO:0016021 | C:integral component of membrane |
| group_13788 | hypothetical protein PAERUG_P8_2_London_17_VIM_2_07_13_05531 | | | |
| group_13789 | Uncharacterised protein | | | |
| group_13790 | Uncharacterised protein | | | |
| group_13791 | hypothetical protein PAERUG_P8_2_London_17_VIM_2_07_13_05534 | | | |
| group_13792 | putative non-ribosomal peptide synthetase | | | |
| group_13793 | probable non-ribosomal peptide synthetase | | | |

| | | | | |
|---|---|---|---|---|
| viuB | siderophore-interacting protein | 2 | F:GO:0016491; P:GO:0055114 | F:oxidoreductase activity; P:oxidation-reduction process |
| group_13794 | putative hemagglutinin | | | |
| group_261 | filamentous hemagglutinin | | | |
| group_16808 | ---NA--- | | | |
| group_13798 | putative non-ribosomal peptide synthetase | | | |
| group_13799 | hemagglutinin | | | |
| group_262 | hemagglutinin | | | |
| group_7420 | hypothetical protein RLJV_24115 | | | |
| group_6289 | phenazine-specific anthranilate synthase component I | 1 | P:GO:0006541 | P:glutamine metabolic process |
| group_6286 | trans-2,3-dihydro-3-hydroxyanthranilate isomerase | 2 | P:GO:0002047; F:GO:0102943 | phenazine biosynthetic process; F:trans-2,3-dihydro-3-hydroxy-anthranilate isomerase activity |
| group_13800 | hypothetical protein LT18_06554 | | | |
| lgrD_2 | non-ribosomal peptide synthetase | | | |
| group_13801 | putative non-ribosomal peptide synthetase | | | |
| group_13802 | ---NA--- | | | |
| dltA_4 | non-ribosomal peptide synthetase | | | |
| group_13803 | ---NA--- | | | |
| group_6295 | phospho-2-dehydro-3-deoxyheptonate aldolase | | | |
| cheW_3 | chemotaxis protein CheW | 2 | P:GO:0006935; P:GO:0007165 | P:chemotaxis; P:signal transduction |
| group_14990 | Uncharacterised protein | | | |
| group_14991 | ---NA--- | | | |
| group_14992 | Uncharacterised protein | | | |
| group_14993 | 3-oxoacyl-ACP synthase | 1 | F:GO:0004315 | F:3-oxoacyl-[acyl-carrier-protein] synthase activity |
| group_14994 | DUF4150 domain-containing protein | | | |
| group_14995 | Uncharacterised protein | | | |
| group_15003 | hypothetical protein Q080_05192 | | | |
| group_5260 | phage tail protein | | | |
| group_15004 | phage protein | | | |
| group_15006 | HDIG domain-containing protein | | | |
| group_15007 | MULTISPECIES: hypothetical protein | | | |
| group_15008 | structural protein 2 | | | |
| group_15009 | hypothetical protein | | | |
| group_15010 | hypothetical protein | | | |
| group_15011 | peptidase U35, phage prohead HK97 | 2 | P:GO:0006508; F:GO:0008233 | P:proteolysis; F:peptidase activity |
| group_15012 | phage major capsid protein | | | |
| group_15013 | DUF3077 domain-containing protein | | | |
| group_15014 | Uncharacterised protein | | | |
| group_15015 | Uncharacterised protein | | | |
| group_15016 | Uncharacterised protein | | | |
| group_15017 | AlpA family transcriptional regulator | | | |
| group_1843 | RHS repeat protein | | | |
| group_1846 | sugar-binding protein | | | |
| group_15020 | right-handed parallel beta-helix repeat-containing protein | | | |
| group_1417 | transposase | 3 | F:GO:0003677; F:GO:0004803; P:GO:0006313 | F:DNA binding; F:transposase activity; P:transposition, DNA-mediated |
| group_1415 | IS3 family transposase | | | |
| group_15021 | hypothetical protein | | | |
| group_15022 | hypothetical protein | | | |
| group_15023 | DNA translocase FtsK | 5 | F:GO:0003677; F:GO:0005524; P:GO:0006457; F:GO:0051082; P:GO:0051301 | )NA binding; F:ATP binding; P:protein folding; F:unfolded protein binding; P:cell division |
| group_15024 | DUF262 domain-containing protein | | | |
| group_15025 | SIR2 family protein | 1 | C:GO:0016021 | C:integral component of membrane |
| group_15026 | ATP-binding protein | 1 | F:GO:0005524 | F:ATP binding |
| group_22 | transposase | 3 | F:GO:0003677; F:GO:0004803; P:GO:0006313 | F:DNA binding; F:transposase activity; P:transposition, DNA-mediated |
| group_21 | IS3 family transposase | | | |
| group_15027 | ShlB/FhaC/HecB family hemolysin secretion/activation protein | | | |
| group_15028 | hemagglutination protein | | | |
| group_15029 | integrase | 3 | F:GO:0003677; P:GO:0006310; P:GO:0015074 | F:DNA binding; P:DNA recombination; P:DNA integration |
| group_15030 | DUF4224 domain-containing protein | | | |
| group_15031 | HNH endonuclease | 2 | F:GO:0004519; P:GO:0090305 | F:endonuclease activity; P:nucleic acid phosphodiester bond hydrolysis |
| group_15032 | hypothetical protein | | | |
| group_15033 | hypothetical protein OB07_02450 | | | |
| group_15034 | putative upf86.8 | | | |
| group_15035 | hypothetical protein OB07_02454 | 2 | C:GO:0016020; C:GO:0016021 | C:membrane; C:integral component of membrane |
| group_15036 | hypothetical protein OB07_02456 | 2 | C:GO:0016020; C:GO:0016021 | C:membrane; C:integral component of membrane |
| group_15037 | hypothetical protein | | | |
| recT | recombinase RecT | 2 | F:GO:0003677; P:GO:0006259 | F:DNA binding; P:DNA metabolic process |
| group_15039 | endonuclease | 2 | F:GO:0004519; P:GO:0090305 | F:endonuclease activity; P:nucleic acid phosphodiester bond hydrolysis |
| group_15040 | hypothetical protein | | | |
| group_15041 | hypothetical protein Q061_01735 | | | |
| group_15042 | MULTISPECIES: hypothetical protein | | | |
| group_15043 | regulatory protein | | | |
| group_15044 | nucleotide pyrophosphohydrolase | 3 | P:GO:0009143; F:GO:0046872; F:GO:0047429 | : triphosphate catabolic process; F:metal ion binding; F:nucleoside-triphosphate diphosphatase activity |
| group_15045 | HNH endonuclease | 1 | F:GO:0003677 | F:DNA binding |
| group_15046 | HNH endonuclease | 3 | F:GO:0003676; F:GO:0004519; P:GO:0090305 | nucleic acid binding; F:endonuclease activity; P:nucleic acid phosphodiester bond hydrolysis |
| group_15047 | Uncharacterised protein | | | |
| group_15048 | XRE family transcriptional regulator | 2 | F:GO:0003677; F:GO:0016787 | F:DNA binding; F:hydrolase activity |
| group_15049 | HicB family protein | | | |
| group_15050 | phage holin, lambda family | 1 | C:GO:0016021 | C:integral component of membrane |
| group_15051 | glycoside hydrolase family 19 | | | |
| group_15052 | phage lysis regulatory protein, LysB family | | | |
| group_15053 | hypothetical protein | | | |
| group_15054 | DNA-packaging protein | | | |
| group_15055 | terminase | | | |
| group_15056 | preprotein translocase subunit SecA | 1 | P:GO:0019058 | P:viral life cycle |
| group_15057 | Clp protease ClpP | | | |
| group_15058 | head decoration protein | | | |
| group_15059 | major capsid protein E | | | |
| group_15060 | Uncharacterised protein | | | |
| group_15061 | MULTISPECIES: hypothetical protein | | | |
| group_15062 | MULTISPECIES: hypothetical protein | | | |
| group_15063 | MULTISPECIES: hypothetical protein | | | |
| group_15064 | RICIN domain-containing protein | | | |
| group_15065 | tape measure domain-containing protein | | | |
| group_15066 | MULTISPECIES: hypothetical protein | | | |
| group_15067 | hypothetical protein | | | |
| group_15068 | MULTISPECIES: hypothetical protein | | | |
| group_15069 | hypothetical protein | | | |
| group_15070 | Uncharacterised protein | | | |
| group_15071 | MULTISPECIES: hypothetical protein | | | |
| group_1848 | sugar-binding protein | | | |
| group_15072 | Uncharacterised protein | | | |
| group_192 | non-ribosomal peptide synthetase | 2 | F:GO:0003824; F:GO:0031177 | F:catalytic activity; F:phosphopantetheine binding |
| group_15073 | MULTISPECIES: hypothetical protein | | | |
| group_15074 | transcriptional regulator | 1 | F:GO:0003677 | F:DNA binding |
| group_15075 | phage protein | 1 | P:GO:0006260 | P:DNA replication |
| group_15076 | attachment protein | | | |
| group_15082 | DUF4747 domain-containing protein | | | |
| group_15085 | DUF262 domain-containing protein | | | |
| group_15086 | SIR2 family protein | | | |
| group_15087 | ParA family protein | | | |
| group_15088 | hypothetical protein Q092_03300 | | | |
| group_15089 | integrase | 3 | F:GO:0003677; P:GO:0006310; P:GO:0015074 | F:DNA binding; P:DNA recombination; P:DNA integration |
| group_2228 | Rhs element Vgr protein | | | |
| group_15092 | aldolase | 1 | F:GO:0008701 | F:4-hydroxy-2-oxovalerate aldolase activity |
| group_15093 | methyltransferase type 12 | 2 | F:GO:0008168; P:GO:0032259 | F:methyltransferase activity; P:methylation |
| group_15094 | methyltransferase, FkbM family protein | | | |
| kpsU | 3-deoxy-manno-octulosonate cytidylyltransferase | 1 | F:GO:0008690 | F:3-deoxy-manno-octulosonate cytidylyltransferase activity |
| group_15096 | MULTISPECIES: hypothetical protein | | | |
| group_15100 | hemophilus-specific protein, uncharacterized | | | |
| group_15101 | hypothetical protein | | | |
| group_15102 | Uncharacterised protein | | | |
| cas2 | CRISPR-associated endonuclease Cas2 | 5 | F:GO:0004521; P:GO:0043571; F:GO:0046872; P:GO:0051607; P:GO:0090502 | 'f CRISPR repeat elements; F:metal ion binding; P:defense response to virus; P:RNA phosphodiester bond hydrolysis, endonucleolytic |
| cas4-cas1 | subtype I-C CRISPR-associated endonuclease Cas1 | 6 | :0003677; F:GO:0004520; P:GO:0043571; F:GO:0046872; P:GO:0051607; P:GO:00 | P:maintenance of CRISPR repeat elements; F:metal ion binding; P:defense response to virus; P:nucleic acid phosphodiester bond hydrolysis |
| group_15105 | CRISPR-associated protein Cas4 | 5 | F:GO:0004527; F:GO:0046872; F:GO:0051536; P:GO:0051607; P:GO:0090305 | al ion binding; F:iron-sulfur cluster binding; P:defense response to virus; P:nucleic acid phosphodiester bond hydrolysis |
| group_15106 | type I-C CRISPR-associated protein Cas7/Csd2 | 1 | P:GO:0043571 | P:maintenance of CRISPR repeat elements |
| group_15107 | type I-C CRISPR-associated protein Cas8c/Csd1 | | | |
| cas5d | type I-C CRISPR-associated protein Cas5 | 4 | F:GO:0004519; P:GO:0043571; P:GO:0051607; P:GO:0090305 | maintenance of CRISPR repeat elements; P:defense response to virus; P:nucleic acid phosphodiester bond hydrolysis |
| group_15109 | CRISPR-associated helicase/endonuclease Cas3 | 4 | F:GO:0003676; F:GO:0004519; F:GO:0005524; P:GO:0090305 | cid binding; F:endonuclease activity; F:ATP binding; P:nucleic acid phosphodiester bond hydrolysis |
| group_15111 | hypothetical protein OB07_04185 | 1 | C:GO:0016021 | C:integral component of membrane |
| group_15113 | Thermostable hemolysin | | | |
| group_15114 | cytochrome C | | | |
| shp | DUF1924 domain-containing protein | 4 | F:GO:0009055; F:GO:0020037; P:GO:0022900; F:GO:0046872 | ':electron transfer activity; F:heme binding; P:electron transport chain; F:metal ion binding |
| hmp_3 | 2Fe-2S iron-sulfur cluster binding domain-containing protein | 5 | F:GO:0009055; C:GO:0016021; P:GO:0022904; F:GO:0046872; F:GO:0051537 | egral component of membrane; P:respiratory electron transport chain; F:metal ion binding; F:2 iron, 2 sulfur cluster binding |
| group_1418 | transposase | 3 | F:GO:0003677; F:GO:0004803; P:GO:0006313 | F:DNA binding; F:transposase activity; P:transposition, DNA-mediated |
| group_1416 | IS3 family transposase | | | |
| group_15117 | periplasmic protein | 1 | F:GO:0016872 | F:intramolecular lyase activity |
| group_15118 | metal ABC transporter ATPase | | | |
| group_15119 | DUF2878 domain-containing protein | 1 | C:GO:0016021 | C:integral component of membrane |
| group_15120 | DUF1365 domain-containing protein | | | |
| group_15121 | short-chain dehydrogenase | 2 | F:GO:0016491; P:GO:0055114 | F:oxidoreductase activity; P:oxidation-reduction process |
| group_15122 | transcriptional regulator | | | |
| group_15123 | MerR family transcriptional regulator | 2 | F:GO:0003677; P:GO:0006355 | F:DNA binding; P:regulation of transcription, DNA-templated |
| group_15124 | DNA helicase UvrD | 4 | F:GO:0003677; F:GO:0003678; F:GO:0005524; P:GO:0032508 | F:DNA binding; F:DNA helicase activity; F:ATP binding; P:DNA duplex unwinding |
| group_15125 | ATP-dependent endonuclease | 2 | F:GO:0004519; P:GO:0090305 | F:endonuclease activity; P:nucleic acid phosphodiester bond hydrolysis |
| group_15128 | ---NA--- | | | |
| group_15129 | ---NA--- | | | |
| group_15130 | tRNA-dependent cyclodipeptide synthase | 1 | F:GO:0016755 | F:transferase activity, transferring amino-acyl groups |
| group_15131 | isopenicillin N synthase family oxygenase | 3 | F:GO:0016491; F:GO:0046872; P:GO:0055114 | F:oxidoreductase activity; F:metal ion binding; P:oxidation-reduction process |
| group_15132 | 2OG-Fe(II) oxygenase | 3 | F:GO:0016491; F:GO:0046872; P:GO:0055114 | F:oxidoreductase activity; F:metal ion binding; P:oxidation-reduction process |
| group_15133 | cytochrome | 4 | F:GO:0005506; F:GO:0016705; F:GO:0020037; P:GO:0055114 | ivity, acting on paired donors, with incorporation or reduction of molecular oxygen; F:heme binding; P:oxidation-reduction process |
| group_15134 | isopenicillin N synthase family oxygenase | 2 | F:GO:0016491; P:GO:0055114 | F:oxidoreductase activity; P:oxidation-reduction process |
| group_15135 | 2OG-Fe(II) oxygenase superfamily protein | 3 | F:GO:0016491; F:GO:0046872; P:GO:0055114 | F:oxidoreductase activity; F:metal ion binding; P:oxidation-reduction process |
| group_15136 | isopenicillin N synthase family oxygenase | 3 | F:GO:0046872; F:GO:0051213; P:GO:0055114 | F:metal ion binding; F:dioxygenase activity; P:oxidation-reduction process |
